# Context-dependent utility and robustness of pretrained single-cell foundation model representations across analytical tasks

**DOI:** 10.64898/2026.06.18.733285

**Authors:** Tianyuan Liu, Tingze Feng, Xianrun Pan, Yangyang Chen, Liping Ren, Xiucai Ye, Hao Lin, Yang Zhang

**Author notes:** Corresponding author. (Yang Zhang); (Hao Lin); (Xiucai Ye). These authors contributed equally to this work (Tianyuan Liu, Tingze Feng).

## Abstract

Single-cell foundation models (scFMs) have emerged as powerful representation learning approaches for single-cell transcriptomics. However, the utility and robustness of their pretrained representations across diverse analytical tasks and data conditions remain insufficiently characterized, particularly in zero-shot settings without task-specific fine-tuning. Here, we systematically analyze zero-shot performance of single-cell transcriptomic representations across 20 methods, 6 downstream tasks and 1,607 datasets comprising nearly 21.8 million cells. We evaluate model behavior along three complementary dimensions: utility on original datasets, robustness to controlled changes in dataset structure, and exploratory associations between dataset characteristics and performance variation. Our results show that scFM performance is strongly task dependent, with no single method consistently outperforming others across cell- and gene-level analyses. Notably, high utility on original datasets did not necessarily translate into robustness under structural perturbations, and several top-ranking methods were sensitive to changes in cell number, gene number, class composition, class imbalance, and batch complexity. Conventional statistical and task-specific methods remained competitive in several settings, while greater computational cost did not consistently correspond to better performance. Driver analyses further identified task-specific associations between performance and dataset characteristics, including cell-type complexity, train-test class overlap, batch number, and regulatory target-set size. Together, these findings show that the zero-shot utility and robustness of pretrained scFM representations depend jointly on analytical task and dataset structure. Our study provides a practical basis for context-aware representation selection and underscores the importance of evaluating structural robustness alongside utility when developing and applying scFMs.

## INTRODUCTION

Single-cell transcriptomics is now generating data at unprecedented scale (*1, 2*), enabling detailed characterization of cellular heterogeneity across tissues (*3*), developmental stages (*4*) and disease contexts (*5*). As single-cell RNA-sequencing (scRNA-seq) atlases expand to hundreds of millions of cells, this growth has driven intense interest in large-scale representation learning for biology (*6*). Inspired by advances in foundation models in natural language processing (*7–9*), several single-cell foundation models (scFMs) have been developed to learn generalizable embeddings from large-scale pretraining (*9–20*). In particular, models such as scFoundation (*10*), UCE (*18*) and GenePT (*15*) have explicitly emphasized zero-shot applicability, proposing that pretrained representations can be reused across downstream tasks without task-specific retraining. If this premise holds broadly, representation learning could shift from task-centered model construction toward reusable analytical backbones for single-cell data.

Despite these expectations, the extent to which scFMs have been incorporated into routine analytical workflows remains uncertain. Our citation-context analysis of 1,330 studies citing original scFM publications suggests that direct downstream analytical adoption remains limited (Supplementary Note 1, Supplementary Fig. 1). While these models are widely referenced in the literature, their routine incorporation into practical analytical pipelines appears less common than anticipated. In practice, extensive fine-tuning demands substantial computational resources, mature software infrastructures and systematic hyperparameter optimization, conditions that are not universally available. Under such constraints, zero-shot deployment therefore becomes a fundamental prerequisite for practical adoption rather than an optional convenience. Researchers commonly extract pretrained embeddings directly for downstream analysis, particularly when processing large-scale atlases. Therefore, the critical question is whether zero-shot representations alone provide stable and reproducible advantages over simpler statistical baselines such as principal component analysis (PCA), which remain widely adopted because of their interpretability and computational efficiency.

To support the rapid development of this field, several benchmarking efforts have mapped an initial landscape of scFM capabilities across three complementary tracks (*21–26*). Technical benchmarks, such as scEval (*25*), focus on model optimization and training stability to provide guidelines for fine-tuning. Conversely, biological application-centric benchmarks, including BioLLM (*23*) and Wu et al. (*21*), investigate model performance in complex biomedical scenarios to guide model selection for specific biological tasks. Concurrently, critical evaluations targeting specific tasks, such as cell annotation (*24*) or batch integration (*22*), aim to investigate whether massive pretrained architectures can outperform simpler baselines. A general consensus from these diverse studies is that no single scFM consistently dominates. While these benchmark studies establish an essential landscape, three critical methodological gaps remain in evaluating practical zero-shot applicability. First, current benchmarks rely almost exclusively on widely adopted datasets, leaving the vulnerability of pretrained embeddings to structural data shifts, such as changes in cell or gene numbers, largely untested. Second, zero-shot cross-task assessments lack the algorithmic breadth and dataset scale required for comprehensive validation. Third, the quantitative relationship between zero-shot efficacy and specific dataset-level characteristics remains unexplored. Resolving these uncertainties is essential for single-cell analyses in translational research (*27–29*), as overlooking representation stability under realistic variability risks overestimating generalization claims while neglecting simpler, highly robust alternatives.

Here we define the practical limits of zero-shot generalization in scFMs through a structured evaluation framework spanning both cell-level and gene-level analyses. We evaluated 20 representation methods across 6 downstream tasks on 1,607 datasets encompassing nearly 21.8 million cells. These include original datasets alongside their structural perturbations. Specifically, we characterize model performance along three complementary dimensions: (i) baseline utility on original datasets, (ii) robustness on structurally perturbed datasets and (iii) exploratory associations between dataset-level features and performance. By employing structured data augmentation to alter dataset properties, such as cell and gene count, we provide a principled methodology to evaluate representation stability under controlled data shifts. Rather than focusing solely on average task-level rankings, this framework enables direct characterization of the practical zero-shot utility boundaries of scFMs. It elucidates the critical relationship between baseline utility and structural robustness, quantitatively capturing model sensitivity to dataset-level heterogeneity. Together, these analyses contribute to a more transparent basis for evaluating generalization claims and offer practical guidance for model selection in routine single-cell analysis.

## RESULTS

### Overview of the benchmarking framework

To examine how scFMs perform under routine analytical conditions, we established a structured evaluation framework that assesses pretrained representations across diverse biological tasks and perturbations of dataset structural properties (Fig. 1). Rather than focusing solely on average task-level rankings, this design measures the robustness of zero-shot representations against realistic heterogeneity in single-cell data. Pretrained models transform scRNA-seq data into biological representations, including cell embeddings, gene embeddings and gene-gene attention scores (Fig. 1a). These representations were evaluated under zero-shot conditions, without task-specific fine-tuning. We examined six downstream applications spanning unsupervised cell clustering, supervised annotation and network inference, reflecting common analytical scenarios in single-cell studies (Fig. 1b). For each task, original benchmark datasets were paired with systematically generated variants produced through perturbations of dataset structure, including cell number, gene number, cell-type number, class imbalance and batch complexity. This matched design enables direct comparison between unperturbed and structurally altered datasets. Model behavior was characterized along three complementary dimensions (Fig. 1c). Utility was defined as performance on original datasets. Robustness quantified the magnitude of performance change under dataset structural perturbations. In addition, driver analysis examined associations between dataset-level characteristics and performance variation using multivariable linear regression model. Overall, this framework provides a principled basis for delineating zero-shot utility and s robustness across heterogeneous single-cell contexts.

**Fig. 1.**
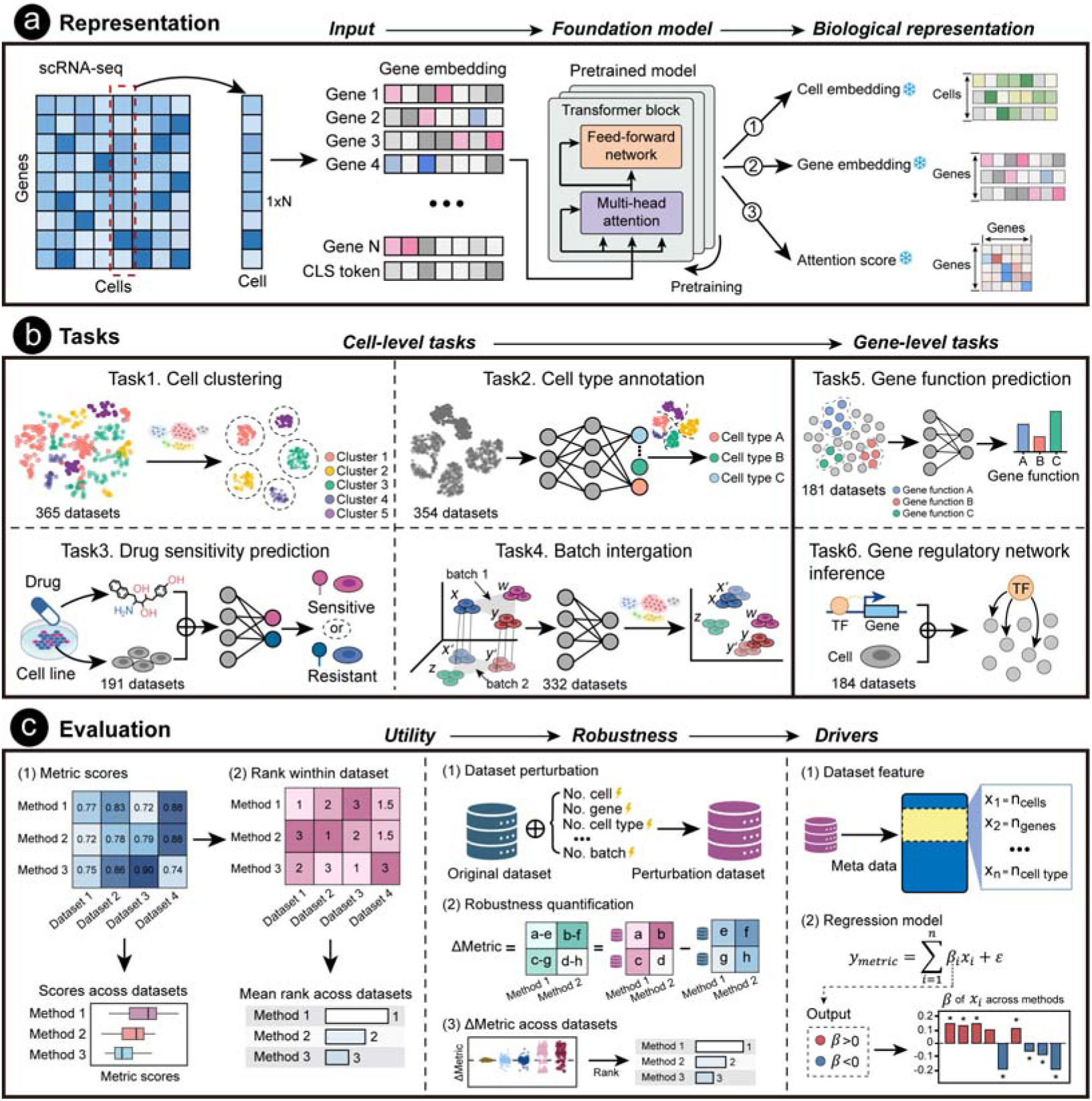
Overview of the benchmarking framework. **a**, Representation extraction. Input scRNA-seq data are processed by pre-trained foundation models to generate distinct biological representations: cell embeddings, gene embeddings, and attention scores. All representations are extracted under zero-shot conditions, without task-specific fine-tuning (denoted by snowflake icons). **b,** Downstream tasks. The benchmarking pipeline encompasses six downstream tasks, categorized into cell-level analyses (cell clustering, cell-type annotation, drug sensitivity prediction, and batch integration) and gene-level analyses (gene function prediction and gene regulatory network inference). **c,** Evaluation dimensions. Model performance is characterized across three complementary axes. Utility quantifies baseline performance on original datasets through direct metric scoring and cross-dataset rank aggregation. Robustness measures performance stability under systematic perturbations of dataset structural properties. Drivers employs regression modeling to identify how specific dataset-level meta-features dictate model efficacy.

### Cell clustering under zero-shot representations

We first evaluated zero-shot cell embeddings in the context of cell clustering across 26 original datasets. Eighteen representation methods were compared, including scFMs together with PCA and scGNN (*30*) as reference baselines (Methods). For each dataset, embeddings were extracted without task-specific retraining and clustered using a unified scIB (*31*) workflow. Performance was assessed both on original datasets (utility) and on 339 datasets generated through perturbations of dataset structural properties (robustness). Clustering accuracy was quantified using adjusted Rand Index (ARI), normalized mutual information (NMI) and average silhouette width (ASW), capturing agreement with reference cell labels and geometric separability in embedding space.

Under original dataset conditions, models exhibited clear performance stratification (Fig. 1b,d,f). Based on aggregated ranks, UCE4L, SCimilarity (*16*) and CellPLM (*17*) achieved the highest overall clustering utility (Fig. 1a, Supplementary Fig. 2-3). UCE4L and SCimilarity consistently ranked among the top performers across ARI, NMI and ASW, whereas CellPLM remained within the top tier for ARI and ASW with a modest decline in NMI. Across datasets, several scFMs significantly outperformed PCA based on paired comparisons, indicating that large-scale pretraining can improve recovery of cell-type structure under zero-shot settings. However, higher utility on original datasets did not necessarily translate into robustness under structural perturbations. Performance deviations under perturbations varied substantially across methods (Fig. 1c,e,g). Robustness rankings revealed a pattern distinct from utility rankings. CellFMmin (*14*), scBERTMean (*20*) and scBERTCLS exhibited smaller deviations across multiple metrics and ranked highly in overall robustness (Supplementary Fig. 4-5). Utility and robustness were not aligned. Methods with lower utility sometimes exhibited smaller absolute deviations, suggesting that limited dynamic range may partly constrain observed fluctuations. Robustness rankings alone therefore do not fully characterize practical performance.

To jointly examine clustering utility and robustness, models were mapped in a two-dimensional space (Fig. 1h), where lower ranks indicate better performance. Using asymmetric thresholds (top quartile for utility and median for robustness), we identified models combining high utility with acceptable robustness. In the overall space, UCE4L and CellPLM occupied the preferred region, indicating a balance between clustering utility and robustness. SCimilarity achieved strong utility but fell below the robustness threshold. Metric-specific projections further revealed heterogeneity. In ARI space, SCimilarity satisfied both thresholds, whereas in NMI space UCE4L and CellPLM entered the preferred region. In contrast, ASW rankings were reordered, with scFoundation uniquely satisfying both criteria, indicating divergence between geometric separability and label-consistency metrics.

Drivers analysis identified systematic statistical associations between clustering performance and dataset structural properties. For ARI and NMI, cell-type number and total cell count exhibited the largest average absolute standardized coefficients, indicating that clustering performance is associated with population complexity and sample scale (Supplementary Fig. 6a). For ASW, cell-type number exhibited the strongest association, whereas total cell number and gene number showed weaker effects, suggesting that geometric separability is more sensitive to class structure than dataset size alone. Across models, the direction of association was largely consistent (Supplementary Fig. 6b). Heterogeneity across models was primarily reflected in effect magnitude rather than direction. These results suggest that clustering performance under zero-shot representations is systematically related to dataset composition, although effect sizes differ across models. In addition, the time for cell representation extraction ranged from approximately 1-2 minutes to over 30 minutes, with substantial variation in CPU and GPU memory usage (Supplementary Table 2). Computational cost did not directly track clustering utility, underscoring that model selection in routine single-cell analysis may involve trade-offs among utility, robustness and resource consumption.

**Figure 2.**
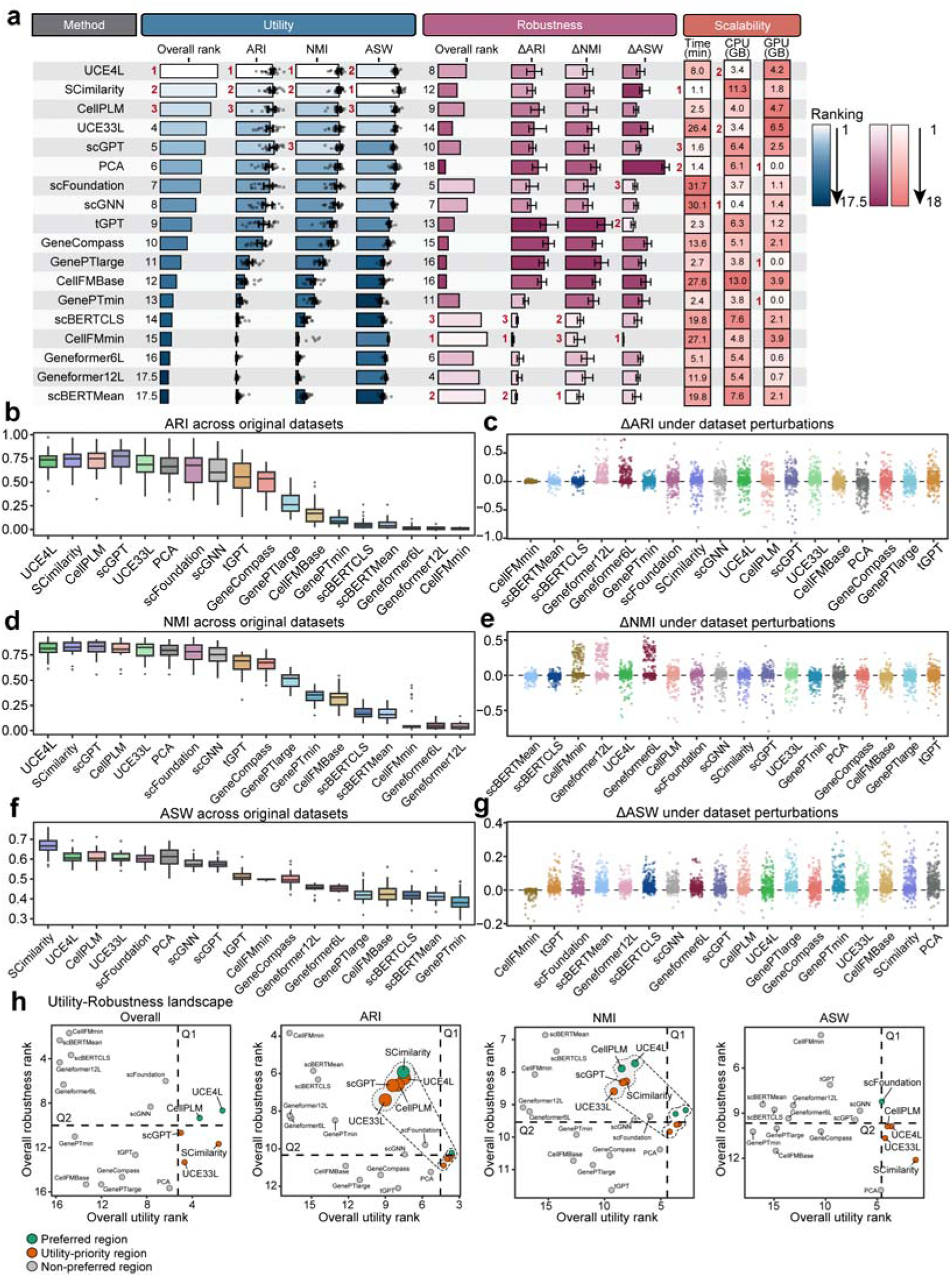
Cell clustering under zero-shot representations. **a,** Performance overview across original datasets (n=26), perturbation-derived datasets (n=339), and computational scalability metrics (runtime, CPU and GPU memory usage). Utility and robustness were summarized using rank-based aggregation across ARI, NMI and ASW. Heatmap color gradients indicate relative method rankings. **b,d,f,** Clustering performance on original datasets measured by ARI **(b)**, NMI **(d)** and ASW **(f)**. Box plots display the distribution of metric values across datasets for each of the 18 methods. **c,e,g,** Performance deviations under structural perturbations measured by ARI **(c)**, NMI **(e)**, and ASW **(g)**. Each point represents the deviation from the matched original dataset across perturbation datasets. **h,** Utility-robustness landscapes based on overall rank aggregation and individual evaluation metrics. Each point represents a single method. The x-axis indicates utility rank and the y-axis indicates robustness rank, where lower numerical values denote superior performance. Colors denote decision-oriented regions defined by asymmetric thresholds: top quartile for utility (Q1) and the median for robustness (Q2). ARI, adjusted Rand Index; NMI, normalized mutual information; ASW, average silhouette width.

### Cell type annotation under zero-shot representations

We next evaluated zero-shot representations for supervised cell type annotation across 26 original datasets. Eighteen representation methods were compared, including scFMs together with highVariable (*32*) and scGNN as reference baselines (Methods). Stratified five-fold cross-validation was applied within each dataset, and embeddings were extracted independently for training and testing partitions without sharing intermediate statistics. A unified fully connected classifier was trained on all embeddings to ensure comparability across methods. Performance was assessed on original datasets (utility) and on 328 datasets generated through perturbations of dataset structural properties (robustness). Classification accuracy was quantified using F1-score, AUROC (area under the receiver operating characteristic curve) and AUPRC (area under the precision-recall curve).

Under original dataset conditions, models exhibited clear stratification in supervised discriminative performance (Fig. 3b,d,f). Based on aggregated ranks, scFoundation achieved the highest overall utility, followed by highVariable and CellPLM (Fig. 3a, Supplementary Fig. 7-8). scFoundation ranked consistently at the top across F1-score, AUROC and AUPRC. Notably, highVariable remained competitive under the unified classifier, indicating that traditional statistical representations retain strong discriminative capacity in supervised settings. While AUROC values approached saturation for multiple models, larger separations were observed in F1-score and AUPRC. Cross-validation variance was low across folds, suggesting stable performance estimation (Supplementary Fig. 9). Under perturbations of dataset structural properties, most methods exhibited performance deviations, particularly in F1-score and AUPRC (Fig. 3c,e,g). Robustness rankings differed from utility rankings. CellFMmin, GenePTmin and GenePTlarge showed smaller performance deviations overall, whereas some methods with high utility, such as highVariable and CellPLM, displayed greater sensitivity to structural perturbations (Supplementary Fig. 10-11). As observed in clustering, higher utility did not necessarily imply greater robustness.

To evaluate the trade-off between model utility and robustness, models were mapped in the two-dimensional utility-robustness landscape (Fig. 3h). Using the same asymmetric thresholds as in clustering (top quartile for utility and median for robustness), scFoundation, UCE4L and CellPLM occupied the preferred region. highVariable and UCE33L achieved high utility but fell below the robustness threshold. Metric-specific projections showed that scFoundation maintained favorable positioning across all three metrics, whereas CellPLM and UCE4L ranked highly in utility but did not satisfy the robustness criterion under F1-score and AUPRC. These results indicate that high supervised utility does not consistently translate into robustness under structural perturbations. Furthermore, drivers analysis identified strong statistical associations between classification performance and dataset structural properties. The proportion of overlapping cell types between training and testing partitions exhibited the largest standardized coefficients across F1-score, AUROC and AUPRC, indicating that supervised performance is strongly associated with class composition consistency (Supplementary Fig. 12a). Associations with cell number, class imbalance and gene number were smaller in magnitude. Across models, the direction of association was broadly consistent, with heterogeneity primarily reflected in effect size rather than sign (Supplementary Fig. 12b,c and Supplementary Fig. 13). These results indicate that cell annotation performance under zero-shot representations depends strongly on the overlap of cell types between training and testing partitions.

**Fig. 3.**
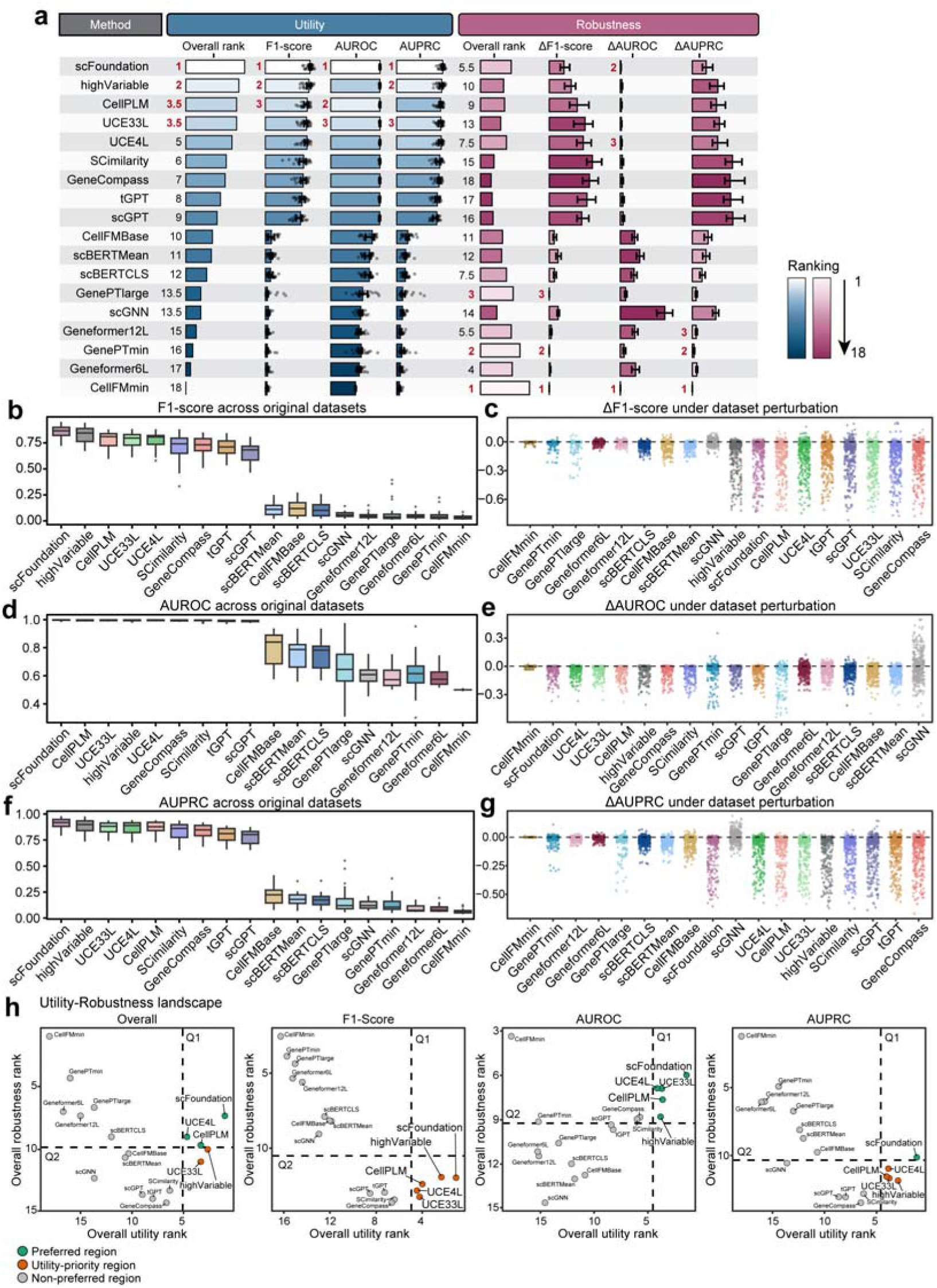
Cell type annotation under zero-shot representations. **a,** Performance overview across original datasets (n=26) and perturbation-derived datasets (n=328). Utility and robustness were summarized by rank-based aggregation of F1-score, AUROC and AUPRC. **b,d,f,** Distribution of F1-score **(b)**, AUROC **(d)** and AUPRC **(f)** across original datasets. Box plots show the distribution of metric values across datasets for each of the 18 methods. **c,e,g,** Performance deviations under structural perturbations across perturbation-derived datasets, shown as paired changes relative to corresponding original datasets for F1-score **(c)**, AUROC **(e)** and AUPRC **(g)**. Each point represents one perturbation instance. **h,** Utility-robustness landscape for 18 methods under overall ranking and individual metrics. Colors indicate predefined decision regions based on asymmetric rank thresholds: top quartile for utility (Q1) and median for robustness (Q2). AUROC, area under the receiver operating characteristic curve; AUPRC, area under the precision-recall curve.

### Drug sensitivity prediction under zero-shot representations

We next evaluated zero-shot representations in a phenotype prediction setting, focusing on drug sensitivity prediction across 14 curated single-cell datasets derived from DRMref (*33*). In contrast to clustering and annotation tasks that assess structural recovery or label discrimination, this task evaluates whether pretrained embeddings capture cell-state representations that supports drug sensitivity prediction under zero-shot conditions. For each dataset, a binary classification problem (resistant versus non-resistant) was constructed at the cell level. All methods provided cell embeddings as input features, extracted under zero-shot settings without fine-tuning. A unified dual-tower neural architecture integrated cell embeddings with drug molecular features encoded as 1,024-dimensional Morgan fingerprints derived from SMILES representations. All models were trained under identical cross-validation, optimization and architectural conditions to ensure that performance differences reflected embedding quality rather than classifier design (Methods). Performance was evaluated on original datasets (utility) and on 177 datasets generated through perturbations of dataset structural properties and domain-shift scenarios (robustness). F1-score, AUROC and AUPRC were used as complementary metrics.

Under original dataset conditions, models exhibited clear stratification in predictive performance (Fig. 4b,d,f). Based on aggregated ranks, highVariable and scFoundation achieved the highest overall utility, followed by GeneCompass (*13*) and tGPT (*19*) (Fig. 4a, Supplementary Fig. 14-15). Notably, highVariable achieved the top comprehensive ranking and performed strongly across all three metrics, indicating that statistic-based features remain competitive in drug sensitivity prediction.

scFoundation also maintained consistently high rankings across F1-score, AUROC and AUPRC under zero-shot settings, suggesting that its representations are informative for drug sensitivity prediction. Cross-validation variance was low across folds, supporting stability of performance estimation (Supplementary Fig. 16). In addition, under perturbations of dataset structural properties, methods exhibited measurable performance deviations (Fig. 4c,e,g). Robustness rankings were broadly aligned with utility rankings in this task. highVariable and scFoundation demonstrated smaller overall deviations across metrics (Supplementary Fig. 17-18). In contrast to clustering and annotation tasks, the consistency between utility and robustness was higher in drug sensitivity prediction.

To jointly assess utility and robustness, models were mapped in the two-dimensional utility-robustness landscape (Fig. 4h). Given the stronger consistency between utility and robustness observed in this task, we applied symmetric thresholds (top quartile for both dimensions) to identify models simultaneously achieving high utility and robustness. With this threshold, highVariable, scFoundation, GeneCompass and tGPT occupied the preferred region in the overall space. Furthermore, all four methods consistently remained within the preferred region across F1-score, AUROC and AUPRC subspaces. Overall, drug sensitivity prediction exhibited comparatively stronger alignment between utility and robustness than observed in previous tasks. Furthermore, drivers analysis revealed associations between predictive performance and dataset structural properties (Supplementary Fig. 19-21). Across metrics, the degree of overlap in cell-type composition between training and testing partitions exhibited the largest standardized coefficients, indicating that consistency in cellular composition strongly influences predictive performance (Supplementary Fig. 19a). Training-testing drug overlap also showed positive associations with AUROC, AUPRC and F1-score, suggesting that cross-drug structural consistency contributes to stable prediction. In contrast, total cell number and class imbalance showed comparatively smaller effects. These findings indicate that drug sensitivity prediction performance depends strongly on structural consistency across both cellular and pharmacological domains.

**Fig. 4.**
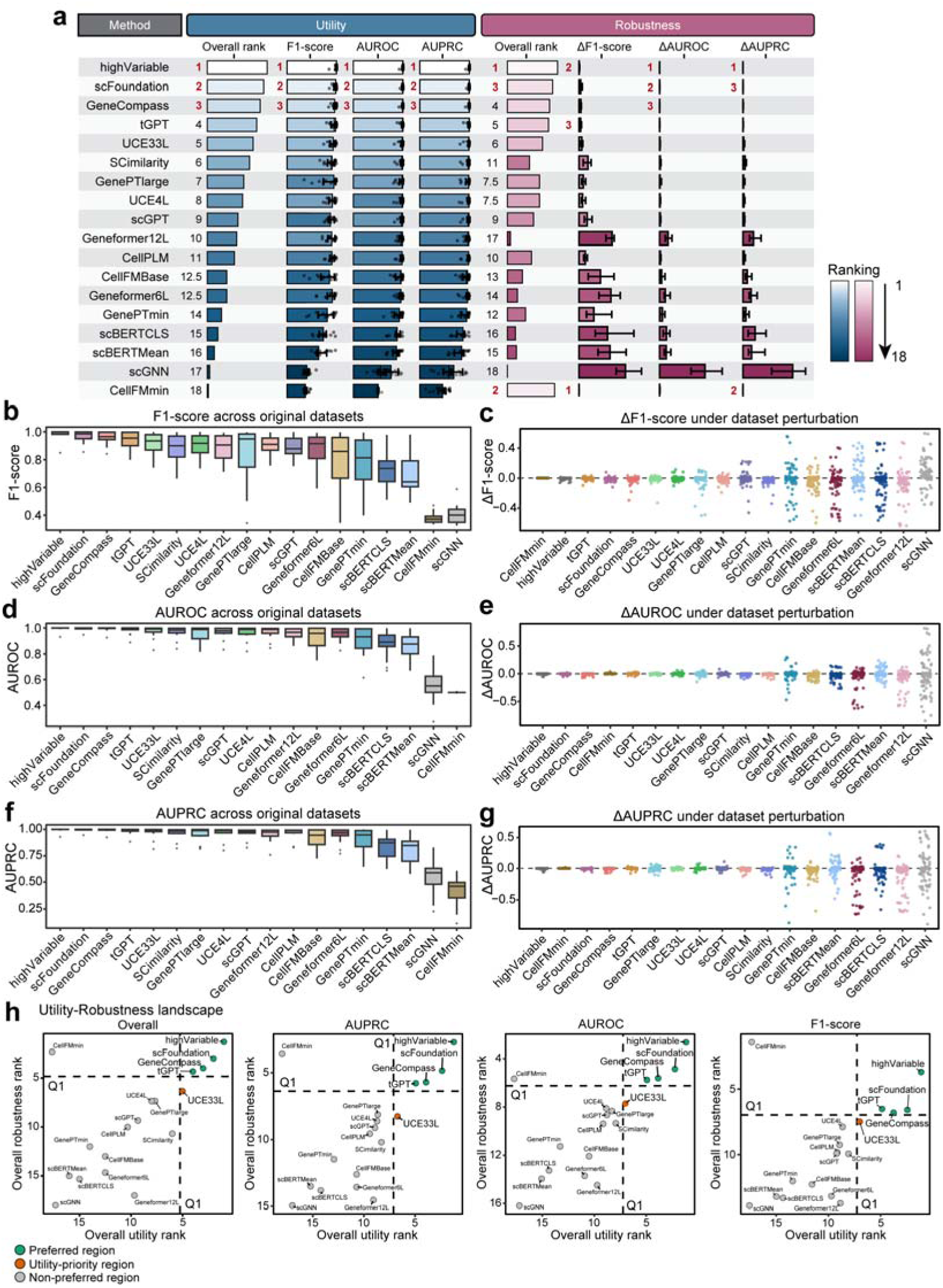
Drug sensitivity prediction under zero-shot representations. **a,** Performance overview across original datasets (n=14) and perturbation-derived datasets (n=79). Utility and robustness were summarized using rank-based aggregation across F1-score, AUROC and AUPRC. **b,d,f,** Boxplots showing performance distributions of 18 methods across original datasets measured by F1-score, AUROC and AUPRC, respectively. **c,e,g,** Scatter plots showing performance deviations under perturbation-derived datasets for F1-score, AUROC and AUPRC, respectively. Each point represents a perturbation instance. **h,** Utility-robustness landscape under overall ranking and individual metrics. Each point represents one method; lower ranks indicate better performance. Colors denote predefined decision regions based on symmetric thresholds (top quartile for both utility and robustness).

### Batch integration under zero-shot representations

We next evaluated zero-shot representations in the context of batch integration on seven widely used and representative multi-technology single-cell datasets. In contrast to supervised tasks, batch integration is an unsupervised integration problem that seeks to attenuate technical variation while preserving biological structure. Eighteen representation methods were compared, including scFMs together with Harmony (*34*) and scGNN as reference baselines. For all scFMs, embeddings were extracted under zero-shot settings without batch-specific fine-tuning and evaluated using a unified scIB workflow (Methods). Integration quality was assessed on original datasets (utility) and on 325 datasets generated through perturbations of dataset structural properties (robustness). Integration quality was quantified using complementary metrics. Biological conservation was evaluated using ARI, NMI and ASW to measure preservation of cell-type structure after integration. Batch mixing was evaluated using iLISI (inverse local inverse Simpson’s index), kBET (k-nearest-neighbor batch effect test) and batch ASW to quantify the degree of batch alignment in embedding space.

Under original dataset conditions, integration utility exhibited clear stratification across methods (Fig. 5b,d and Supplementary Fig. 22). Harmony achieved the highest overall rank, followed by scGPT (*11*) and SCimilarity (Fig. 5a and Supplementary Fig. 23-26). Harmony ranked first in ARI and NMI and remained highly competitive across batch mixing metrics, indicating consistently strong performance across both conservation and mixing objectives. scGPT ranked within the top five in five of the six evaluated metrics and achieved the highest score in kBET, demonstrating broad competitiveness across integration criteria. SCimilarity performed strongly in ARI, NMI and ASW but ranked comparatively lower in batch mixing metrics, resulting in a slightly reduced overall standing. These results indicate that while several zero-shot foundation models exhibit measurable integration capacity, explicit batch-aware integration strategies remain highly competitive in this setting. Under perturbations of dataset structural properties, robustness patterns diverged from baseline utility rankings (Fig. 5c,e and Supplementary Fig. 27). scFoundation and CellFMmin exhibited smaller performance deviations across multiple metrics and ranked highly in overall robustness (Fig. 5a and Supplementary Fig. 28-31). In contrast, Harmony and scGPT, despite strong baseline utility, showed larger fluctuations under structural perturbations, particularly in ASW and selected batch mixing metrics. As in previous tasks, higher utility did not consistently correspond to greater robustness under structural variability.

To better understand this divergence, we jointly evaluated utility and robustness. Mapping models into the two-dimensional utility-robustness space further clarified this relationship (Fig. 5f-h). Using the same asymmetric thresholds as in clustering (top quartile for utility and median for robustness), UCE4L and SCimilarity occupied the preferred region, indicating a relatively balanced integration-stability profile. Harmony and scGPT remained utility-priority methods, reflecting strong integration performance with comparatively lower structural stability. Conversely, scFoundation and CellFMmin demonstrated favorable robustness but insufficient baseline integration quality to enter the preferred region. These patterns reinforce that integration quality and structural stability are not uniformly aligned across methods. Drivers analysis further revealed systematic statistical associations between integration performance and dataset structural properties. Across methods, the number of batches exhibited the largest average absolute standardized coefficients in five of the six evaluated metrics, indicating that integration performance is predominantly associated with batch complexity (Supplementary Fig. 32a). In contrast, for ASW, the number of cell types showed the largest average effect size, suggesting that geometric separability after integration is more strongly influenced by biological population complexity. Other structural variables, including total cell number and gene number, showed comparatively smaller effects across metrics. Although effect magnitudes varied across methods, the direction of association was largely consistent (Supplementary Fig. 32b and Supplementary Fig. 33). These findings indicate that variation in integration performance under zero-shot representations is primarily shaped by batch number, with biological complexity exerting a more specific influence on geometry-based conservation metrics.

**Fig. 5.**
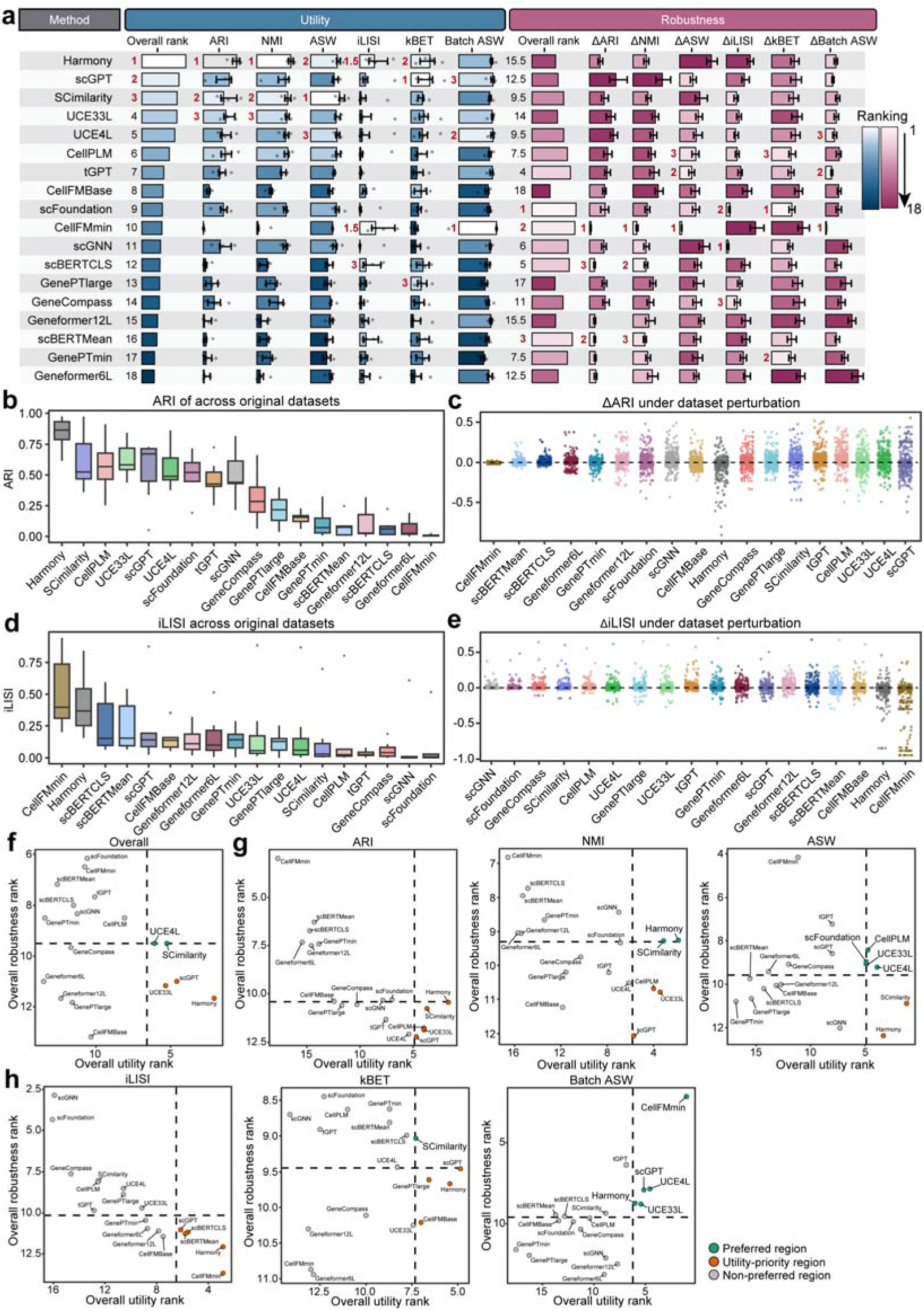
Batch effect correction under zero-shot representations. **a,** Performance overview across original datasets (n=7) and perturbation-derived datasets (n=325). Utility and robustness were summarized using rank-based aggregation across biological conservation and batch mixing metrics. **b,d,** Boxplots showing performance distributions of 18 methods across original datasets measured by ARI and iLISI, respectively. **c,e,** Scatter plots showing performance deviations under perturbation-derived datasets for ARI and iLISI, respectively. Each point represents one perturbation instance. **f,** Utility-robustness landscape under overall ranking across all integration metrics. Each point represents one method; lower ranks indicate better performance. Colors denote predefined decision regions based on asymmetric thresholds (top quartile for utility and median for robustness). **g,** Utility-robustness landscapes for biological conservation metrics. **h,** Utility-robustness landscapes for batch mixing metrics. iLISI, integration Local Inverse Simpson’s Index; kBET, k-nearest neighbor Batch Effect Test.

### Gene function prediction under zero-shot representations

We next evaluated whether pretrained gene embeddings encode functional semantics sufficient to support tissue-specific gene classification. In contrast to cell-level tasks, this analysis directly used zero-shot gene embeddings as input features. A total of 1,797 genes annotated to 15 tissue-specific categories from the Human Protein Atlas (HPA) (*35*) were used to construct a single-label multiclass task (Methods). Five-fold cross-validation was applied. Only models providing explicit gene embeddings were included, resulting in 12 foundation models. A unified two-layer multilayer perceptron (MLP) classifier was trained for all embeddings, and a one-hot encoding baseline was included as a non-semantic control. Gene function prediction was assessed on original datasets (utility) and on 180 datasets generated through perturbations of dataset structural properties (robustness). Performance was quantified using F1-score, AUROC and AUPRC.

Under original dataset conditions, models exhibited clear stratification. GeneCompass achieved the highest overall utility, followed by scGPT and Geneformer6L (*12*) (Fig. 6a, Supplementary Fig. 34-35). Rankings were largely consistent across metrics. The one-hot baseline performed near random expectation, indicating that gene identity alone is insufficient for tissue-specific classification. Cross-validation variance was low, suggesting stable performance estimation (Supplementary Fig. 36). Under perturbations of dataset structural properties, robustness rankings differed from utility. CellFMmin and scFoundation exhibited relatively smaller performance deviations, whereas GeneCompass showed larger fluctuations (Supplementary Fig. 37-39). As observed in other tasks, higher utility did not necessarily imply greater robustness.

Mapping models into the two-dimensional utility-robustness space revealed that scGPT and Geneformer6L fell within the preferred region, whereas GeneCompass showed high utility but did not consistently satisfy the robustness threshold (Fig. 6b). Notably, under AUPRC, which is more sensitive to class prevalence, only scGPT remained in the preferred region, whereas other high-utility models shifted toward a utility-priority region. These patterns indicate that functional classification accuracy and stability are only partially aligned under zero-shot gene embeddings. Drivers analysis revealed systematic statistical associations between gene function prediction performance and dataset structural properties. Across methods, the number of training classes exhibited the largest average absolute standardized coefficients across all metrics, indicating that performance is primarily associated with the breadth of label space observed during training (Supplementary Fig. 40a). The number of training genes showed second-largest positive associations, whereas class imbalance ratio exerted comparatively smaller and predominantly negative effects. Weighted train-test class overlap showed the moderate effects and was consistently positive. Although effect magnitudes varied among methods, the direction of association was broadly consistent (Supplementary Fig. 40b). Overall, these findings indicate that gene-level prediction performance is more closely associated with label-space structure than with dataset size or class imbalance alone.

**Fig. 6.**
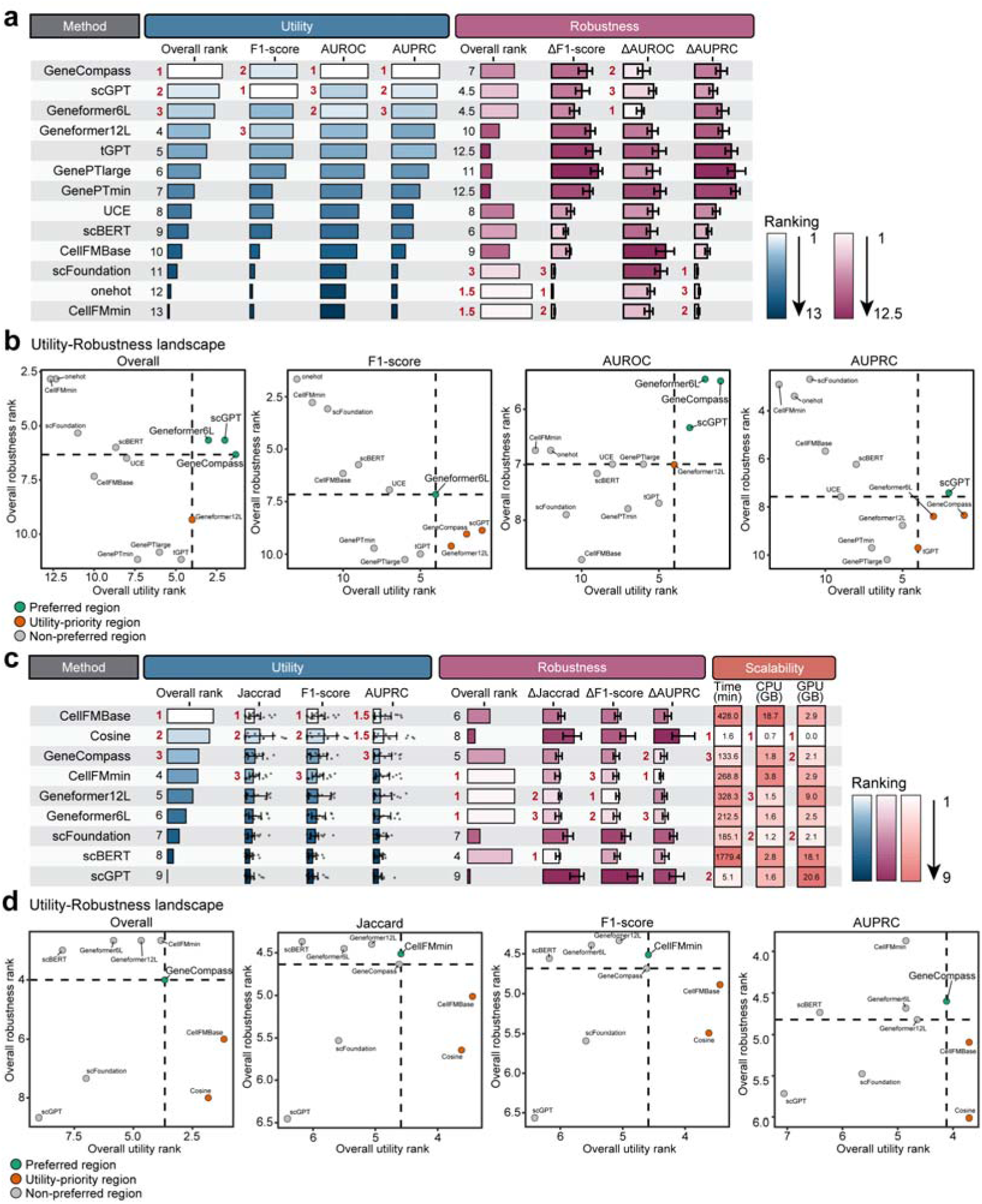
Gene-level tasks under zero-shot representations. **a,** Performance overview for gene function prediction across original datasets (n=1) and perturbation-derived datasets (n=120). Utility and robustness were summarized using rank-based aggregation across F1-score, AUROC and AUPRC. **b,** Utility-robustness landscapes for gene function prediction under overall ranking and individual metrics. Each point represents one of the 13 evaluated methods; lower ranks indicate better performance. Colors denote predefined decision regions based on asymmetric thresholds (top quartile for utility and median for robustness). **c,** Performance overview for perturbation-derived gene regulatory network inference across original datasets (n=17), perturbation-derived datasets (n=167), and computational cost (runtime, CPU memory and GPU memory). Utility and robustness were summarized using rank-based aggregation across Jaccard, F1-score and AUPRC. **d,** Utility-robustness landscapes for gene regulatory network inference under overall ranking and individual metrics. Each point represents one of the 9 evaluated methods; lower ranks indicate better performance. Colors denote predefined decision regions.

### Gene regulatory network inference under zero-shot representations

We next evaluated zero-shot representations for gene regulatory network (GRN) inference across 17 single-cell datasets. This task assesses the ability of pretrained gene association representations to recover transcription factor (TF)-target regulatory relationships defined by combined expression-response and motif-support criteria (Methods). Eight foundation models and one cosine similarity baseline were included. For all models, gene association matrices were extracted directly from pretrained weights without task-specific fine-tuning. Network aggregation and ranking procedures were standardized across methods. Performance was quantified using Jaccard index, F1-score and AUPR, capturing both early retrieval precision and overall ranking quality. Utility was evaluated on the 17 original datasets, robustness on 167 datasets generated through perturbations of dataset structural properties.

Under original dataset conditions, GRN recovery performance showed clear stratification (Fig. 6c and Supplementary Fig. 41a,b,c). CellFMBase ranked first overall, achieving the highest performance in both Jaccard and F1-score and performing comparably to the Cosine baseline in AUPR (Supplementary Fig. 42-43). The Cosine similarity baseline ranked second and exhibited consistently strong performance across all three metrics. GeneCompass followed as a stable third-tier method. In contrast, scBERT and scGPT showed comparatively lower recovery performance. These results indicate substantial differences across models in their ability to capture regulatory associations under zero-shot settings. Under perturbations of dataset structural properties, robustness rankings diverged from utility rankings (Supplementary Fig. 41d,e,f and Supplementary Fig. 44-45). CellFMmin and Geneformer variants exhibited smaller performance deviations across metrics and ranked highly in overall robustness. In contrast, CellFMBase and the cosine baseline, despite strong utility, showed larger fluctuations under structural perturbations. As observed in previous tasks, higher utility did not consistently correspond to greater robustness.

Mapping models into the two-dimensional utility-robustness space further clarified this trade-off. GeneCompass occupied the preferred region, reflecting a relatively balanced profile between recovery accuracy and stability (Fig. 6d). CellFMBase and Cosine remained utility-priority methods, characterized by strong baseline recovery but reduced robustness. Conversely, CellFMmin and Geneformer variants demonstrated favorable robustness yet insufficient baseline recovery performance to enter the preferred region. Drivers analysis revealed that ground-truth target set size exhibited the largest average absolute standardized coefficients across metrics, indicating that GRN recovery performance is strongly associated with the size of the positive regulatory set (Supplementary Fig. 46). Across models, the direction of association was largely consistent, with heterogeneity primarily reflected in effect magnitude. These findings suggest that GRN inference performance under zero-shot representations is influenced more by candidate space scale than by sample composition. In addition, computational profiling showed substantial differences in runtime and memory requirements across methods (Fig. 6c). The cosine baseline was the most efficient and did not require GPU resources, whereas larger foundation models incurred higher GPU memory consumption. As in previous tasks, computational cost did not directly track predictive utility, underscoring trade-offs among utility, robustness and resource efficiency in practical applications.

## DISCUSSION

In this study, we established a structured framework to evaluate scFMs under routine analytical constraints. All embeddings are extracted strictly in a zero-shot manner, downstream pipelines are standardized, and performance is assessed on original datasets as well as under perturbations of dataset structural properties. Across 6 tasks and 1,607 datasets, no single method maintained consistently strong performance across datasets. Instead, method preferences exhibit strong task dependency, indicating that zero-shot performance cannot be reduced to a single global ranking.

Joint evaluation of utility and robustness revealed additional performance space beyond baseline rankings. For cell clustering UCE4L and CellPLM satisfied the joint utility-robustness criteria; scFoundation met the joint thresholds in cell type annotation; and highVariable together with scFoundation showed balanced performance in drug sensitivity prediction. In batch integration, Harmony ranked highest in utility, whereas UCE4L and SCimilarity met both thresholds. At the gene level, scGPT and Geneformer6L satisfied the joint criteria for gene function prediction, whereas GeneCompass showed high utility with lower robustness. In gene regulatory network inference, GeneCompass met the joint criteria, whereas a cosine-similarity baseline showed competitive utility but weaker robustness. These patterns indicate that zero-shot generalization in scFMs cannot be inferred from utility alone and is better understood as a balance between utility and robustness under realistic analytical variability.

Several broader implications emerge from these cross-task comparisons. First, utility and robustness can decouple. Methods that rank highly on original datasets can exhibit larger paired deviations under structural perturbations, as observed for SCimilarity in clustering and for Harmony and scGPT in batch integration. Reporting both baseline utility and perturbation-based deviation therefore improves the reliability of method recommendations, particularly when dataset composition differs from benchmark conditions. Second, baseline methods remain practically competitive in specific contexts when downstream pipelines are standardized and fine-tuning is avoided. For example, highVariable performed strongly in supervised annotation and met joint criteria in drug sensitivity prediction, and Harmony achieved the highest utility in batch integration. These observations suggest that gains from large pretrained embeddings are task- and data-dependent rather than universal. Third, computational cost varied substantially across methods and did not consistently track performance. Resource constraints can therefore rationally influence method choice without implying uniform losses in accuracy. This observation aligns with the accessibility challenges for large single-cell models (*6*) and supports reporting comparison of resource requirements alongside performance metrics when introducing new scFMs. In addition, drivers analyses indicated a small set of dataset descriptors that consistently aligned with performance differences across tasks, such as overlap of classes in supervised settings and batch number in integration. Although these associations are descriptive rather than causal, they define practical axes for stress-testing new representations.

Together, these findings refine the interpretation and boundaries of zero-shot generalization in scFMs. Utility on a fixed benchmark captures one dimension of practical value, but stability under shifts to dataset structure is a separate and complementary dimension that can reorder methods. A comprehensive evaluation therefore benefits from pairing standard utility summaries with robustness under perturbations of dataset structural properties and from presenting recommendations conditional on dataset structure and user constraints.

Several limitations should be considered. First, we intentionally avoided fine-tuning and used standardized downstream heads to isolate representation content. As a result, best-case performance achievable through extensive tuning or customized architectures was not evaluated. Second, perturbations were defined along selected structural properties and do not encompass all sources of biological and technical variability. Third, the drivers analyses are associative and may reflect residual confounding; they should therefore not be interpreted as evidence of causality. Finally, gene-level tasks necessarily excluded models that do not expose reusable gene embeddings or gene-gene relations, reflecting current heterogeneity in model interfaces and reporting practices within the ecosystem.

Looking forward, as single-cell datasets scale to millions of cells (*36, 37*), the computational burden of task-specific fine-tuning renders robust zero-shot representations a fundamental requirement for downstream analysis. Consequently, evaluating single-cell foundation models must shift from incremental leaderboard gains to sustained efficacy in dynamic, real-world environments characterized by class imbalance, incomplete annotations, and technical variations (*38*). Models optimizing fixed benchmarks must prove their structural resilience in prospective biological studies. Future progress will therefore depend less on scaling parameters (*6, 39*) and more on standardized stress-testing paradigms that capture analytical volatility. Validating models across complex, multi-institutional cohorts is essential to ensure they serve as reliable inference engines. Ultimately, reframing generalization as a context-dependent property defined by data structure, task abstraction, and computational constraints provides a principled foundation for next-generation modeling.

## MATERIALS AND METHODS

### Experimental Design

To evaluate scFMs across various downstream tasks, we quantified performance along three complementary dimensions: utility on original datasets, robustness to perturbations of dataset structural properties, and exploratory associations with dataset-level characteristics. The same statistical framework was applied across all tasks, with task-specific performance metrics substituted accordingly.

#### (1) Utility on original datasets

Utility was assessed using original datasets. Let *m* ∈{1,…,*M*}denote methods, *i* ∈{1,…,*N*} denote original datasets and *y_i,m_* denote the performance metric of method *m*on dataset *i*. All methods were evaluated on the same set of datasets within each task, yielding a paired comparison structure across methods. Because evaluation metrics may differ in scale, variance and dynamic range across datasets, direct aggregation of raw metric values may overweight datasets with larger numeric ranges. To obtain a scale-invariant summary of relative performance consistency, we adopted a rank-based aggregation strategy. Within each dataset *i*, methods were ranked according to their performance:

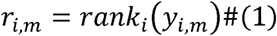

where lower rank indicates better performance. Tied values were assigned average ranks. For each method:

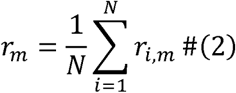

This quantity reflects the average relative standing of the method *m* across heterogeneous datasets. When multiple metrics were used within a task, method-wise average ranks were again ranked and averaged across metrics to yield an overall utility rank. This two-level ranking procedure emphasizes performance consistency rather than absolute scale and mitigates cross-dataset heterogeneity. Furthermore, for pairwise comparison between methods A and B, paired differences were defined as:

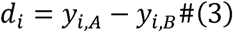

One-sided Wilcoxon signed-rank tests were performed under the predefined directional hypothesis from above utility ranking that higher performance indicates superiority:

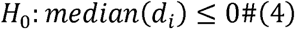

All pairwise comparisons were conducted within each metric. P values were adjusted using the Benjamini-Hochberg procedure within each metric to control the false discovery rate. Adjusted p-value < 0.05 was considered statistically significant. The median paired difference,

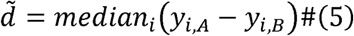

was reported as an effect size to quantify the magnitude of utility differences. Positive values of 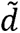 indicate that method A achieves higher performance than method B on most datasets.

#### (2) Robustness under dataset perturbations

To quantify robustness under perturbations of dataset structural properties, we measured the absolute raw deviation of model performance on each perturbed dataset relative to its original counterpart. It provides a direction-invariant measure of stability, as perturbations can induce both performance gains and losses. For each method *m* and each original dataset *i*, baseline performance was defined as 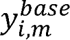. From each original dataset, zero or more perturbed datasets were generated after quality control, indexed by *k*=1,…,*k_i_*, where perturbations modified predefined structural properties, such as cell number, to simulate diverse data conditions. Performance under perturbation was denoted: 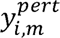. Performance deviation was defined as:

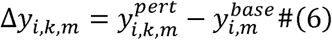

This formulation ensures that deviations reflect structural modification relative to a matched baseline rather than differences across unrelated datasets. When *k_i_* = 0, the corresponding baseline dataset did not contribute to robustness analysis. Robustness was operationalized as the magnitude of deviation:

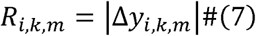

Each perturbation instance (*i*,*k*) was treated as a distinct structural scenario. Within each perturbation scenario, methods were ranked according to *R_i,k,m_* where lower values indicate greater stability:

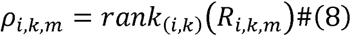

For each method:

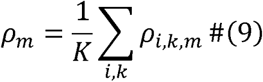

where 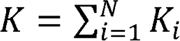 denotes the total number of perturbation scenarios. Cross-metric aggregation followed the same rank-based procedure described for utility analysis. Because robustness comparisons were conducted within identical perturbation scenarios across methods, paired differences were preserved at the perturbation level. For pairwise comparison between methods A and B, paired differences were defined as:

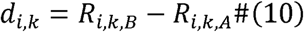

One-sided Wilcoxon signed-rank tests were performed under the predefined directional hypothesis that smaller deviations indicate greater robustness:

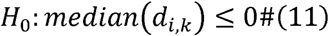

Thus, rejection of the null hypothesis implies that method A exhibits significantly smaller performance deviations than method B. All pairwise comparisons were conducted within each metric. P values were adjusted using the Benjamini–Hochberg procedure within each metric to control the false discovery rate. Adjusted *p-value* < 0.05 was considered statistically significant. The median paired difference,

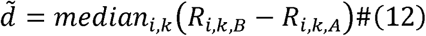

was reported as an effect size to quantify the magnitude of robustness differences.

#### (3) Associations with dataset structural properties

Drivers analysis was designed as an exploratory assessment of whether model performance is systematically associated with dataset structural properties. This analysis was conducted independently for each task, performance metric, and method. Let *i*=1,…,*N* denote the index of each dataset, *m* denote a specific method and *k* denote a specific evaluation metric. For a given method *m* and metric *k*, we define *y_i_^(m,k)^* as the performance value of method *m* on dataset *i*. For each dataset, we define a vector of structural features: 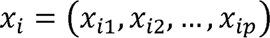, where each *X_ij_* corresponds to a dataset-level characteristic. All analyses were conducted separately for each (*m*,*k*) combination. To address multicollinearity among dataset structural features and reduce overfitting risk, we applied Elastic Net regression using the *glmnet* package in R. The mixing parameter was fixed at *α* = 0.5, assigning equal weight to L1 and L2 penalties. Regularization strength was determined via 10-fold cross-validation, and the optimal value was selected as λ*_min_*, corresponding to the minimum cross-validated mean squared error. Predictors were internally centered and scaled to unit variance during penalized fitting. Features with non-zero coefficients at λ*_min_* were retained for subsequent regression analysis. If no features were selected for a given (*m*,*k*), the model was excluded from further analysis. To estimate directionality and statistical significance, we refitted an ordinary least squares (OLS) regression model using the Elastic Net-selected predictors. Before OLS fitting, selected predictors were standardized:

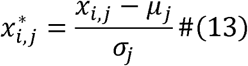

where μ*_j_* and σ*_j_* denote the mean and standard deviation of predictor *j* across datasets. The regression model was specified as:

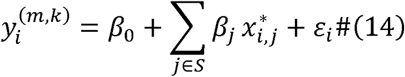

where *S* denotes the set of selected predictors. Because predictors were standardized, the estimated coefficients β*_j_* represent standardized effect sizes and are directly comparable across features. To avoid overinterpretation of weak explanatory models, only regressions with R^2^ ≥ 0.1 were retained for reporting and visualization. The Drivers analysis is exploratory and intended to identify systematic statistical associations between dataset structural properties and model performance. It does not imply causal relationships.

### Model variants of scFMs

To ensure comprehensive and practically relevant benchmarking, multiple variants of specific scFMs were included if the original authors released distinct pre-trained checkpoints or recommended alternative readout strategies. Specifically, we evaluated the publicly available 4-layer and 33-layer checkpoints of UCE (denoted UCE4L and UCE33L), and for Geneformer we included the released 6-layer and 12-layer checkpoints pretrained on GeneCorpus-30M (Geneformer6L and Geneformer12L). For GenePT, we benchmarked the two released configurations (GenePTmin and GenePTlarge) based on different text-embedding backbones used to encode gene summaries. For CellFM, we included the released parameter-scale variants (CellFMmin, ∼80M; CellFMBase, ∼800M). In addition, we compared two representation strategies of scBERT: the first token representation and global mean pooling over gene tokens. All variants were subjected to identical zero-shot evaluation protocols and were treated as independent models throughout the study.

### Task 1: Cell clustering

#### Dataset and preprocessing

Tabula Sapiens 2.0 (*40*) was used to benchmark clustering performance across diverse human tissues and to reduce potential overlap with model pretraining resources. After stratifying by tissue of origin, subsets containing fewer than two cell types or fewer than 1,000 cells were excluded. We applied uniform quality control by filtering genes expressed in fewer than three cells (min.cells = 3) and removing cells with fewer than 1,000 detected genes (min.features = 1000) in R package Seurat R package (*41*). This yielded 26 datasets comprising 541,276 cells (Supplementary Table 2).

#### Representation extraction and baselines

For each dataset, cell embeddings were extracted from each single-cell foundation model under the zero-shot setting, with no task-specific fine-tuning. These embeddings were used as the sole input features for downstream clustering. As baselines, we included (i) PCA using the top 50 principal components computed from the top 2,000 highly variable genes (HVGs), and (ii) scGNN, a self-supervised deep learning approach that learns low-dimensional cell representations.

#### Downstream protocol and evaluation

Clustering was performed using the scIB framework (*31*). For each representation, we constructed a k-nearest neighbor (kNN) graph (n_neighbors=15) and selected the clustering resolution that maximized NMI with ground-truth cell-type labels. This optimal-resolution setting was used to standardize cross-model comparisons and isolate representation quality from resolution choice. Clustering accuracy and separability were quantified by ARI, NMI, and ASW computed on cell types in the embedding space.

#### Robustness perturbations and dataset features

To quantify robustness to changes of dataset structure, we generated 339 perturbed datasets derived from the 26 datasets by (i) random gene subsampling (retaining 80%, 60%, 40% and 20% of genes), (ii) proportional cell subsampling using the same gradients, and (iii) targeted removal of selected cell types. The full clustering pipeline was repeated for each perturbed dataset. To support downstream factor analysis, we computed four dataset features: cell count, gene count, number of cell types, and class imbalance ratio (maximum-to-minimum cell type abundance) (Supplementary Table 1).

### Task 2: Cell annotation

#### Dataset and preprocessing

We used the same 26 datasets as in Task 1. Within each dataset, we performed stratified 5-fold cross-validation by cell-type labels, preserving class proportions across folds. To avoid untrainable splits, each training fold was constrained to contain all cell types present in its corresponding test fold. This design produced 130 train-test splits and 2,706,380 total cell instances across folds (Supplementary Table 3).

#### Representation extraction and baselines

All foundation models generated cell embeddings in the zero-shot manner, without any fine-tuning toward classification. Training and test folds were embedded independently, disallowing the sharing of intermediate statistics. For the PCA baseline, HVGs (top 2,000) were identified using training folds only to prevent information leakage, and the resulting gene set was applied to the paired test fold. scGNN was included as a self-supervised representation baseline. All data-dependent operations were performed on training folds only, and test folds were used exclusively for evaluation.

#### Downstream protocol and evaluation

To attribute performance differences to representation quality, we trained a standardized fully connected classifier comprising two hidden layers of 512 and 256 units with ReLU activations and dropout, on top of each embedding. Within a 5-fold cross-validation framework, each training fold was further partitioned into a 9:1 stratified training and validation split. We optimized the models using multi-class cross-entropy loss and the AdamW algorithm. To mitigate overfitting, we employed dynamic learning rate decay and early stopping, specifically retaining the model weights from the final epoch at the point of termination. We report macro-averaged F1-score, AUROC and AUPRC, denoted as F1-score, AUROC and AUPRC, evaluated on the held-out test folds.

#### Robustness perturbations and dataset features

To assess robustness to feature and training-distribution shifts, we constructed 328 perturbed variants from the training folds while keeping the corresponding test folds fixed. Perturbations included (i) proportional gene subsampling (80%, 60%, 40%, 20%), (ii) stratified cell downsampling at the same gradients while preserving class proportions, and (iii) random exclusion of selected cell types from training, retaining at least two classes. The embedding and classification pipeline was repeated for each perturbation. To support downstream factor analysis, we computed five dataset features: training cell count, training gene count, training cell-type count, imbalance ratio (maximum-to-minimum cell type abundance), and the proportion of train-test cell-type overlap, weighted by test-set cell abundance (Supplementary Table 3).

### Task 3: Drug response prediction

#### Dataset and preprocessing

We evaluated drug response prediction using single-cell drug response datasets from DRMref(*33*). Drug responses were binarized into Resistant (R) and Non-resistant (NR). After quality control (min.cells = 3, min.features = 1000), 14 datasets comprising 93,481 cells were retained. For each dataset, we performed stratified 5-fold cross-validation by R/NR labels and used identical splits across all methods (Supplementary Table 4).

#### Representation extraction and baselines

Cell embeddings were used as the input features. Foundation models produced embeddings under the zero-shot setting without access to drug response labels. As an expression-based baseline, we used HVG features constructed from the top 2,000 HVGs selected within each training fold. scGNN embeddings were obtained via its self-supervised procedure. Embedding extraction for training and test folds was performed independently to avoid leakage, and all data-dependent operations were confined to training folds.

#### Downstream protocol and evaluation

To predict drug sensitivity, we implemented a dual-tower neural network that integrates cell embeddings with drug molecular features. PubChem SMILES (*42*) of drug was converted to 1,024-dimensional Morgan fingerprints in RDKit (radius = 2). Separate cell and drug encoders, each comprising a linear projection, ReLU activation, batch normalization, and dropout, mapped the respective inputs into a shared latent space. These representations were concatenated and processed by a two-layer fusion network to output sensitivity predictions. Within a 5-fold cross-validation framework, each training fold was further partitioned into a 9:1 stratified training and validation split. Models were trained using the AdamW optimizer and binary cross-entropy loss with logits, employing early stopping based on validation accuracy to mitigate overfitting. We report F1-score, AUROC and AUPRC on test folds.

#### Robustness perturbations and dataset features

We generated 177 perturbed variants to evaluate robustness to dataset structural shifts, including (i) gene subsampling in training folds (80%, 60%, 40%, 20%) and (ii) stratified cell subsampling in training folds at the same gradients. In addition, we constructed two generalization settings: unseen cell line (cell lines strictly separated between training and test) and unseen drug (drugs strictly separated between training and test). We repeated the full prediction pipeline for all perturbations and computed dataset features including cell and gene counts, class imbalance ratio, and modality overlap rates for factor analysis (Supplementary Table 4).

### Task 4: Batch integration

#### Dataset and preprocessing

Following two established batch-correction benchmarking studies(*31, 43*), we selected seven representative single-cell datasets spanning cross-technology or cross-experimental conditions. Quality control was performed as above (min.cells = 3, min.features = 1000). In total, 7 datasets comprising 105,750 cells were retained (Supplementary Table 5).

#### Representation extraction and baselines

All foundation models were evaluated in a zero-shot setting by extracting cell embeddings without incorporating batch labels or downstream fine-tuning. Harmony(*34*) was included as a baseline method. scGNN provide cell embeddings from the latent of its a self-supervised model. All embeddings were used as the sole representations for downstream evaluation.

#### Downstream protocol and evaluation

Batch correction performance was evaluated using scIB. For each representation, we constructed a kNN graph (n_neighbors = 15) and selected clustering resolution by maximizing NMI. We quantified two complementary aspects: biological conservation (ARI, NMI, cell-type ASW) and batch mixing (batch ASW, iLISI, kBET).

#### Robustness perturbations and dataset features

We constructed 325 perturbed variants to assess robustness to changes in dataset structure: (i) gene subsampling (80%, 60%, 40%, 20%), (ii) cell downsampling (80%, 60%, 40%, 20%) while preserving batch and cell-type proportions, (iii) random removal of one or multiple batches, (iv) random removal of one or multiple cell types, and (v) hierarchical perturbations by jointly removing selected batch–cell-type combinations. The evaluation pipeline was repeated for each perturbed dataset. We computed four dataset features: cell count, gene count, cell type count, and batch count to support factor analysis (Supplementary Table 5).

### Task 5: Gene function prediction

#### Dataset, labels and model selection

To evaluate whether foundation models encode gene semantics in their gene-level embeddings, we curated 1,797 single-tissue-specific genes across 15 tissue labels from the HPA and formulated a single-label multi-class classification benchmark with 5-fold cross-validation (Supplementary Table 6). Because this task requires reusable gene dictionaries, we screened the 18 previously considered models for eligibility. Models without independent gene vocabularies (CellPLM, SCimilarity) and graph-centric methods (scGNN) were excluded. Models sharing identical gene-to-index dictionaries, notably scBERT and the UCE series, were consolidated to avoid redundant evaluations. In total, 13 models with explicit gene-level embeddings were retained and evaluated under a zero-shot setting with frozen pretrained weights. As a non-semantic baseline, we used one-hot vectors.

#### Downstream protocol and evaluation

All gene representations were evaluated using a standardized two-layer MLP classifier incorporating ReLU activation and dropout. To align the evaluation sets, analysis was restricted to the intersection of genes shared across all embedding dictionaries and the HPA dataset. Performance was assessed within a stratified five-fold cross-validation framework. Models were optimized for 50 epochs using the Adam optimizer with cross-entropy loss. We report macro-averaged F1, AUROC and AUPRC (one-vs-rest), denoted as F1-score, AUROC and AUPRC in the text.

#### Robustness perturbations and dataset descriptors

We generated 180 perturbed variants across three robustness aspects: (i) gene subset sampling, where training folds were subsampled (80%, 60%, 40%, 20%) under fixed test sets and class-balance constraints; (ii) class ratio perturbation, where class abundances were reshuffled and down-weighted from 1.0 to 0.1 to induce increasing imbalance; and (iii) class overlap perturbation, where 1-5 randomly selected classes were held out entirely as unseen classes and assigned to the test set, while the remaining visible classes were split 8:2 for training/testing. For each perturbation, we repeated the prediction pipeline and computed dataset features including training gene count, training gene class count, imbalance ratio (maximum-to-minimum gene class abundance), and train-test overlap for factor analysis (Supplementary Table 6).

### Task 6: Gene regulatory network inference

#### Dataset and ground-truth construction

We evaluated foundation model capabilities for GRN inference using perturbation single-cell datasets and a ground-truth definition for TF targets. We curated 17 datasets comprising 9,096 cells, each corresponding to TF overexpression (OE) or knockdown/knockout (KD/KO) experiments. Controls were downsampled to match perturbed cells to mitigate imbalance, followed by quality control (min.cells = 3, min.features = 1000). Ground-truth TF targets were defined by a two-evidence criterion combining expression response and motif support: (i) “expression-responsive genes” identified by Wilcoxon tests (KD/KO: log2FC < −0.25, p-value < 0.05; OE: log2FC > 0.25, p-value < 0.05), and (ii) motif-supported candidates defined as the top 5,000 genes ranked by the corresponding TF motif in the RcisTarget matrix(*44*). The intersection of the two sets was used as high-confidence targets. Datasets with fewer than five targets were excluded to reduce indirect cascade effects (Supplementary Table 7).

#### Network inference and baselines

We evaluated 8 scFMs and 1 statistical baseline under a zero-shot setting. For scBERT, Geneformer series, scGPT and CellFM series, gene-gene associations were derived from model attention matrices. For scFoundation and GeneCompass, networks were inferred by cosine similarity between gene embeddings following their native designs. As a baseline without pretrained priors, we L2-normalized single-cell expression vectors and computed cosine similarities to obtain a gene association matrix. All association matrices were evaluated using an identical pipeline.

#### Downstream protocol and evaluation

For each TF, gene association matrices were aggregated separately for perturbed and control cells by averaging across cells. Dynamic regulatory strength from the TF to each gene was defined as the absolute difference between the two mean matrices. After excluding TF self-loops, genes were ranked by this score. We evaluated high-confidence recovery using Jaccard index and F1 at a Top-K threshold (top 5%), and assessed ranking quality using AUPRC, denoted as Jaccard, F1-score and AUPRC in the text.

#### Robustness perturbations and dataset features

We generated 167 perturbed variants across the TF datasets, totaling 67,503 cells. Perturbations included (i) gene subsampling (80%, 60%, 40%, 20%) while retaining the target TF, (ii) proportional cell subsampling applied to both perturbed and control conditions (80%, 60%, 40%, 20%), and (iii) imbalanced subsampling where the perturbed-to-control cell ratio was shifted to 2:1 or 1:2 under the same downsampling gradients. We repeated the GRN pipeline for all perturbations and computed dataset features including gene count, cell count, imbalance ratio (case/control) and target gene count for factor analysis (Supplementary Table 7).

#### Computational resources

All experiments were conducted on a Linux server running Ubuntu 22.04.5 LTS, equipped with an AMD 32-core CPU and 503 GB system memory. GPU was a NVIDIA A100 PCIe GPU (40 GB). The CUDA toolkit version was 12.2. In addition, we profiled computational cost for two zero-shot representation stages: (i) cell embedding extraction, where each model produced cell-level embeddings from the input expression matrix without any task-specific fine-tuning; and (ii) gene association matrix extraction, where models produced gene-gene association matrices under the same zero-shot setting.

## General

We thank the providers of open-access databases whose resources made this study possible.

## Author contributions

T.L., H.L. and Y.Z. conceived and designed the study. T.L. and T.F. developed the methodology, implemented the software, performed the investigation and conducted the formal analyses. T.L. curated the datasets. T.L. and T.F. prepared the visualizations. X.P., Y.C. and L.R. provided resources. T.L. wrote the original draft. T.L., T.F., L.R., X.Y., H.L. and Y.Z. reviewed and edited the manuscript. X.Y., H.L. and Y.Z. supervised the study. All authors read and approved the manuscript.

## Data availability

All datasets used in this study were obtained from publicly accessible databases and are summarized in Supplementary Table S8. The scripts used in this study are available via GitHub at https://github.com/ShellyCoder/scFMBench.

