## Supplemental Note and Figure for "Context-dependent utility and robustness of pretrained single-cell foundation model representations across analytical tasks"

### Supplementary Note 1: Citation analysis of single-cell foundation models

#### Model inclusion criteria

Citation analysis was restricted to scFMs for which (i) a primary algorithm publication could be identified in Web of Science and (ii) citation records were available up to 31 December 2025. Based on these criteria, nine models were included: scBERT, Geneformer, CellFM, GenePT, scGPT, scFoundation, GeneCompass, tGPT and SCimilarity. Other representation methods were not included due to the absence of an identifiable primary publication indexed in Web of Science or insufficient citation records within the defined time window. For each included model, all citing publications indexed in Web of Science up to 31 December 2025 were retrieved. A total of 1,330 citing articles were retained for classification.

#### Literature classification criteria

All retained publications were manually reviewed and categorized to assess the context in which the cited model was referenced. The objective was to distinguish between conceptual mention, methodological comparison and direct downstream application.

Three citation categories were defined:

1. Applied citations (Category A) refer to studies in which the cited foundation model was directly used for downstream single-cell data analysis, including cell embedding construction, cell type annotation, drug response prediction, batch integration, gene function prediction or gene regulatory network inference.
2. Methodological citations (Category B) refer to studies that evaluated the model in comparative benchmarking, performance assessment or methodological discussion without direct downstream application.
3. General citations (Category C) refer to studies that mentioned the model in background discussion, reviews or broader conceptual contexts without method comparison or applied analysis.

#### Annotation procedure and agreement assessment

Two independent reviewers with backgrounds in computer science and biomedical engineering (Referee 1 and Referee 2) performed blinded classification according to predefined criteria. Classification decisions were based on titles and abstracts, with fulltext review conducted when necessary. Disagreements were resolved through discussion to reach a final consensus decision. Agreement of each reviewer relative to the final consensus classification was 0.9789 for Referee 1 and 0.9835 for Referee 2, indicating high reproducibility of the categorization scheme (Supplementary Table 1).

**Distribution of citation contexts**

Among the 1,330 included publications, general citations (Category C) accounted for 85.34%, methodological citations (Category B) for 13.31%, and applied citations (Category A) for 1.35% (n = 18) (Supplementary Fig. 1a). To examine disciplinary distribution, citation categories were further stratified according to Web of Science subject classifications. Applied citations were primarily concentrated in disease-oriented research areas, including immunology and oncology (Supplementary Fig. 1b). Methodological citations were more frequently observed in interdisciplinary domains such as computer science and biotechnology (Supplementary Fig. 1c). General citations were broadly distributed across biochemistry, molecular biology, computer science and computational biology (Supplementary Fig. 1d).

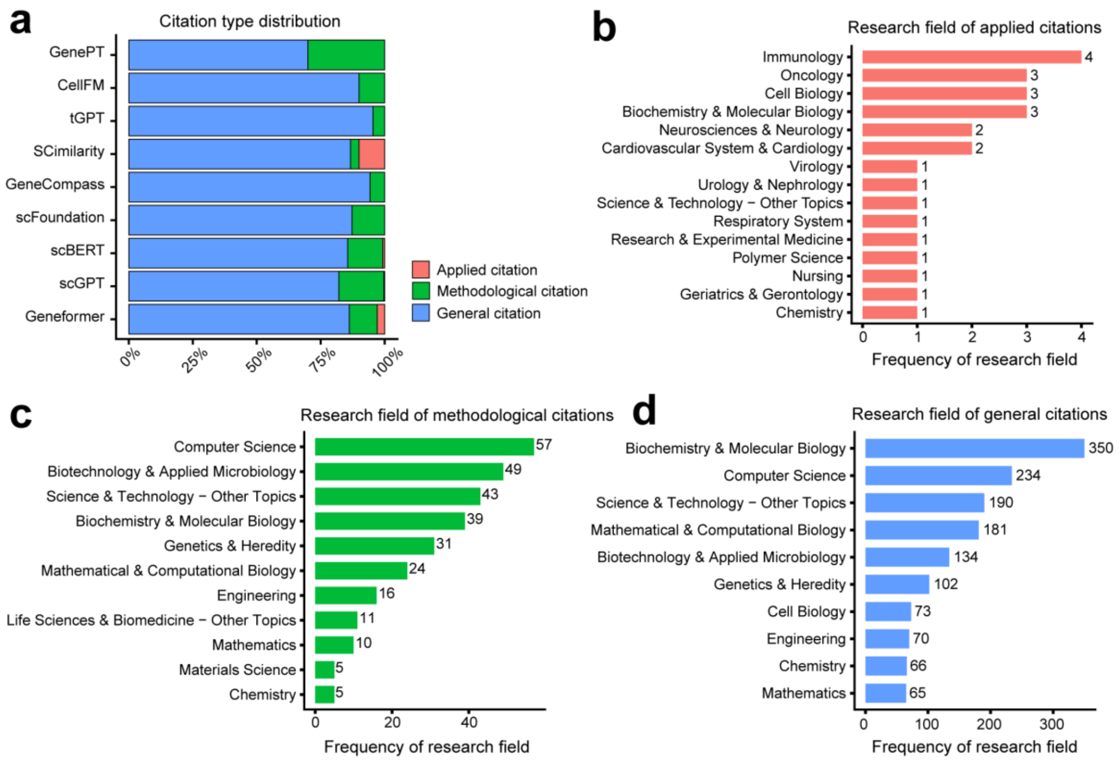


**Supplementary Fig. 1 | Citation analysis of single-cell foundation models. a,** Citation type distribution. Proportion of applied (Category A), methodological (Category B) and general (Category C) citations for each included model. Citation records (n=1,330) were retrieved from Web of Science up to 31 December 2025. **b,** Research fields of applied citations. Web of Science subject categories for publications in which foundation models were directly used in downstream single-cell data analysis (n = 18). **c,** Research fields of methodological citations. Web of Science subject categories for publications involving benchmarking, comparative evaluation or methodological discussion of foundation models without direct downstream application (n = 177). **d,** Research fields of general citations. Web of Science subject categories for publications that referenced foundation models in background or conceptual contexts without methodological comparison or applied use (n = 1,135). Counts correspond to the number of publications assigned to each literature classification.

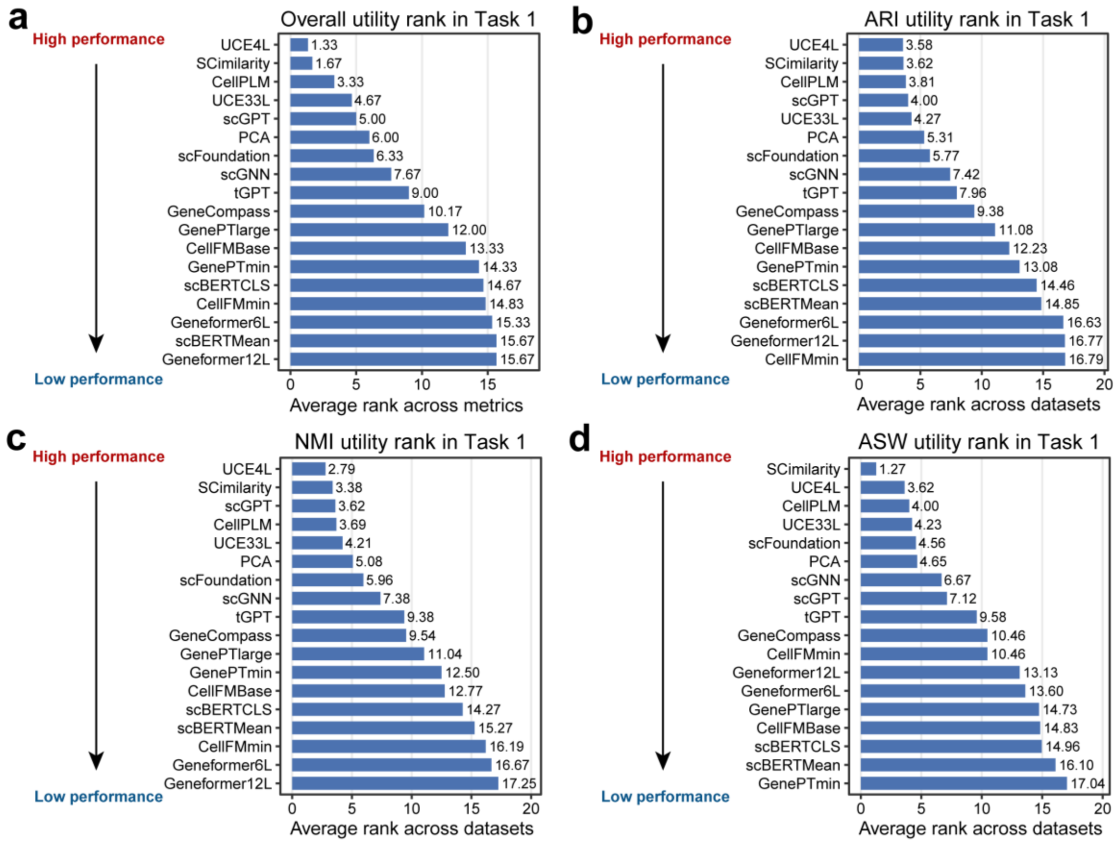


**Supplementary Fig. 2 | Average utility rankings for zero-shot cell clustering.** Bar plots display the overall aggregated rank across all three metrics **(a)**, alongside the individual average ranks across the 26 datasets for ARI **(b)**, NMI **(c)**, and ASW **(d)**. Lower numerical rank values indicate superior method performance.

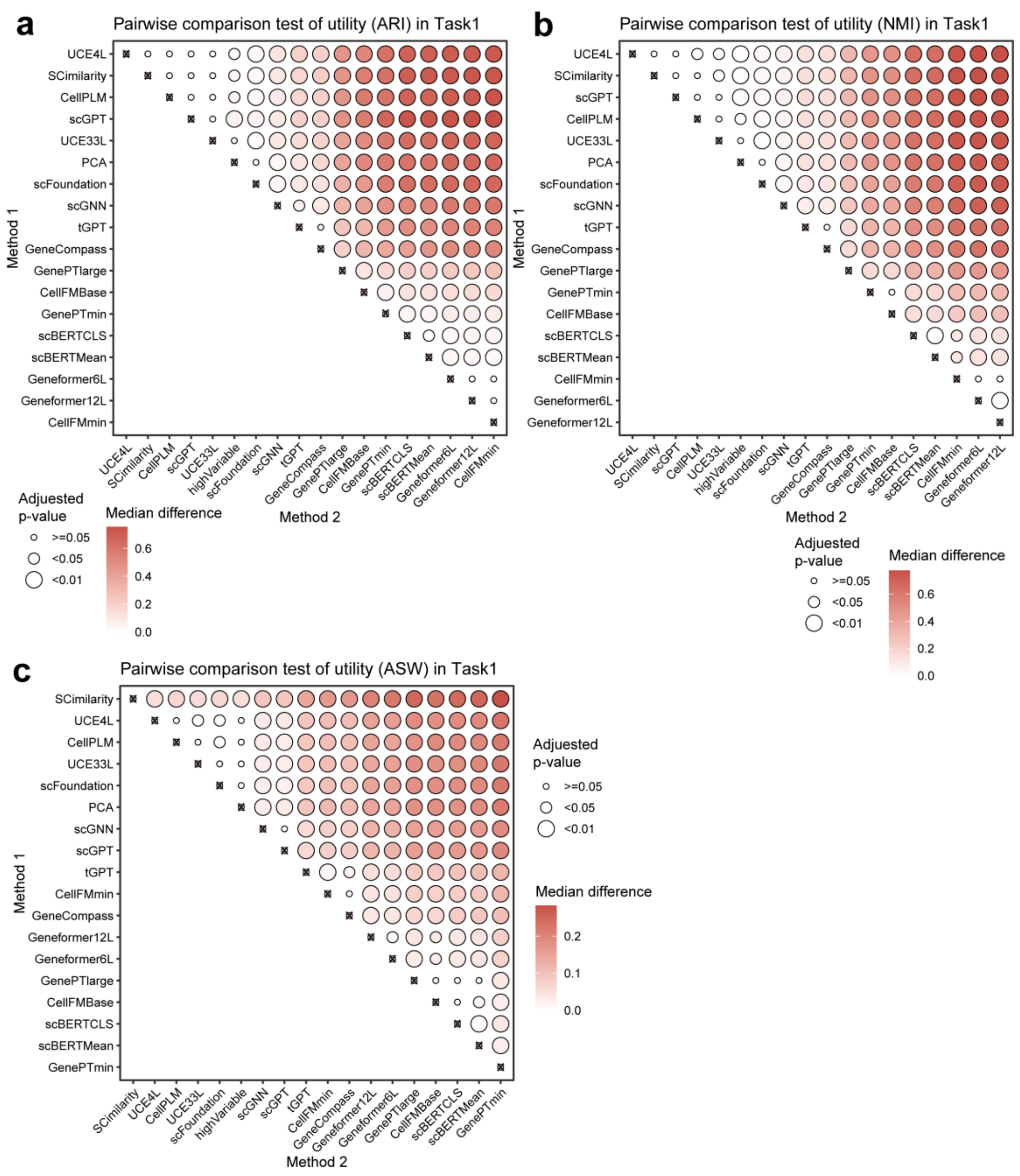


**Supplementary Fig. 3 | Pairwise statistical comparisons of zero-shot utility in cell clustering. a-c,** Bubble plots visualize the results of one-sided Wilcoxon tests comparing method performance across ARI **(a)**, NMI **(b)**, and ASW **(c)** metrics, with algorithms ordered along the axes by their independent average ranks. Bubble color intensity represents the median performance difference between the Method 1 and Method 2 algorithms, while bubble size indicates the adjusted p-value.

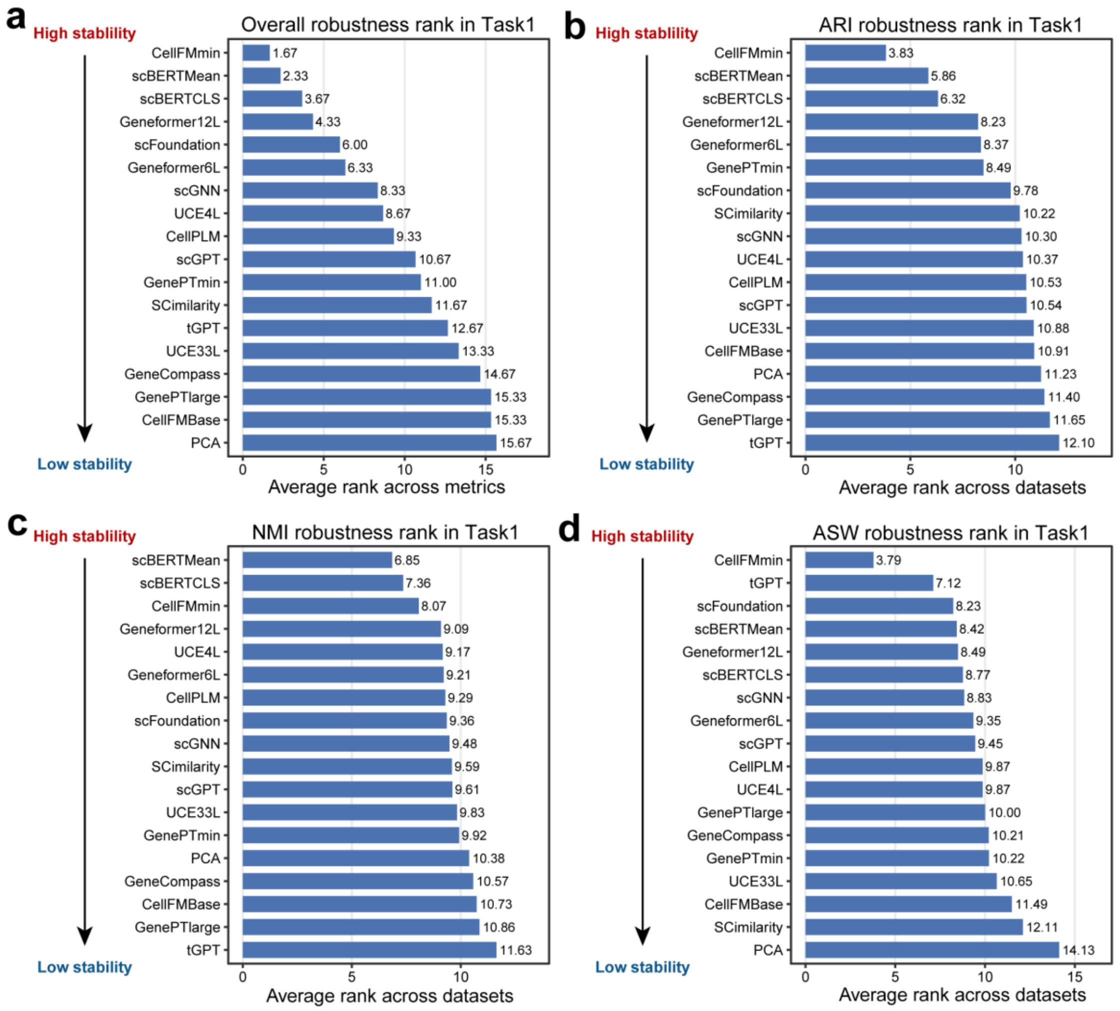


**Supplementary Fig. 4 | Average robustness rankings for zero-shot cell clustering.** Bar plots display the overall aggregated rank across all three metrics **(a)**, alongside the individual average ranks across the 339 datasets for ARI **(b)**, NMI **(c)**, and ASW **(d)**. Lower numerical rank values indicate superior robustness.

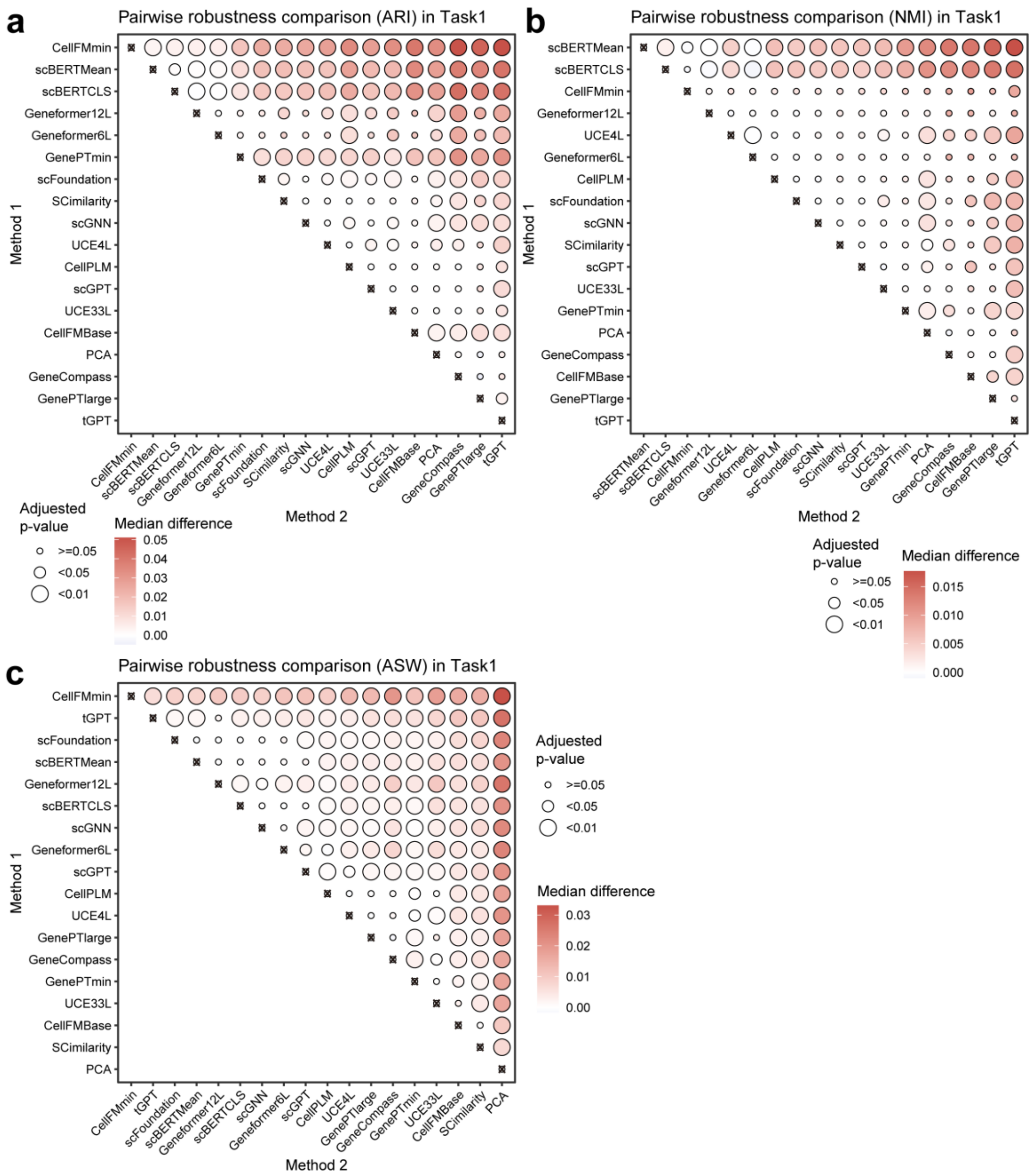


**Supplementary Fig. 5 | Pairwise statistical robustness comparisons of zero-shot cell clustering. a-c,** Bubble plots visualize the results of one-sided Wilcoxon tests comparing method robustness across ARI **(a)**, NMI **(b)**, and ASW **(c)** metrics, with algorithms ordered along the axes by their independent average ranks. Bubble color intensity represents the median robustness difference between the Method 2 and Method 1 algorithms, while bubble size indicates the adjusted p-value.

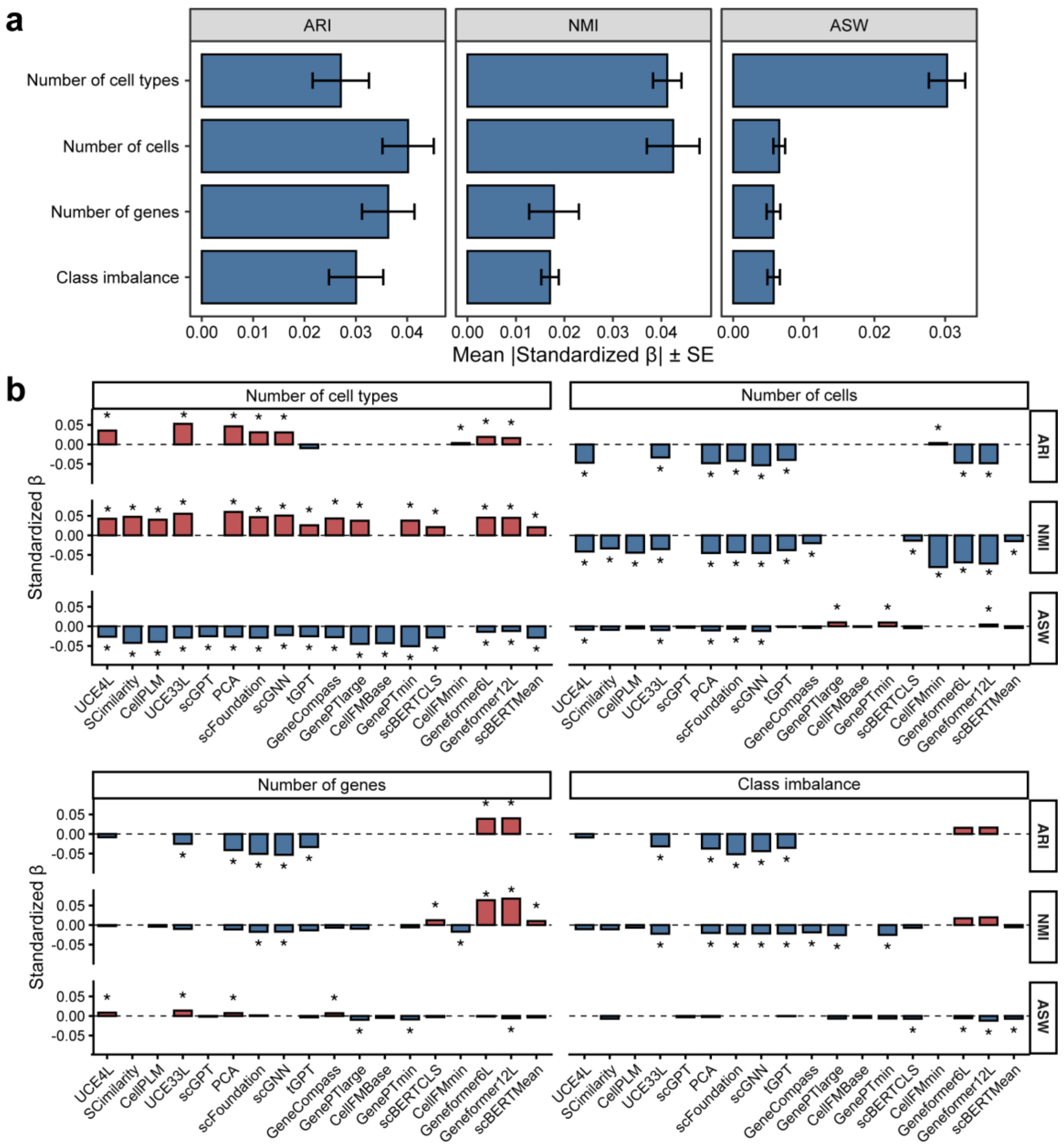


**Supplementary Fig. 6 | Regression-based driver analysis of model performance. a,** Bar plots summarize the aggregated regression coefficients for dataset features across all evaluated methods, with error bars indicating 95% confidence intervals. **b,** Regularized regression coefficients quantify the associations between dataset features and model metrics. The magnitude and direction of these coefficients reveal how specific structural data features systematically drive algorithmic performance across the evaluated methods.

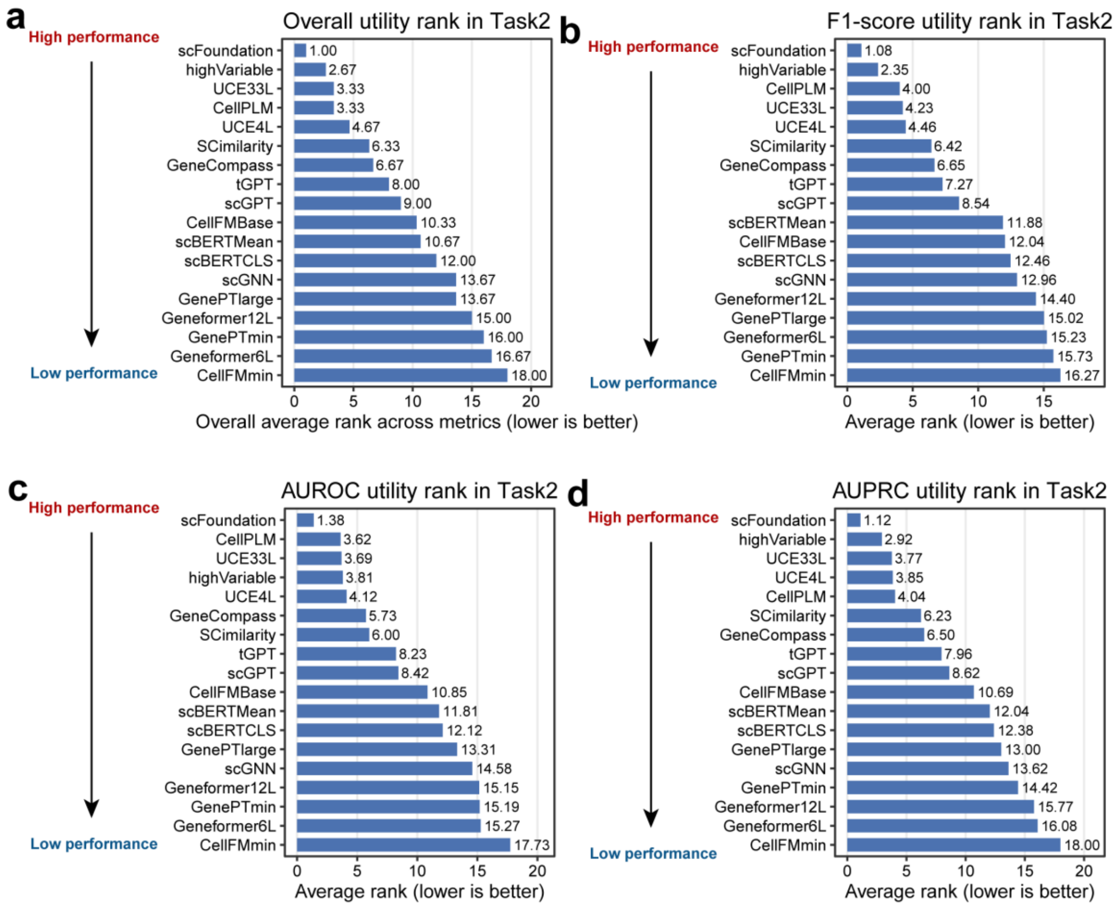


**Supplementary Fig. 7 | Average utility rankings for zero-shot cell annotation.** Bar plots display the overall aggregated rank across all three metrics **(a)**, alongside the individual average ranks across the 26 datasets for F1-score **(b)**, AUROC **(c)**, and AUPRC **(d)**. Lower numerical rank values indicate superior method performance.

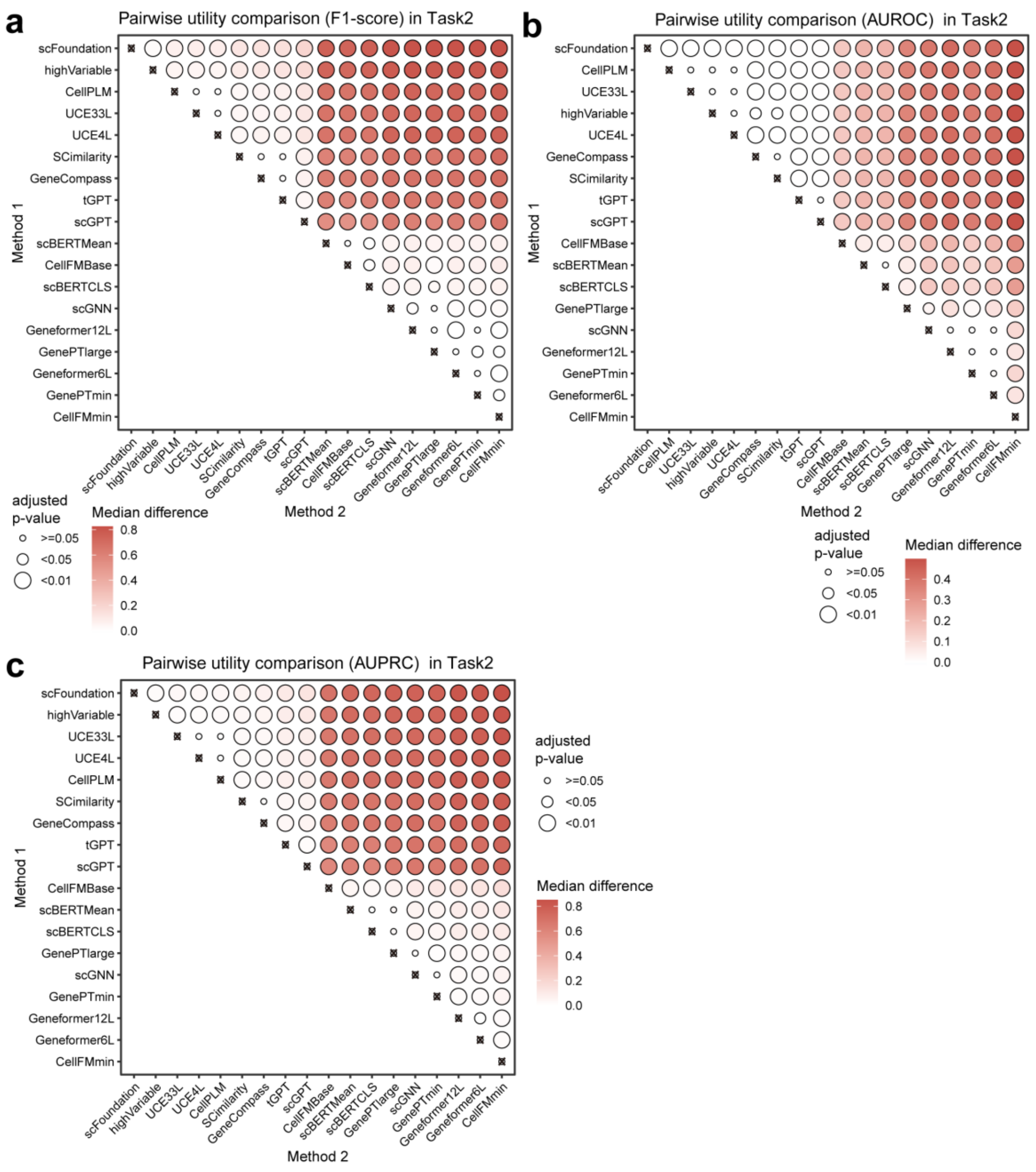


**Supplementary Fig. 8 | Pairwise statistical comparisons of zero-shot utility in cell annotation. a-c,** Bubble plots visualize the results of one-sided Wilcoxon tests comparing method performance across F1-score **(a)**, AUROC **(b)**, and AUPRC **(c)** metrics, with algorithms ordered along the axes by their independent average ranks. Bubble color intensity represents the median performance difference between the Method 1 and Method 2 algorithms, while bubble size indicates the adjusted p-value.

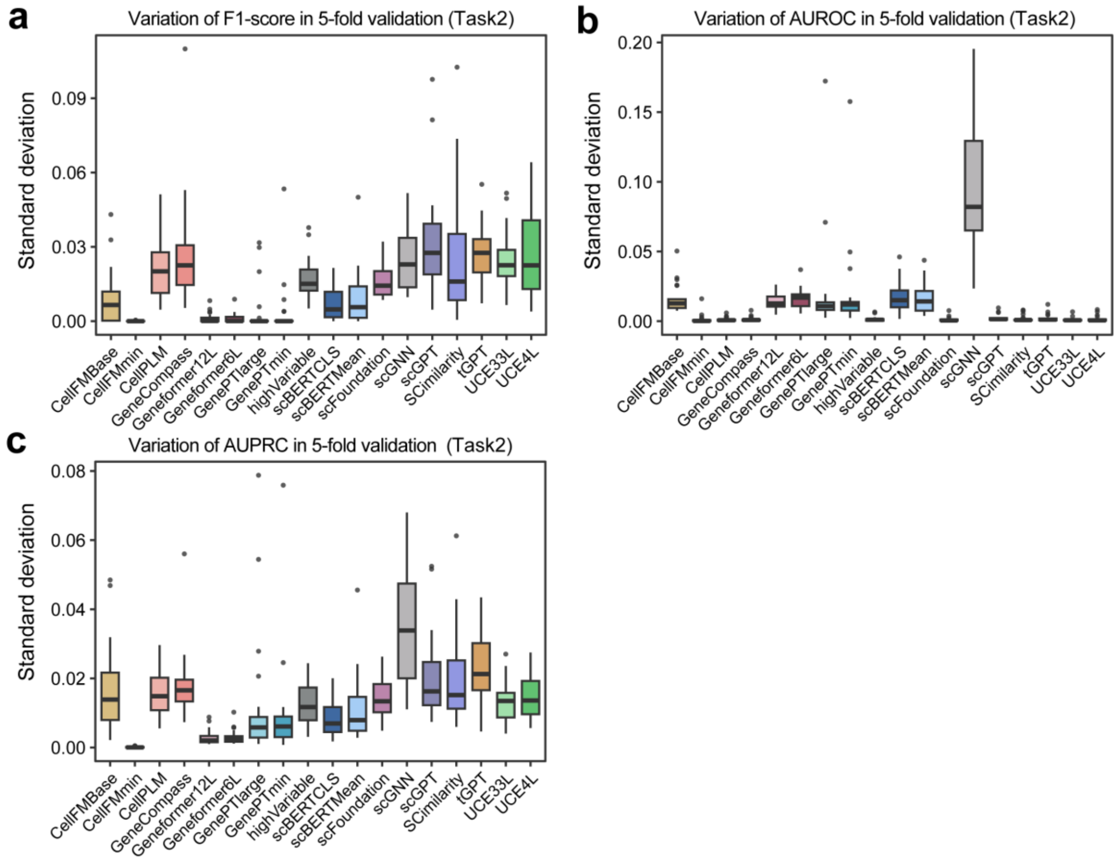


**Supplementary Fig. 9 | Standard deviation of performance metrics across 5-fold cross-validation for cell annotation.** Boxplots illustrate the variance in **(a)** F1-score, **(b)** AUROC, and **(c)** AUPRC for different algorithms. Lower standard deviation values indicate higher model stability across data folds.

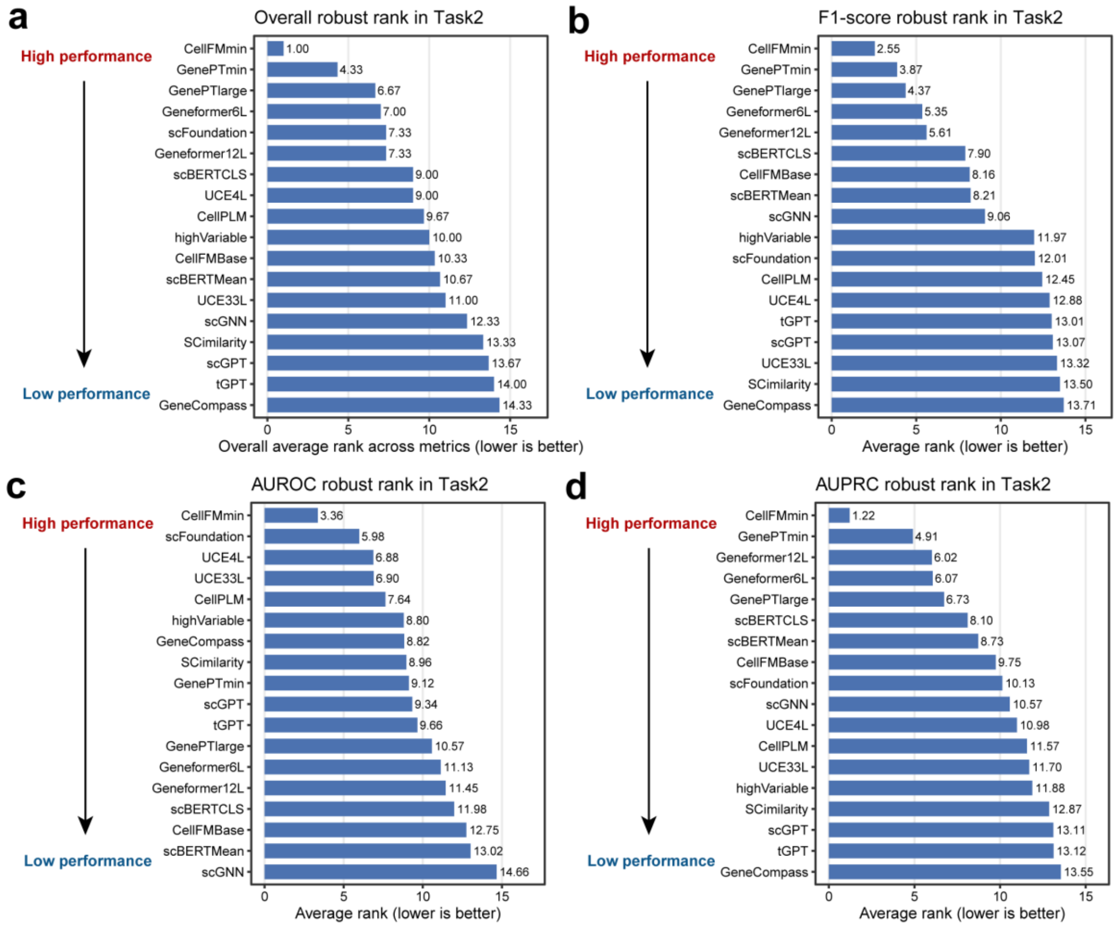


**Supplementary Fig. 10 | Average robustness rankings for zero-shot cell annotation.** Bar plots display the overall aggregated rank across all three metrics **(a)**, alongside the individual average ranks across the 328 datasets for F1-score **(b)**, AUROC **(c)**, and AUPRC **(d)**. Lower numerical rank values indicate superior robustness.

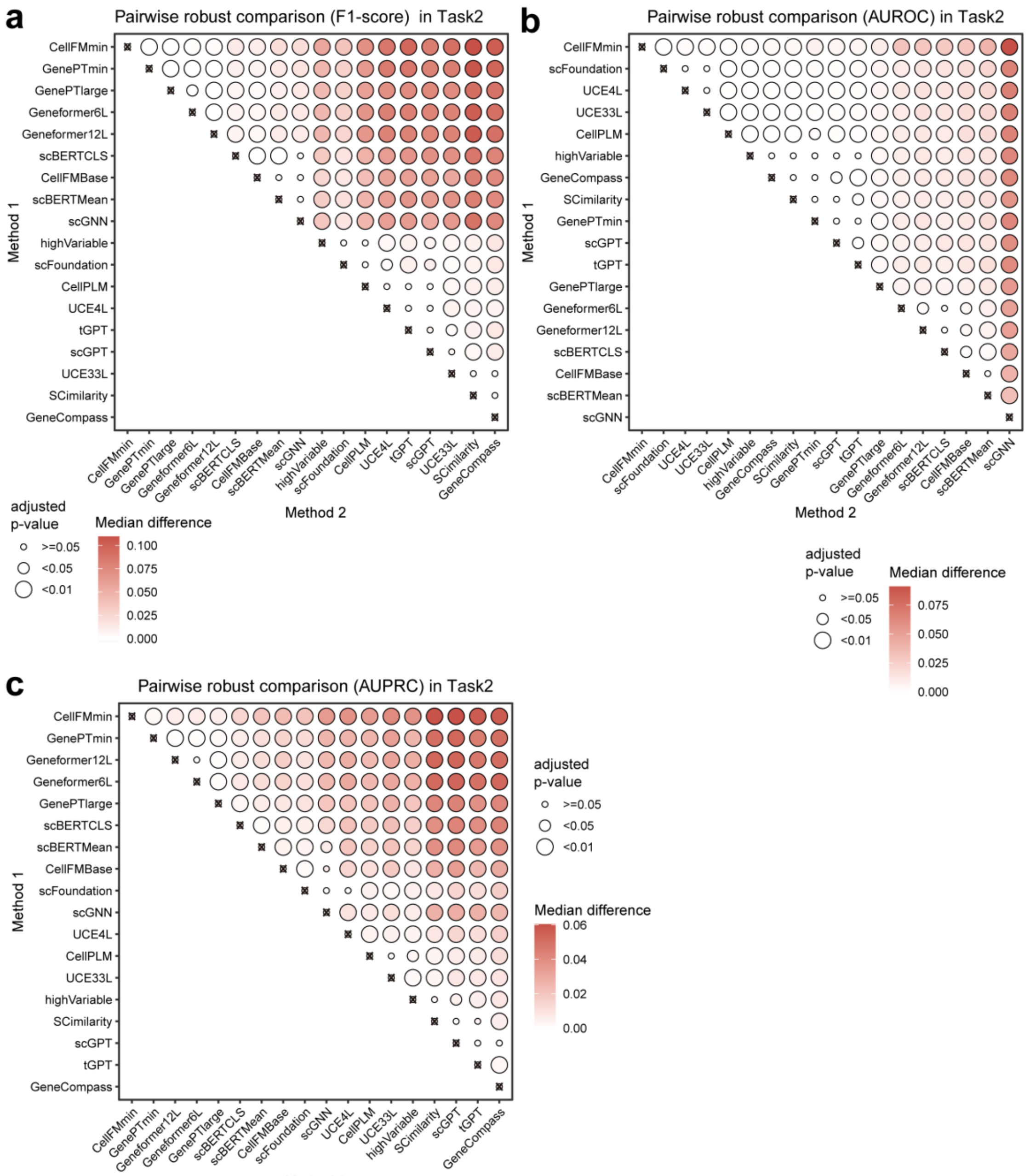


**Supplementary Fig. 11 | Pairwise statistical robustness comparisons of zero-shot cell annotation. a-c,** Bubble plots visualize the results of one-sided Wilcoxon tests comparing method robustness across F1-score **(a)**, AUROC **(b)**, and AUPRC **(c)** metrics, with algorithms ordered along the axes by their independent average ranks. Bubble color intensity represents the median robustness difference between the Method 2 and Method 1 algorithms, while bubble size indicates the adjusted p-value.

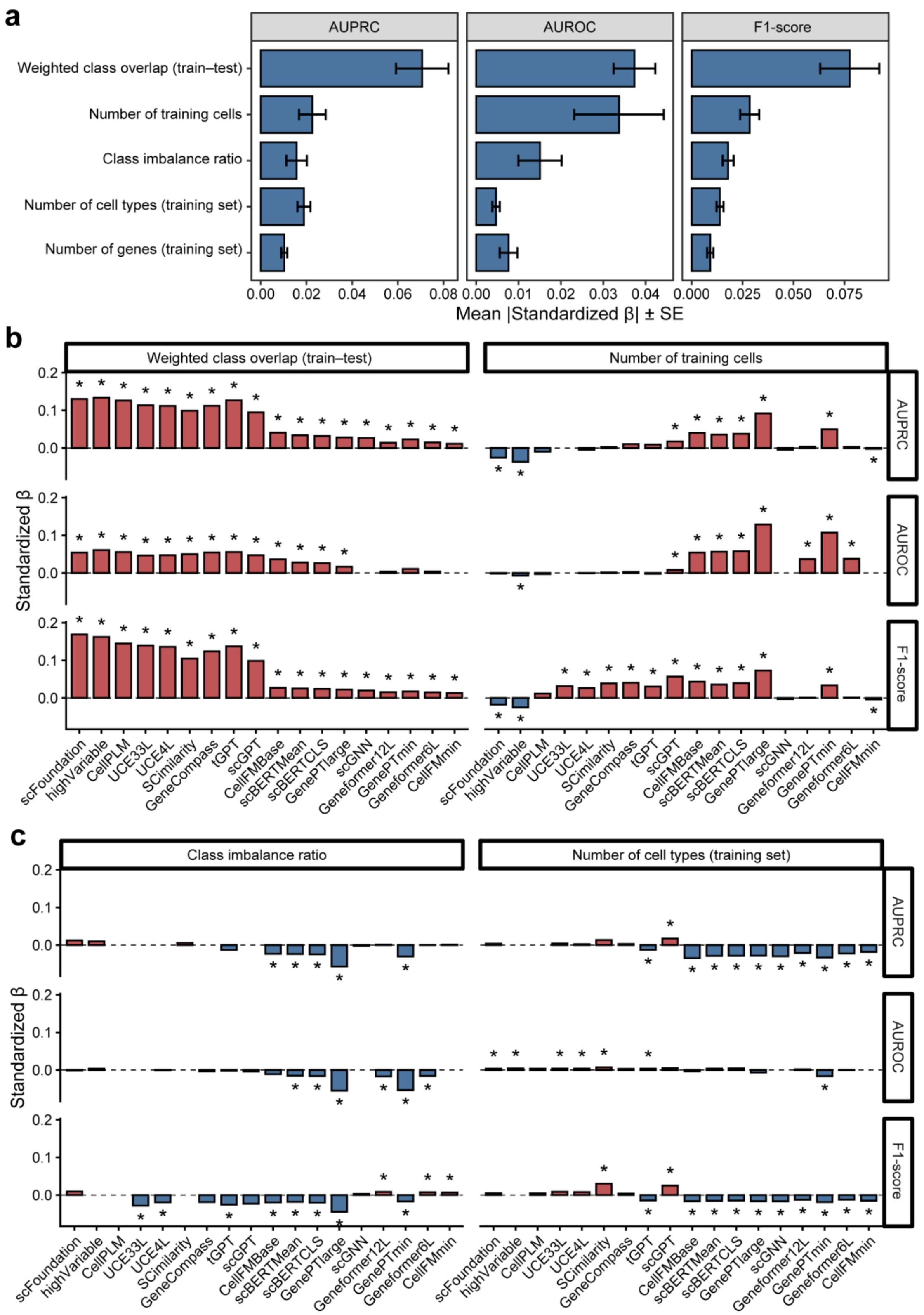


**Supplementary Fig. 12 | Regression-based driver analysis of model performance.** **a,** Bar plots summarize the aggregated regression coefficients for dataset features across all evaluated methods, with error bars indicating 95% confidence intervals. **b,** Regularized regression coefficients quantify the associations between dataset features and model metrics. The magnitude and direction of these coefficients reveal how specific structural data features systematically drive algorithmic performance across the evaluated methods.

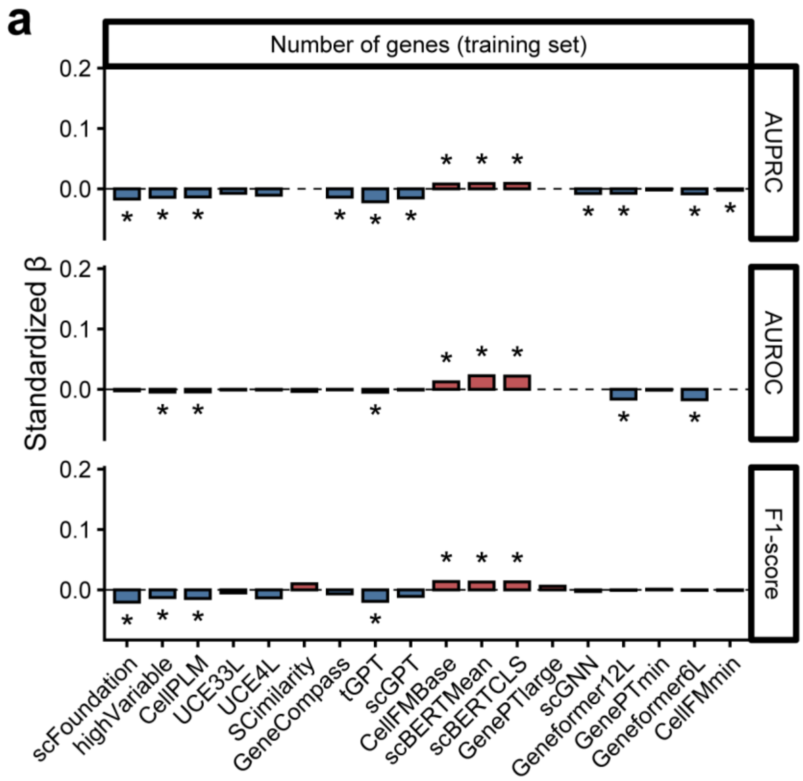


**Supplementary Fig. 13 | Regression-based driver analysis of model performance.** Regularized regression coefficients quantify the associations between dataset features and model metrics. **a,** The impact of the “Number of genes” in the training set across evaluated models.

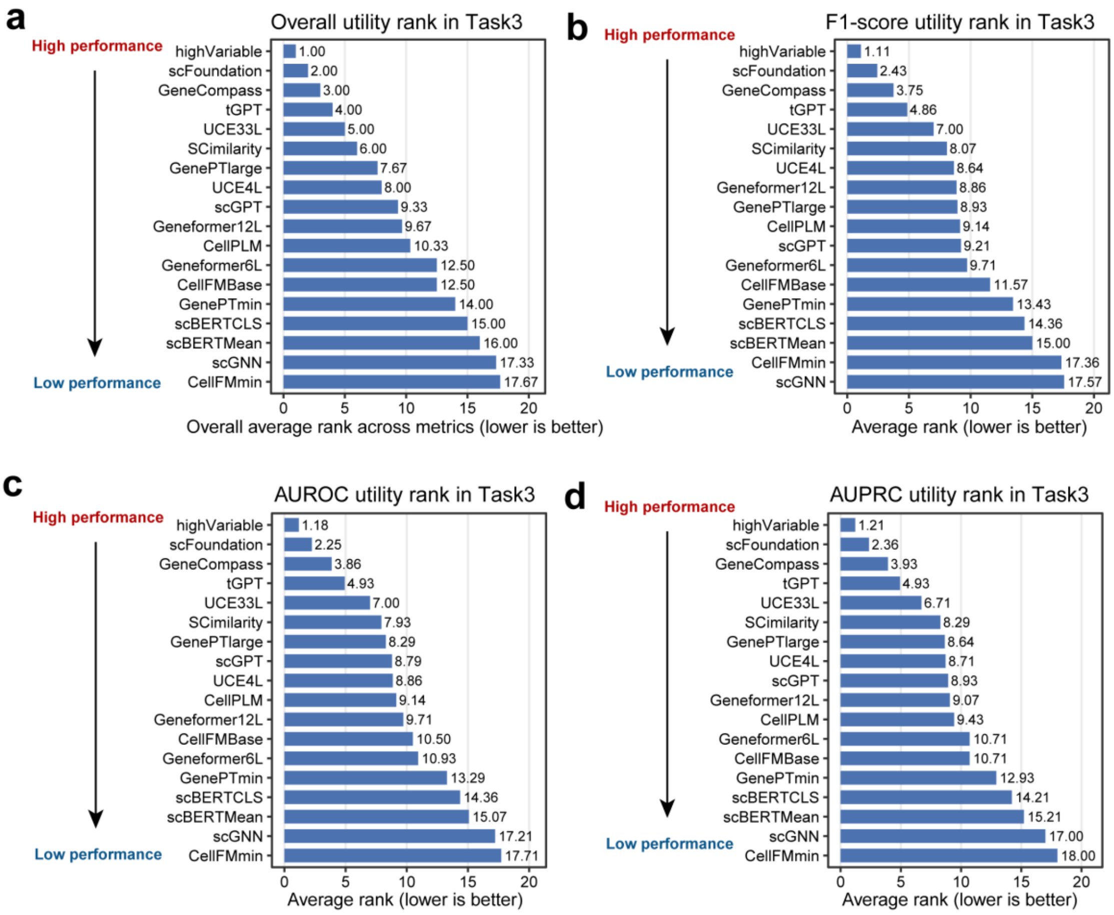


**Supplementary Fig. 14 | Average utility rankings for zero-shot drug sensitivity prediction.** Bar plots display the overall aggregated rank across all three metrics **(a)**, alongside the individual average ranks across the 14 datasets for F1-score **(b)**, AUROC **(c)**, and AUPRC **(d)**. Lower numerical rank values indicate superior method performance.

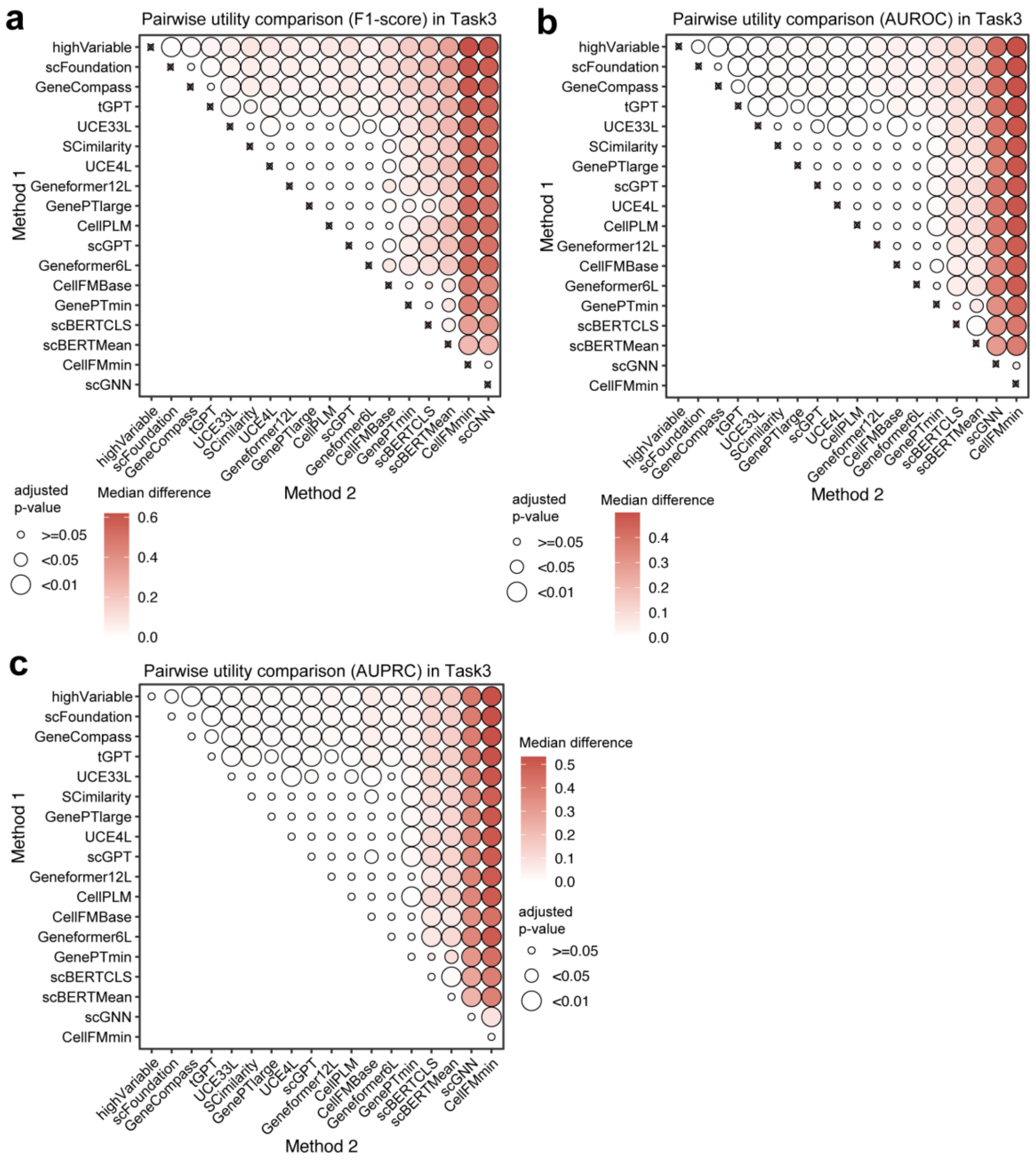


**Supplementary Fig. 15 | Pairwise statistical comparisons of zero-shot utility in drug sensitivity prediction. a-c,** Bubble plots visualize the results of one-sided Wilcoxon tests comparing method performance across F1-score **(a)**, AUROC **(b)**, and AUPRC **(c)** metrics, with algorithms ordered along the axes by their independent average ranks. Bubble color intensity represents the median performance difference between the Method 1 and Method 2 algorithms, while bubble size indicates the adjusted p-value.

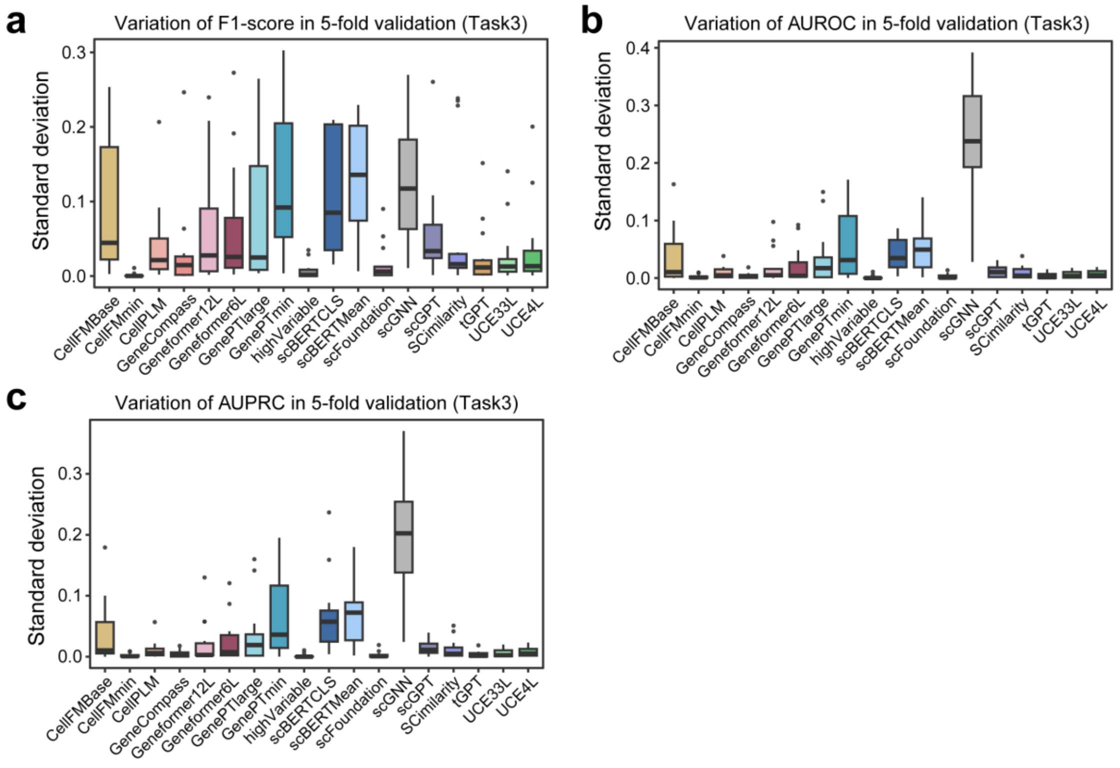


**Supplementary Fig. 16 | Standard deviation of performance metrics across 5-fold cross-validation for drug sensitivity prediction.** Boxplots illustrate the variance in **(a)** F1-score, **(b)** AUROC, and **(c)** AUPRC for different algorithms. Lower standard deviation values indicate higher model stability across data folds.

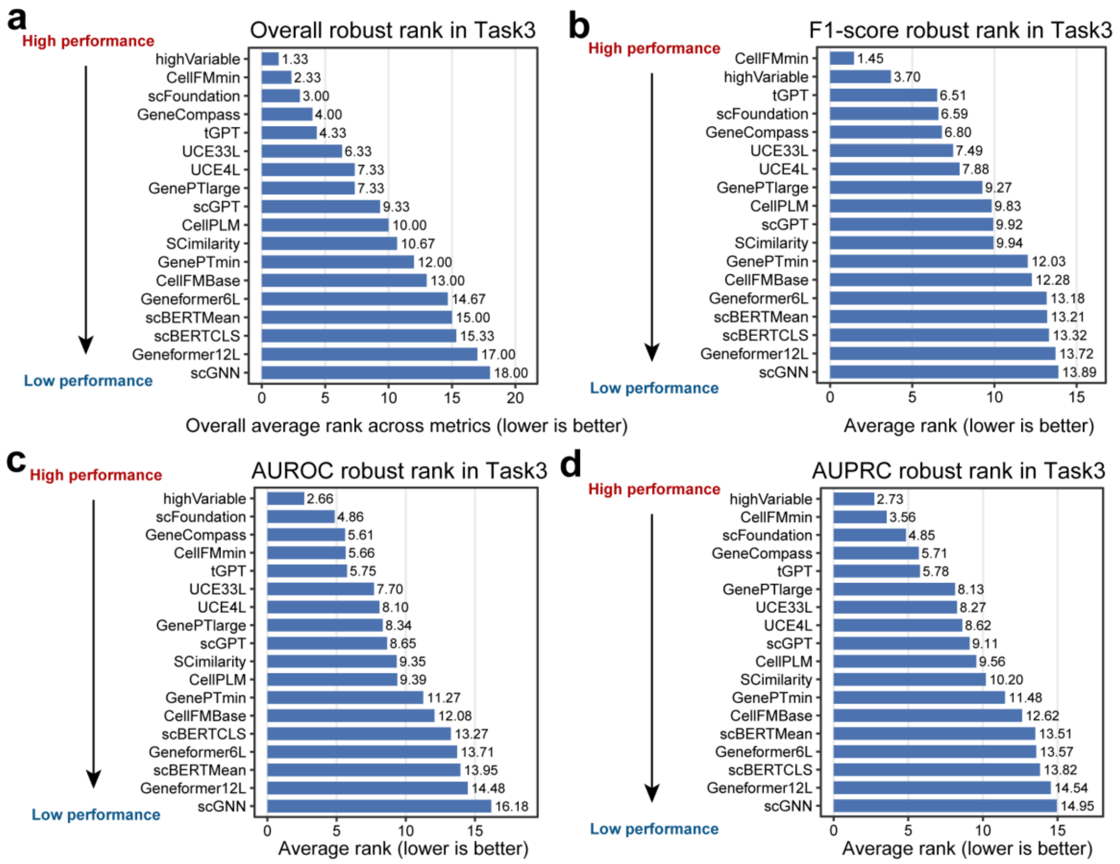


**Supplementary Fig. 17 | Average robustness rankings for zero-shot drug sensitivity prediction.** Bar plots display the overall aggregated rank across all three metrics **(a)**, alongside the individual average ranks across the 79 datasets for F1-score **(b)**, AUROC **(c)**, and AUPRC **(d)**. Lower numerical rank values indicate superior robustness.

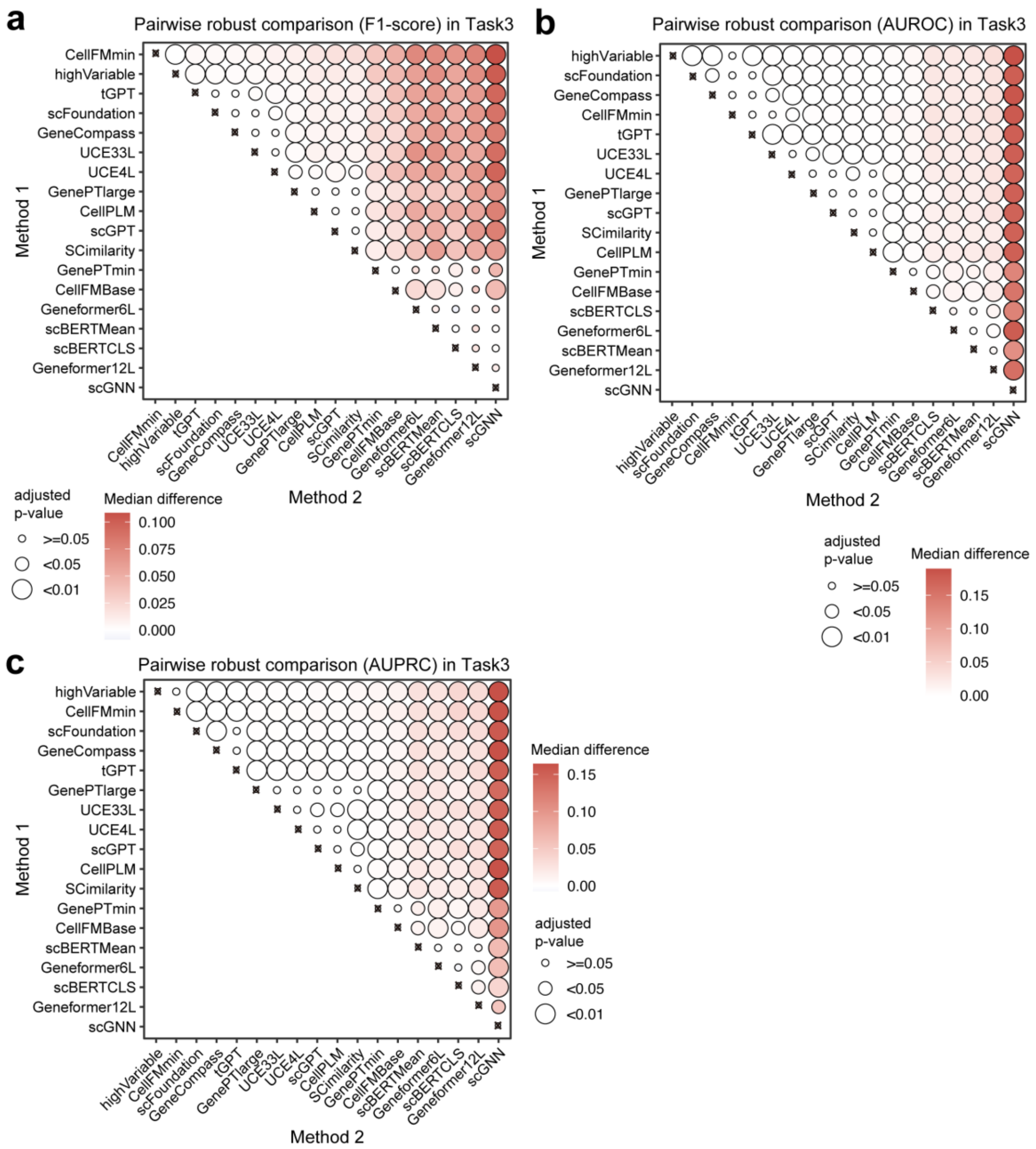


**Supplementary Fig. 18 | Pairwise statistical robustness comparisons of zero-shot drug sensitivity prediction. a-c,** Bubble plots visualize the results of one-sided Wilcoxon tests comparing method robustness across F1-score **(a)**, AUROC **(b)**, and AUPRC **(c)** metrics, with algorithms ordered along the axes by their independent average ranks. Bubble color intensity represents the median robustness difference between the Method 2 and Method 1 algorithms, while bubble size indicates the adjusted p-value.

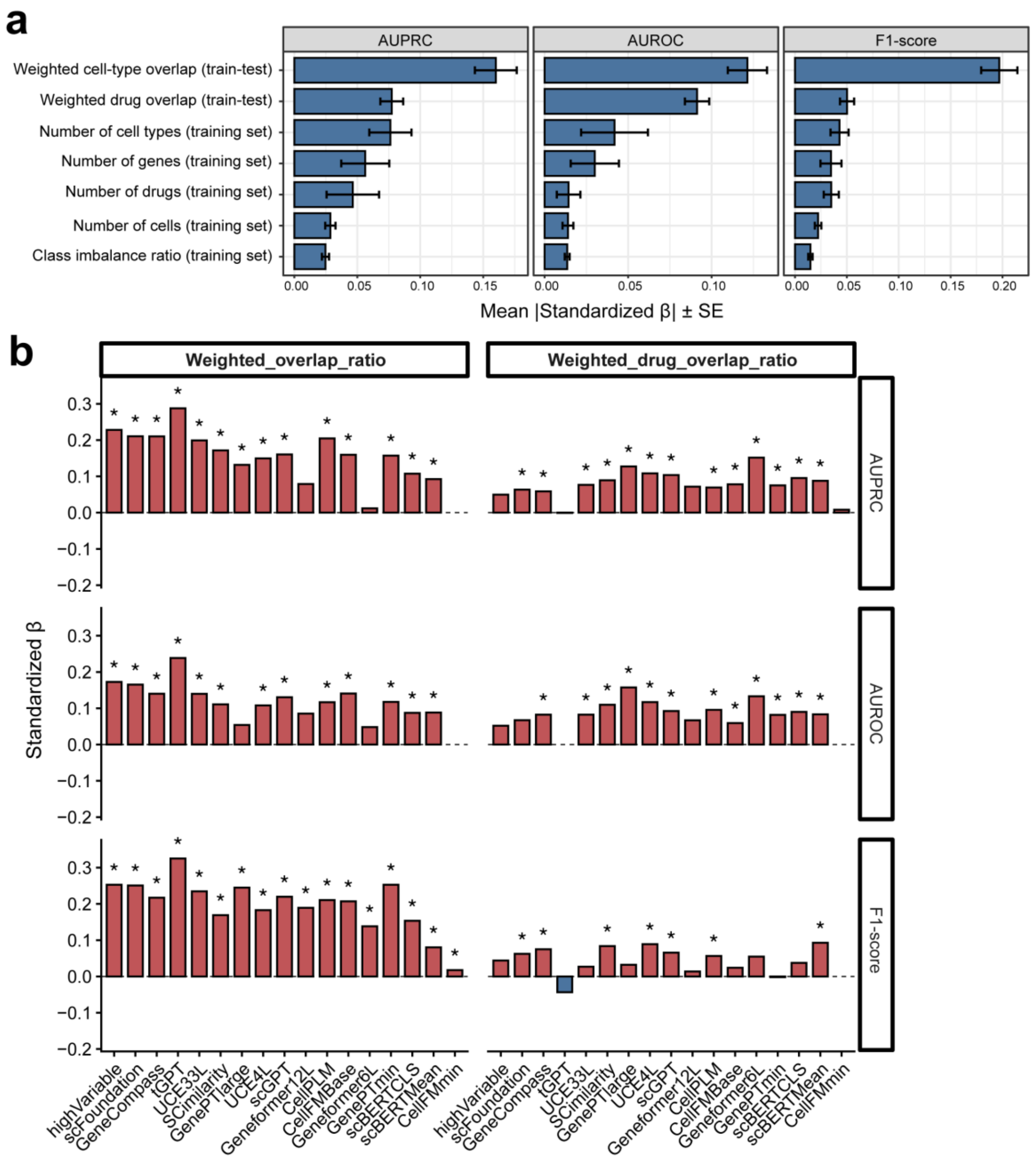


**Supplementary Fig. 19 | Regression-based driver analysis of model performance.** **a,** Bar plots summarize the aggregated regression coefficients for dataset features across all evaluated methods, with error bars indicating 95% confidence intervals. **b,** Regularized regression coefficients quantify the associations between dataset features and model metrics. The magnitude and direction of these coefficients reveal how specific structural data features systematically drive algorithmic performance across the evaluated methods.

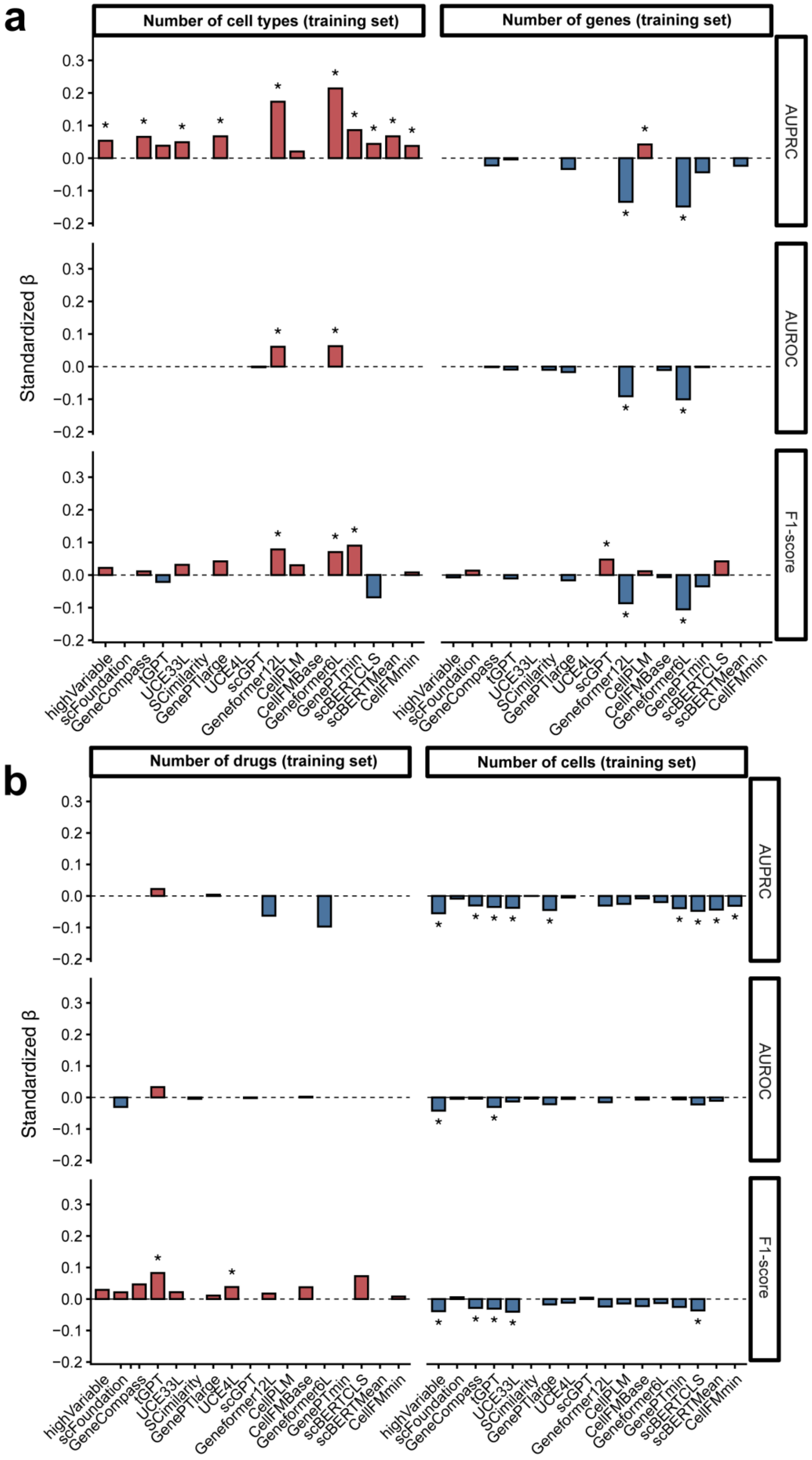

a
Number of cell types (training set)
Number of genes (training set)
AUPRC
AUROC
F1-score
Standardized β
highVariable
scFoundation
GeneCompass
tGPT
UCE33L
SCimilarity
GenePTlarge
UCE4L
scGPT
Geneformer12L
CellPLM
CellFMBase
Geneformer6L
GenePTmin
scBERTCLS
scBERTMean
CellFMmin
b
Number of drugs (training set)
Number of cells (training set)

**Supplementary Fig. 20 | Regression-based driver analysis of model performance.** Regularized regression coefficients quantify the associations between dataset features and model metrics. **a,** The impact of the “Number of cell types” and “Number of genes” in the training set across evaluated models. **b,** The impact of the “Number of drugs” and the “Number of cells” in the training set.

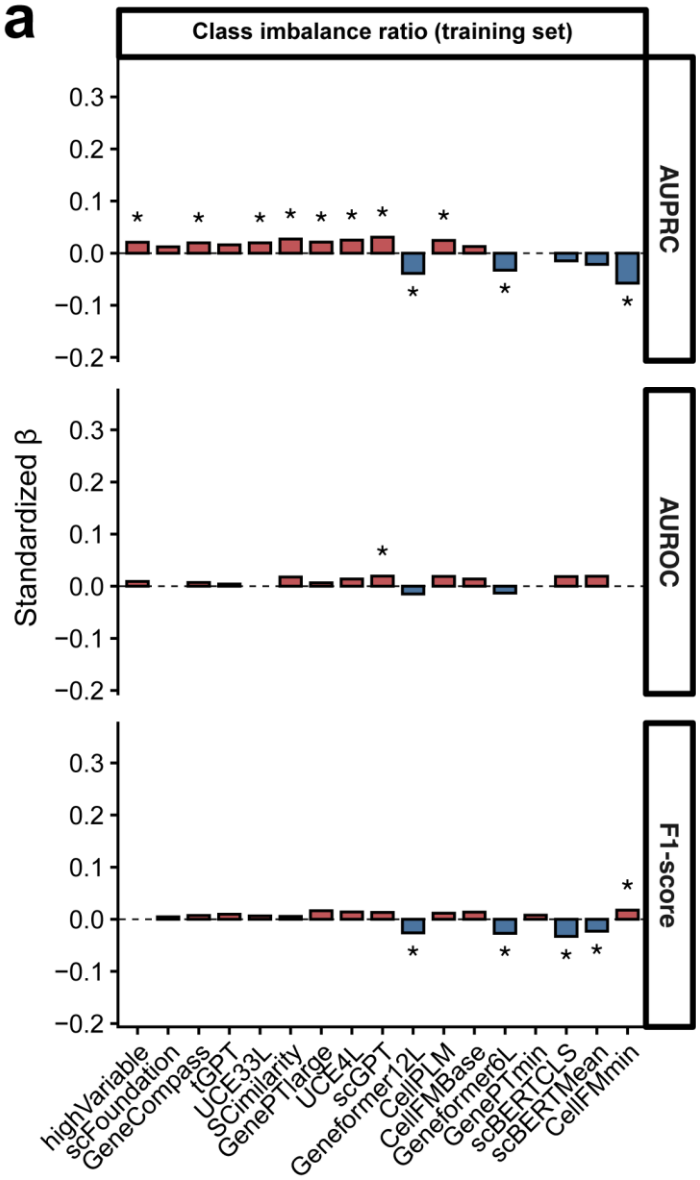


**Supplementary Fig. 21 | Regression-based driver analysis of model performance.** Regularized regression coefficients quantify the associations between dataset features and model metrics. **a,** The impact of the “Class imbalance ratio” in the training set across evaluated models.

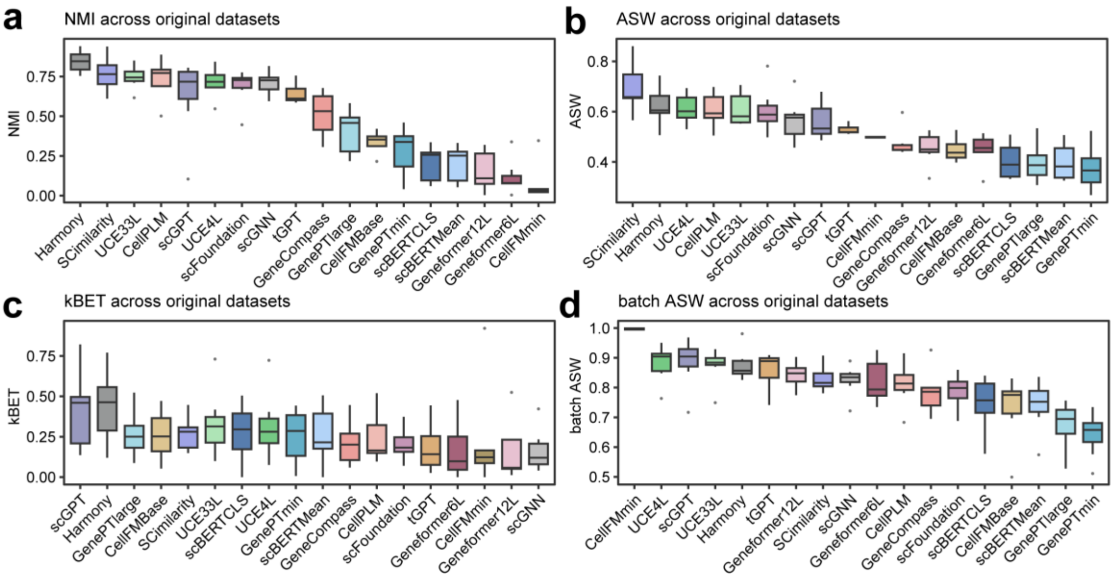


**Supplementary Fig. 22 | Performance comparison of evaluated methods across original datasets in batch integration.** Boxplots showing performance distributions of 18 methods across original datasets measured by NMI (a), ASW (b), kBET (c), and batch ASW (d), respectively. For each metric, models are ranked along the x-axis in descending order.

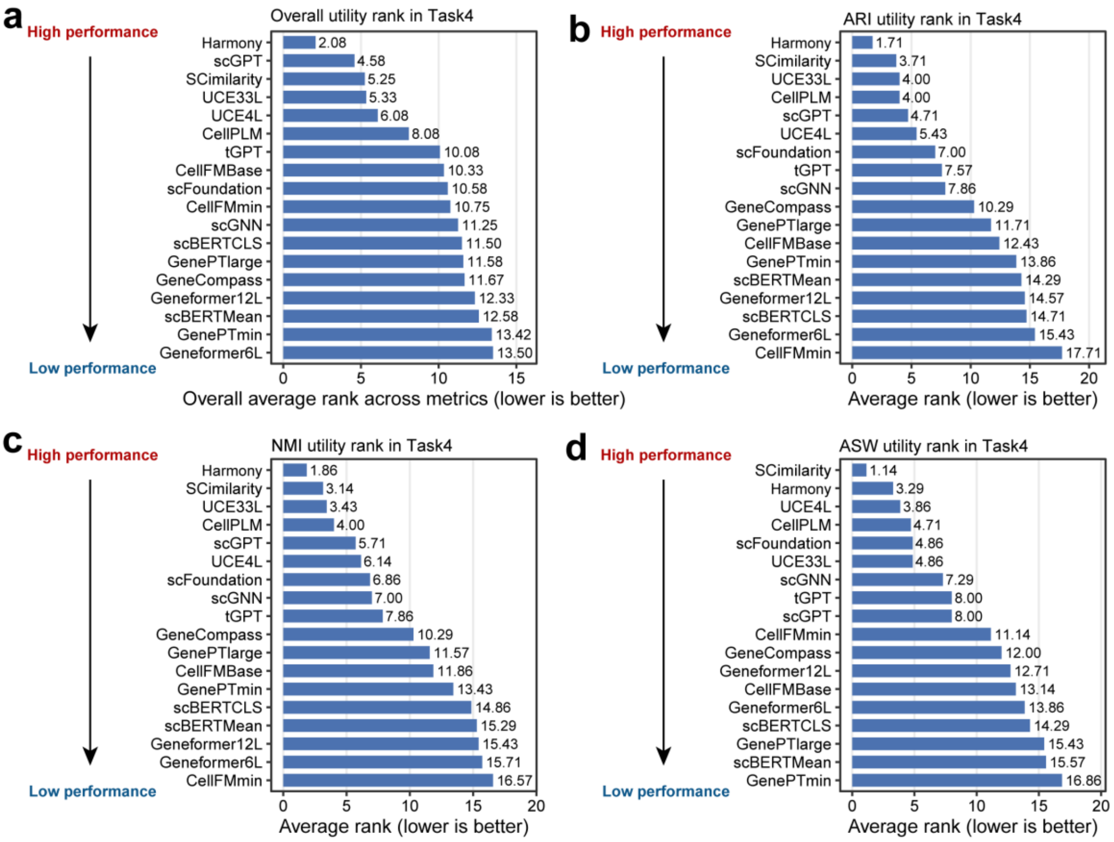

**Supplementary Fig. 23 | Average utility rankings for zero-shot batch integration.** Bar plots display the overall aggregated rank across all six metrics **(a)**, alongside the individual average ranks across the 7 datasets for ARI **(b)**, NMI **(c)**, and ASW **(d)**. Lower numerical rank values indicate superior method performance.

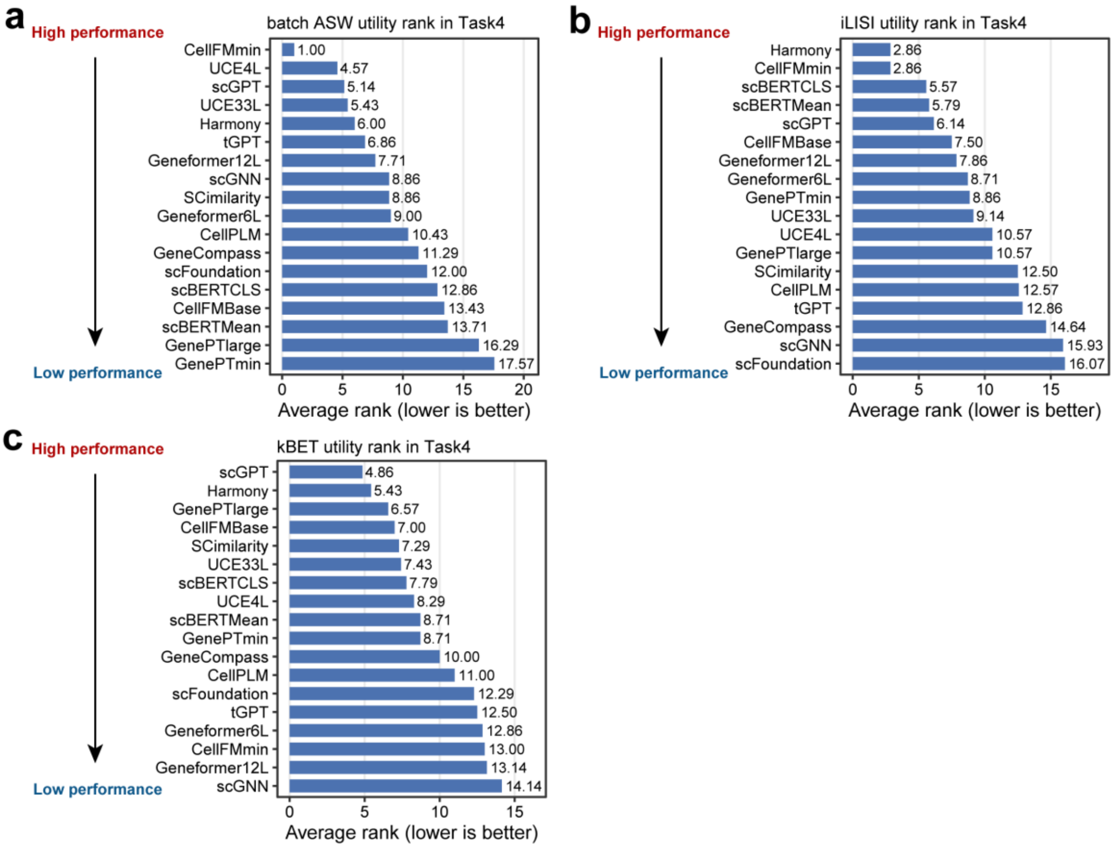


**Supplementary Fig. 24 | Average utility rankings for zero-shot batch integration.** Bar plots display the individual average ranks across the 7 datasets for batch ASW **(a)**, iLISI **(b)**, and kBET **(c)**. Lower numerical rank values indicate superior method performance.

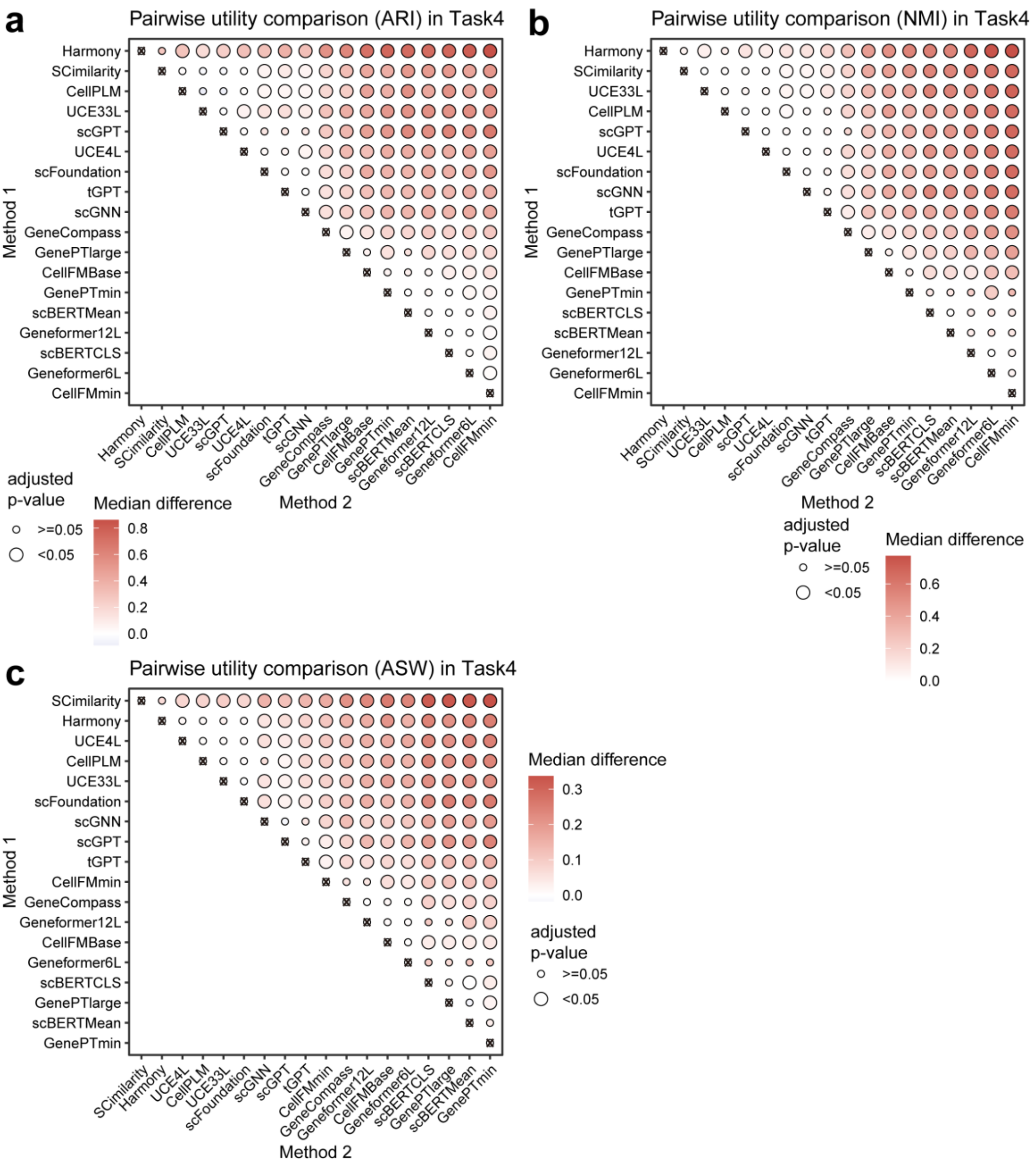


**Supplementary Fig. 25 | Pairwise statistical comparisons of zero-shot utility in batch integration. a-c,** Bubble plots visualize the results of one-sided Wilcoxon tests comparing method performance across ARI **(a)**, NMI **(b)**, and ASW **(c)** metrics, with algorithms ordered along the axes by their independent average ranks. Bubble color intensity represents the median performance difference between the Method 1 and Method 2 algorithms, while bubble size indicates the adjusted p-value.

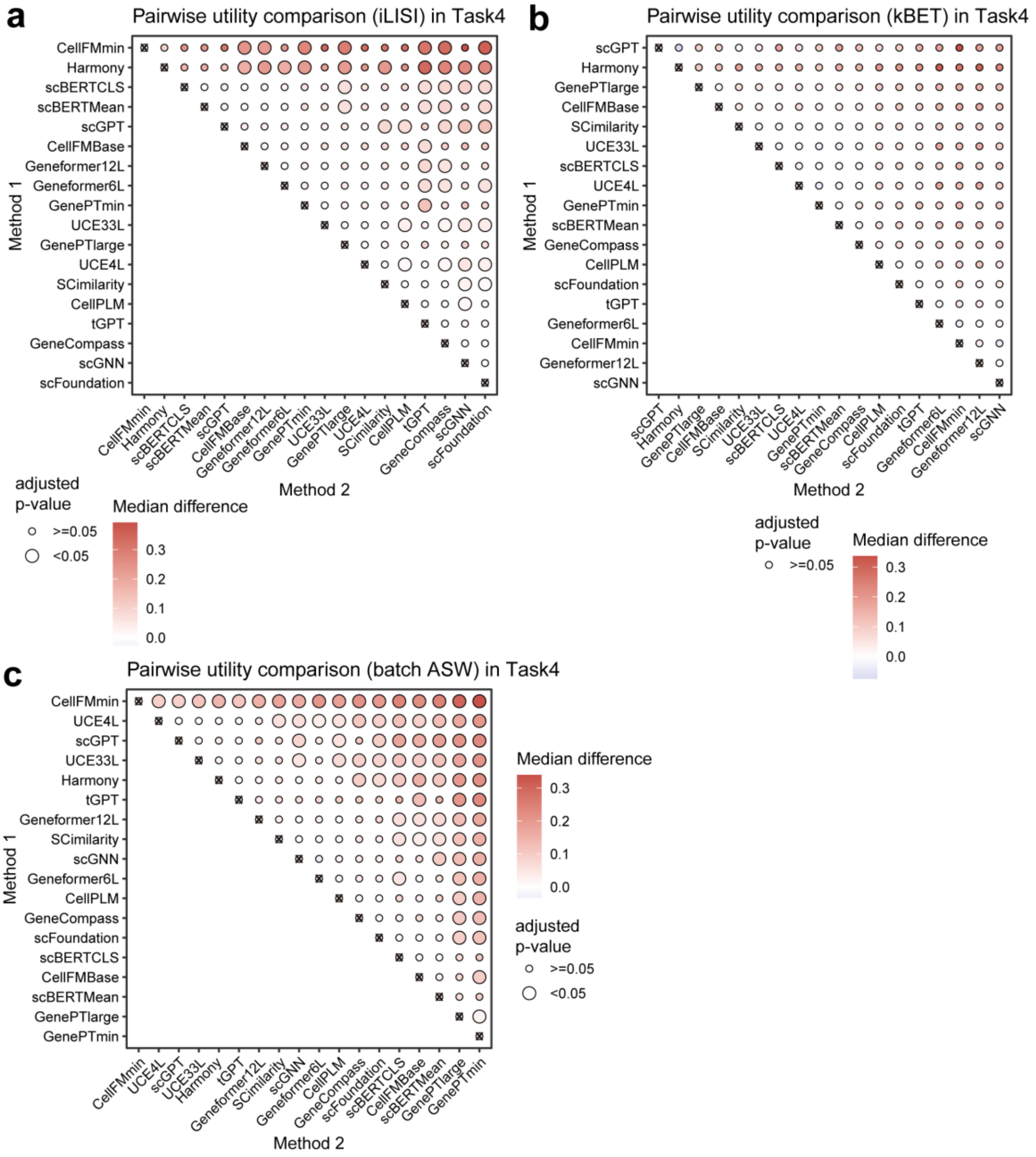


**Supplementary Fig. 26 | Pairwise statistical comparisons of zero-shot utility in batch integration. a-c,** Bubble plots visualize the results of one-sided Wilcoxon tests comparing method performance across iLISI **(a)**, kBET **(b)**, and batch ASW **(c)** metrics, with algorithms ordered along the axes by their independent average ranks. Bubble color intensity represents the median performance difference between the Method 1 and Method 2 algorithms, while bubble size indicates the adjusted p-value.

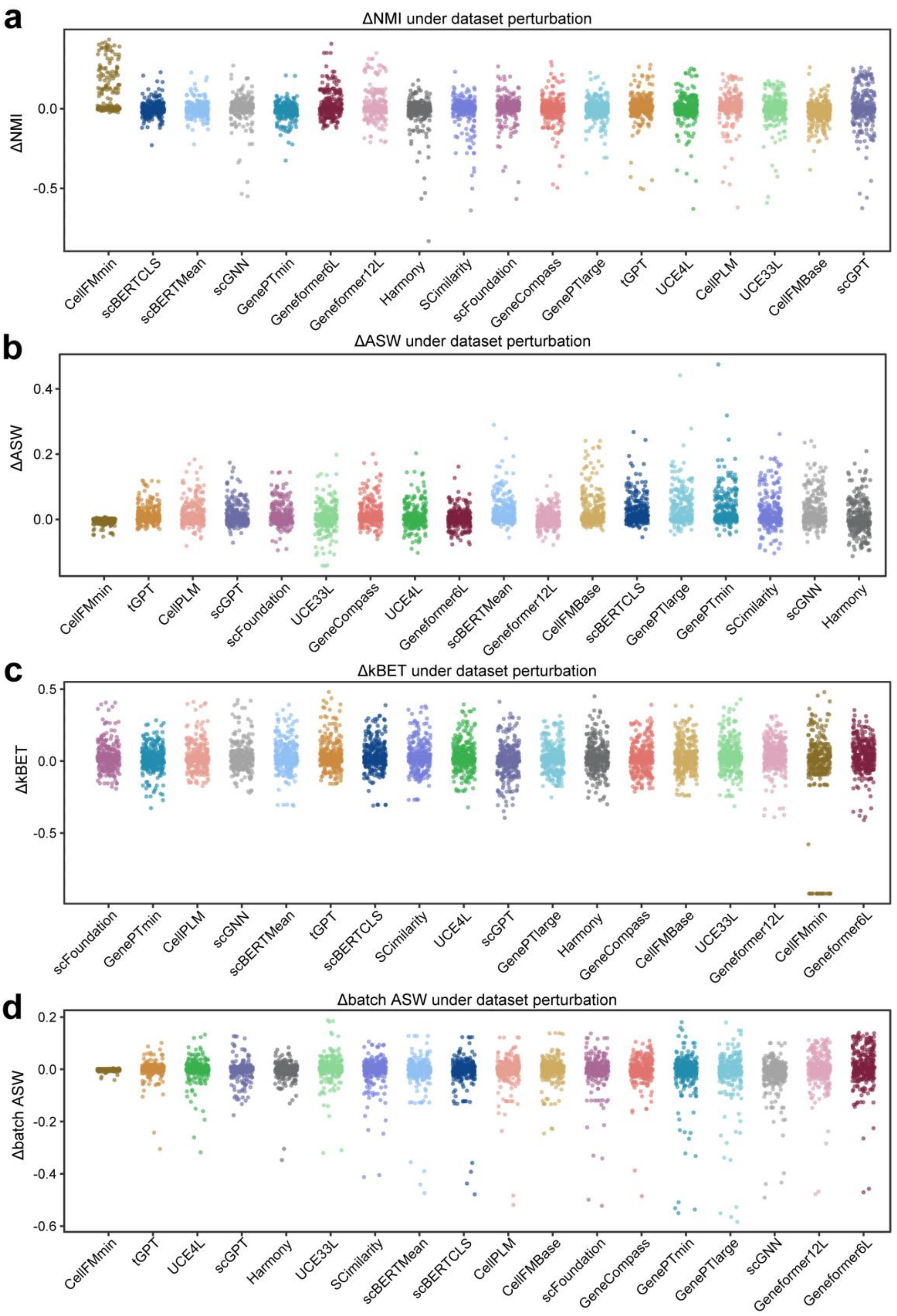


**Supplementary Fig. 27 | Robust of evaluated methods in batch integration.** Scatter plots showing performance deviations under perturbation-derived datasets for NMI **(a),**

ASW **(b),** kBET **(c),** and batch ASW **(d),** respectively. Each point represents one perturbation instance.

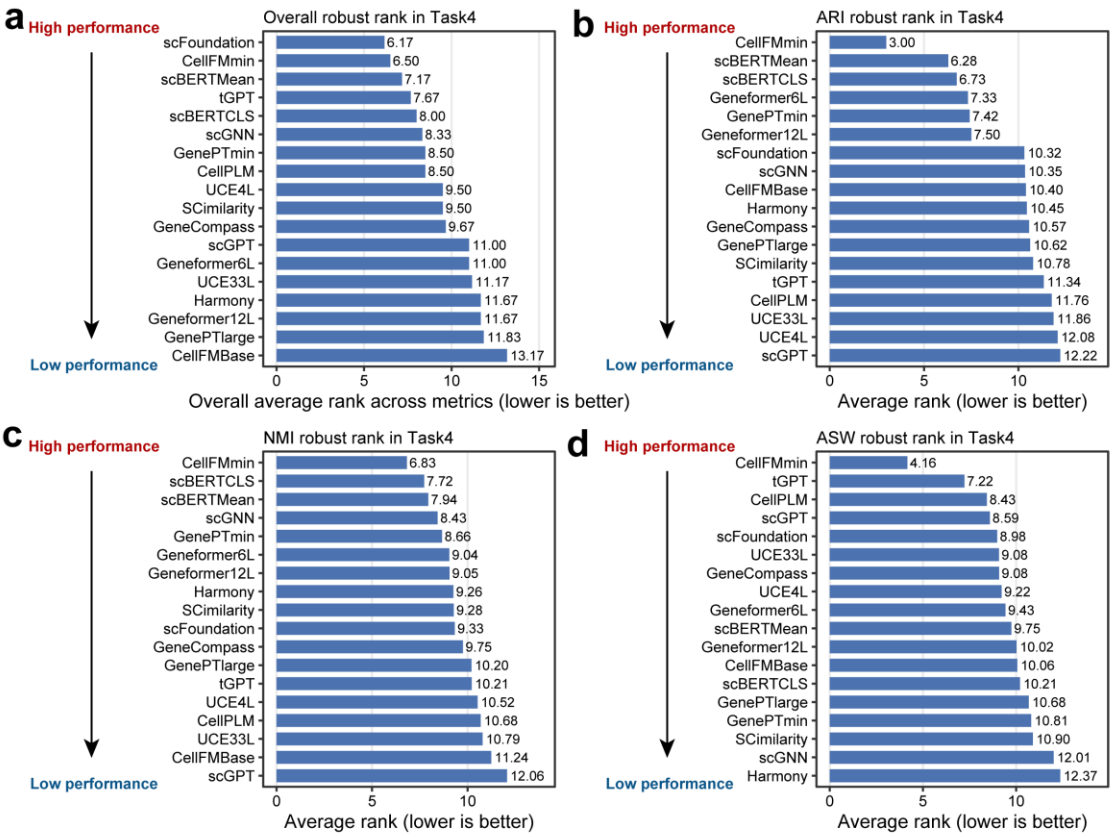


**Supplementary Fig. 28 | Average robustness rankings for zero-shot batch integration.** Bar plots display the overall aggregated rank across all six metrics **(a)**, alongside the individual average ranks across the 325 datasets for ARI **(b)**, NMI **(c)**, and ASW **(d)**. Lower numerical rank values indicate superior robustness.

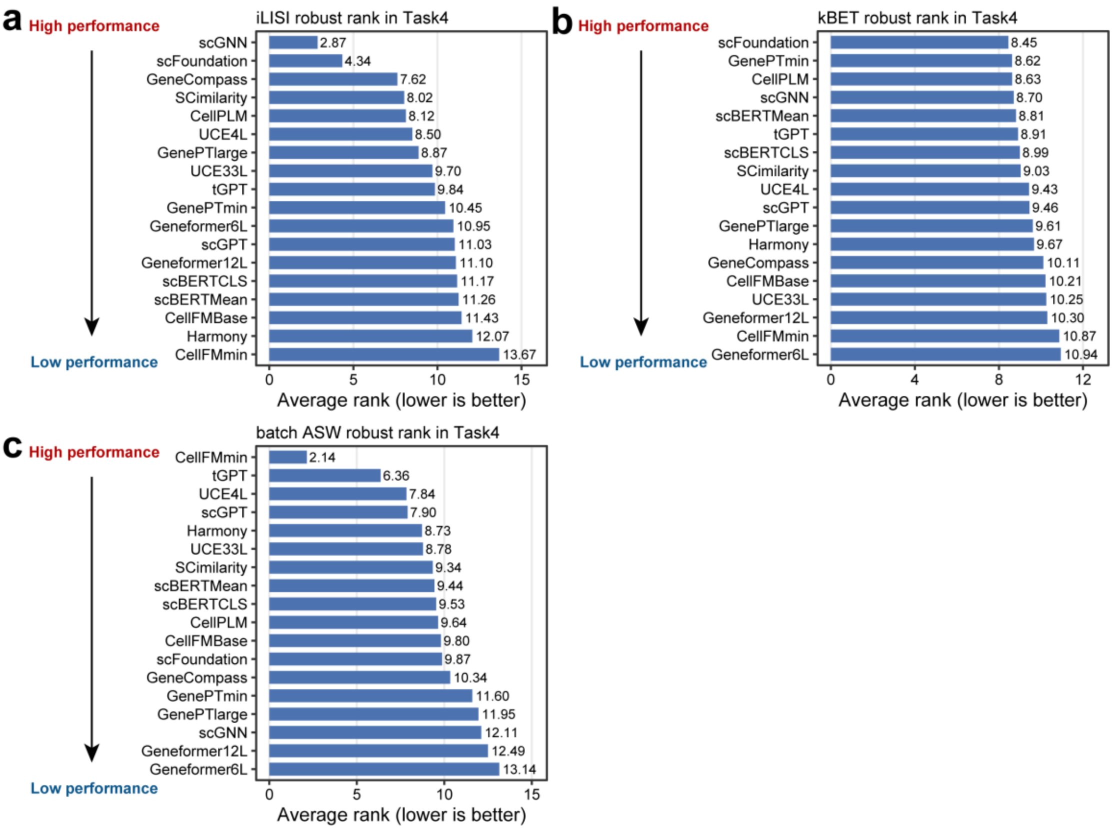


**Supplementary Fig. 29 | Average robustness rankings for zero-shot batch integration.** Bar plots display the individual average ranks across the 325 datasets for iLISI **(b)**, kBET **(c)**, and batch ASW **(d)**. Lower numerical rank values indicate superior robustness.

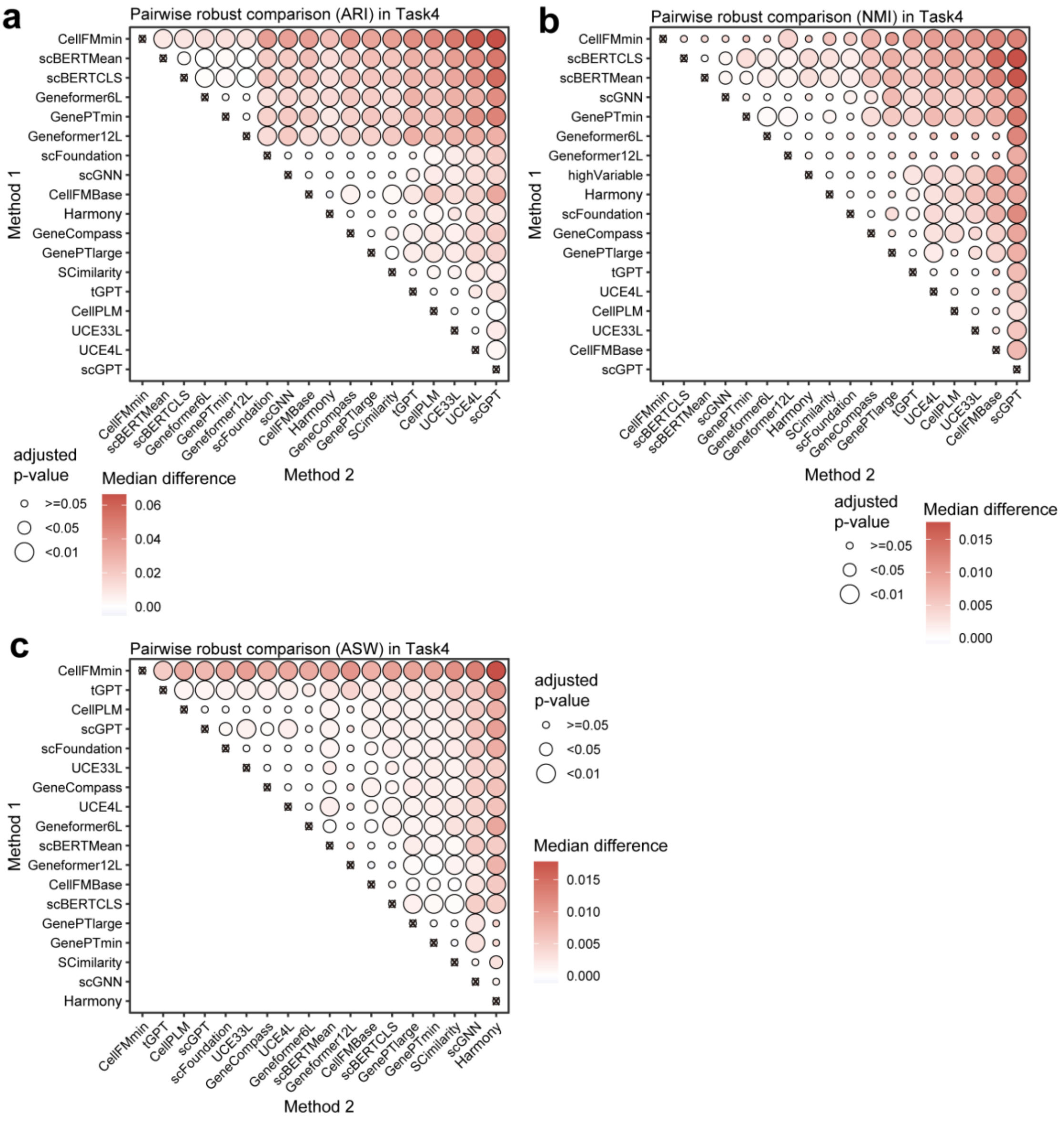


**Supplementary Fig. 30 | Pairwise statistical robustness comparisons of zero-shot batch integration. a-c,** Bubble plots visualize the results of one-sided Wilcoxon tests comparing method robustness across ARI **(a)**, NMI **(b)**, and ASW **(c)** metrics, with algorithms ordered along the axes by their independent average ranks. Bubble color intensity represents the median robustness difference between the Method 2 and Method 1 algorithms, while bubble size indicates the adjusted p-value.

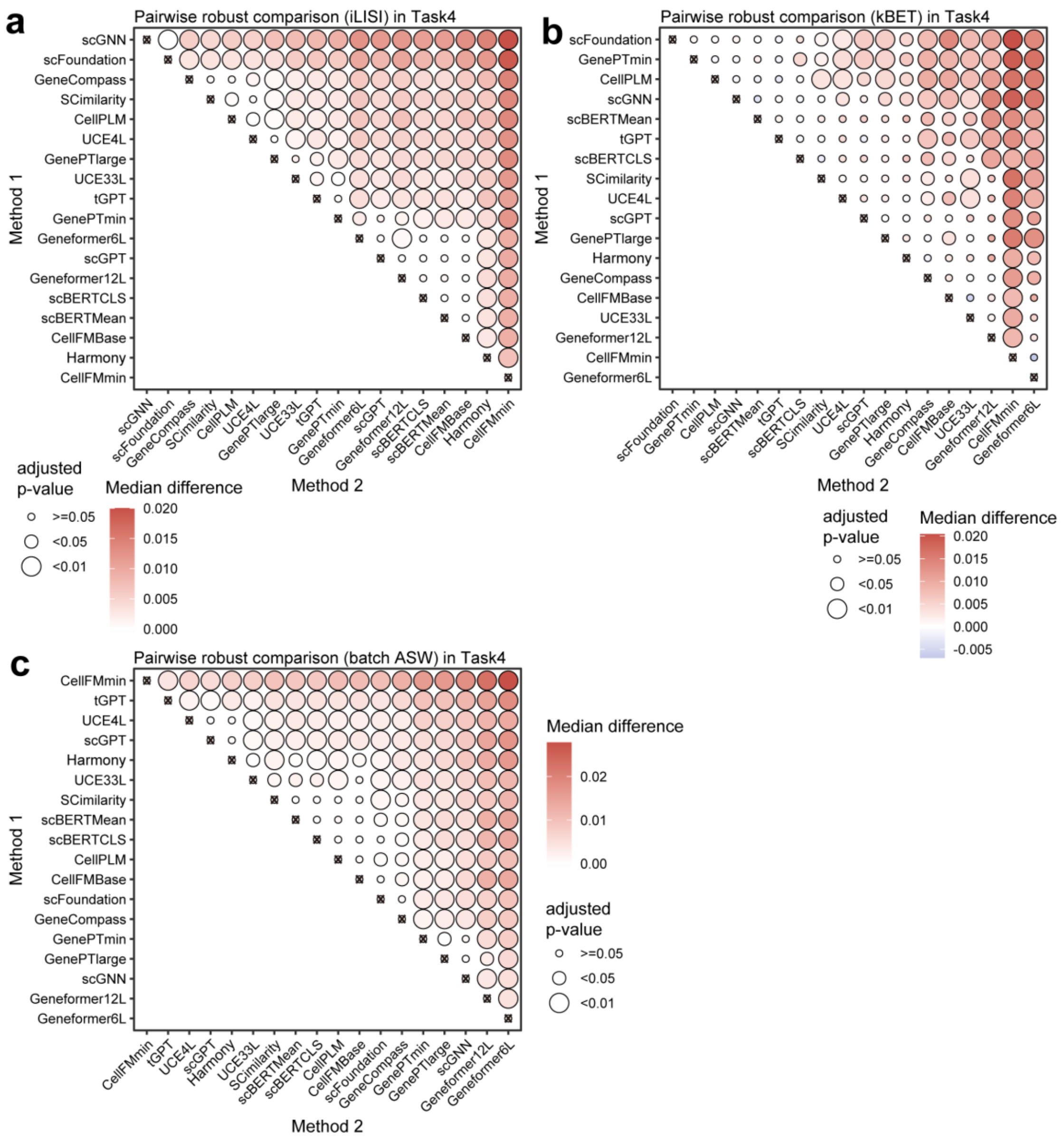


**Supplementary Fig. 31 | Pairwise statistical robustness comparisons of zero-shot batch integration. a-c,** Bubble plots visualize the results of one-sided Wilcoxon tests comparing method robustness across iLISI **(a)**, kBET **(b)**, and batch ASW **(c)** metrics, with algorithms ordered along the axes by their independent average ranks. Bubble color intensity represents the median robustness difference between the Method 2 and Method 1 algorithms, while bubble size indicates the adjusted p-value.

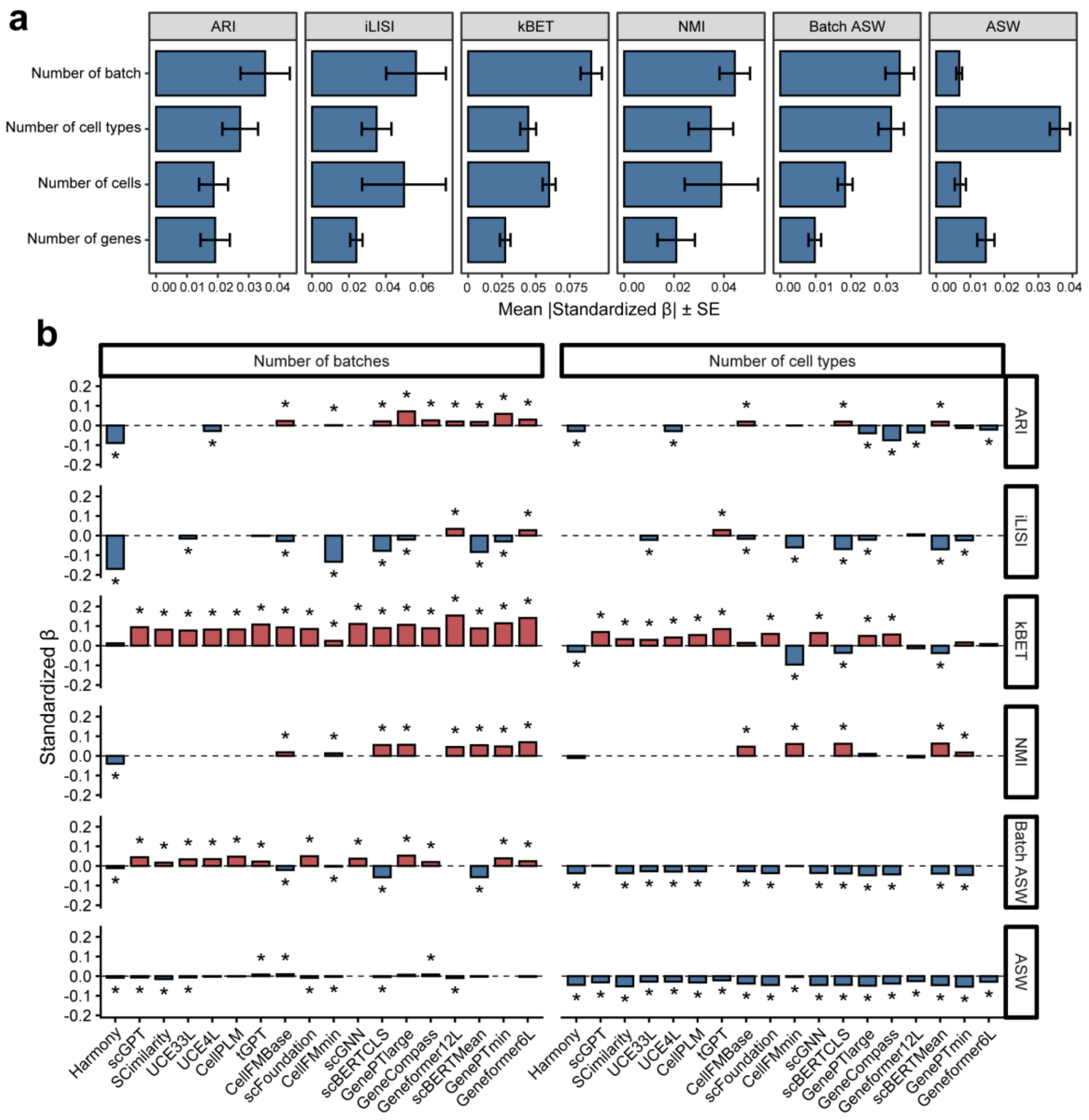


**Supplementary Fig. 32 | Regression-based driver analysis of model performance.** **a,** Bar plots summarize the aggregated regression coefficients for dataset features across all evaluated methods, with error bars indicating 95% confidence intervals. **b,** Regularized regression coefficients quantify the associations between dataset features and model metrics. The magnitude and direction of these coefficients reveal how specific structural data features systematically drive algorithmic performance across the evaluated methods.

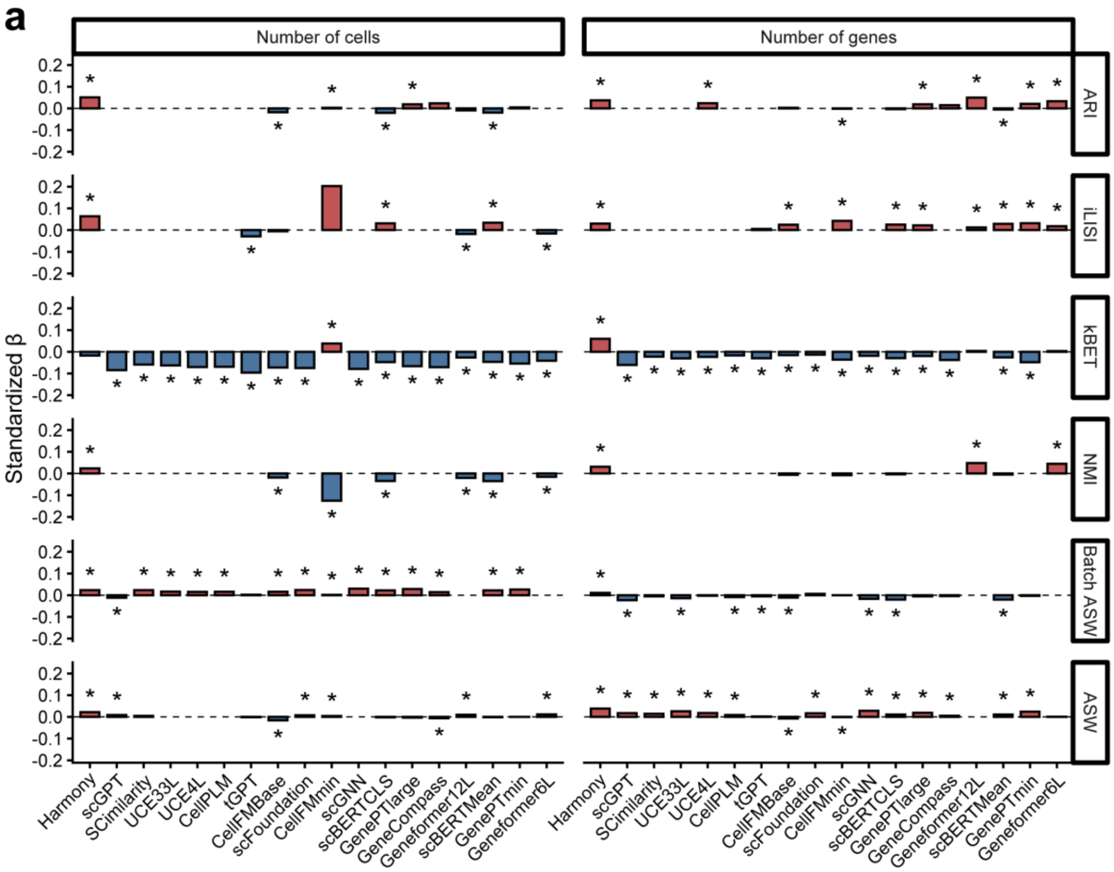


**Supplementary Fig. 33 | Regression-based driver analysis of model performance.** Regularized regression coefficients quantify the associations between dataset features and model metrics. **a,** The impact of the "Number of cells" and "Number of genes" across evaluated models.

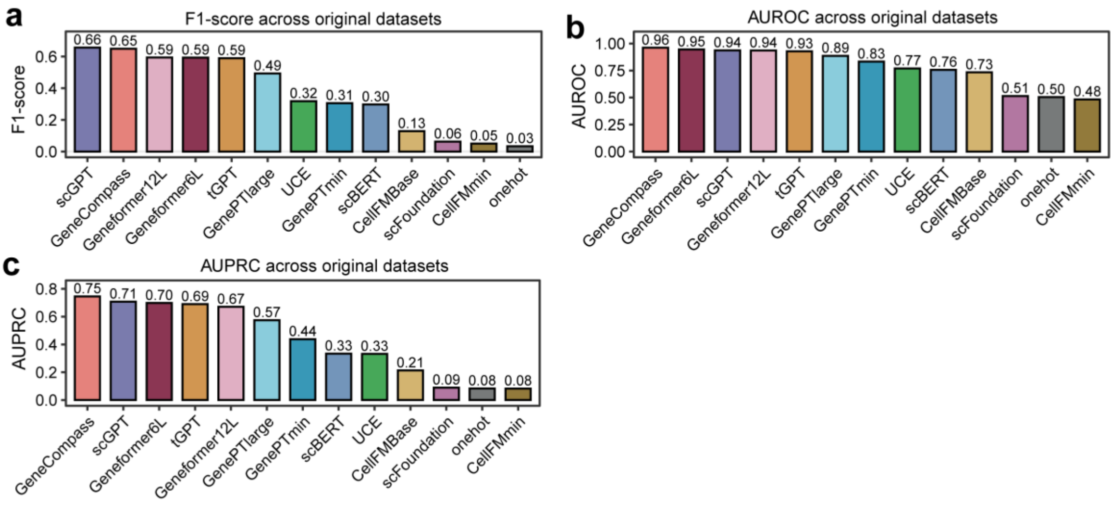


**Supplementary Fig. 34 | Performance comparison of evaluated methods on HPA dataset in gene function prediction.** Bar plots showing performance of 13 methods on HPA datasets measured by F1-score **(a),** AUROC **(b),** AUPRC **(c)** respectively. For each metric, models are ranked along the x-axis in descending order.

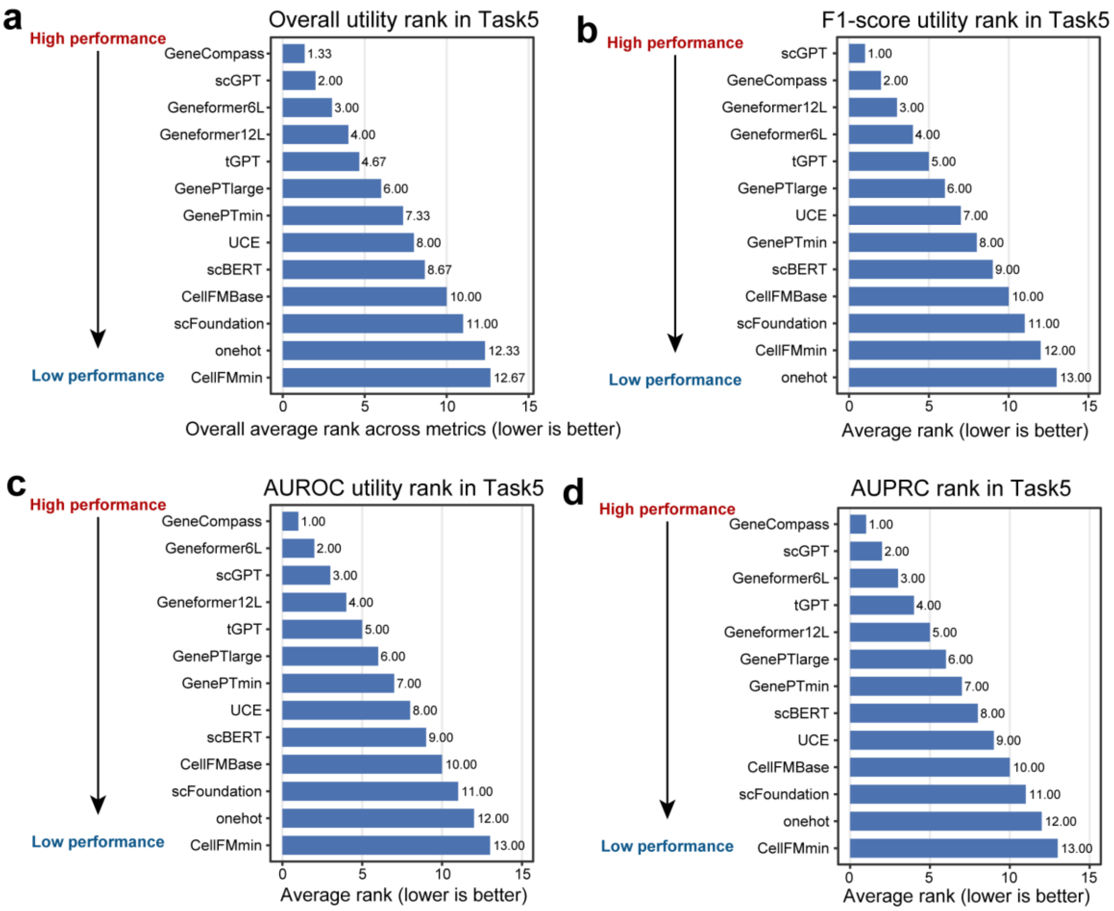


**Supplementary Fig. 35 | Average utility rankings for zero-shot gene function prediction.** Bar plots display the overall aggregated rank across all three metrics **(a)**, alongside the individual average ranks across the 7 datasets for F1-score **(b)**, AUROC **(c)**, and AUPRC **(d)**. Lower numerical rank values indicate superior method performance.

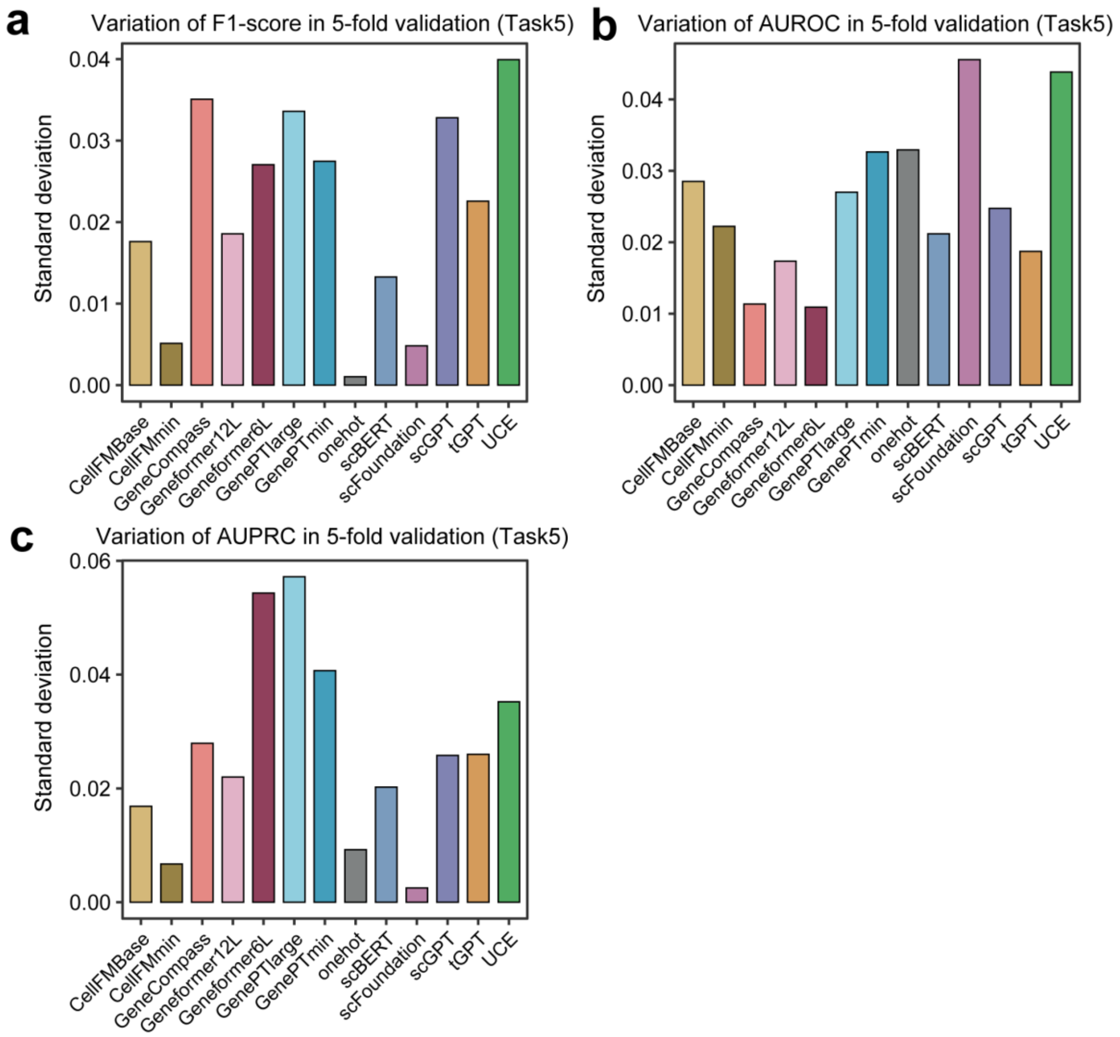


**Supplementary Fig. 36 | Standard deviation of performance metrics across 5-fold cross-validation for gene function prediction.** Barplots illustrate the variance in **(a)** F1-score, **(b)** AUROC, and **(c)** AUPRC for different algorithms. Lower standard deviation values indicate higher model stability across data folds.

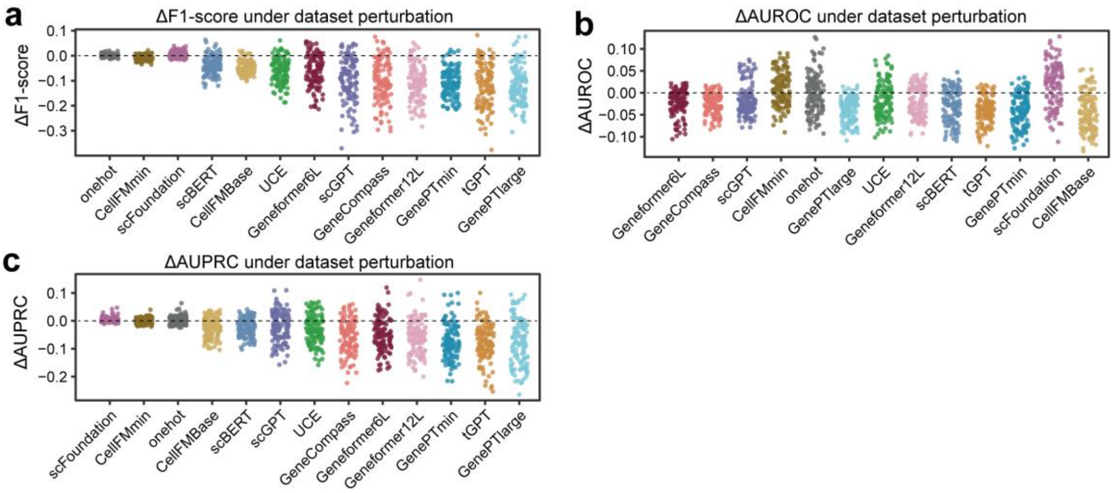


**Supplementary Fig. 37 | Robust of evaluated methods in gene function prediction.** Scatter plots showing performance deviations under perturbation-derived datasets for **(a)** F1-score, **(b)** AUROC, and **(c)** AUPRC respectively. Each point represents one perturbation instance.

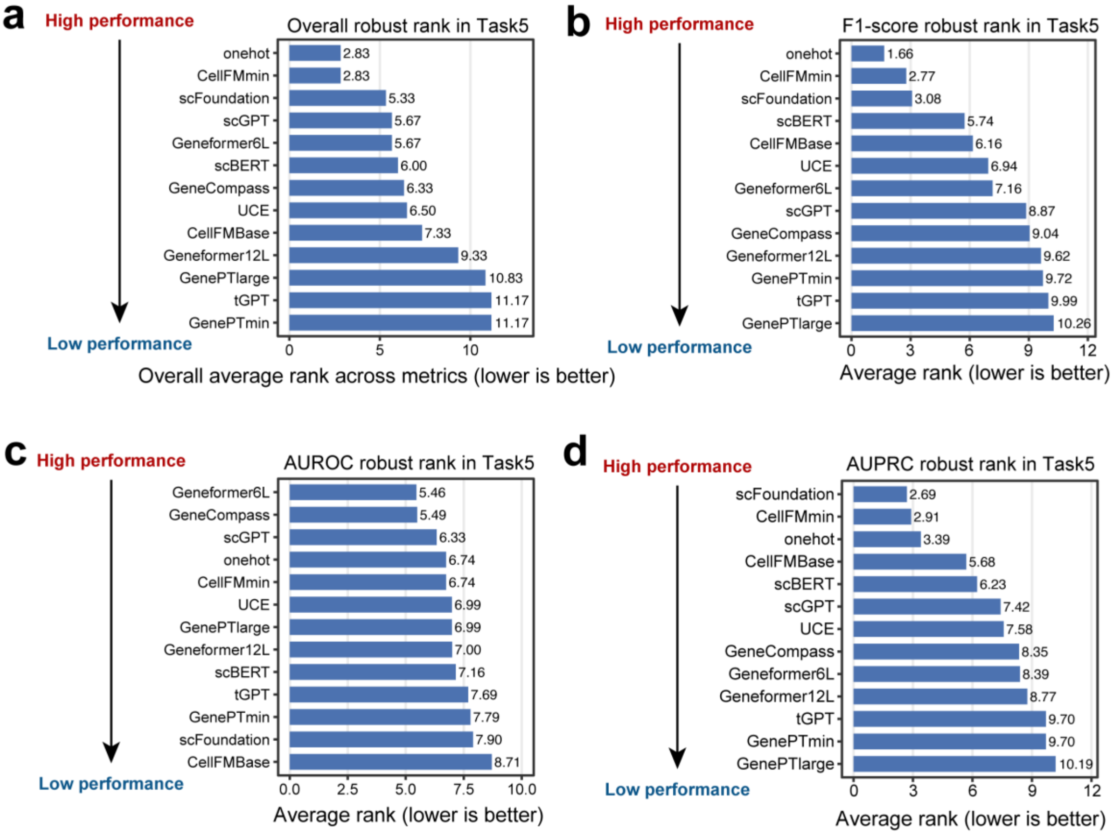


**Supplementary Fig. 38 | Average robustness rankings for zero-shot gene function prediction.** Bar plots display the overall aggregated rank across all three metrics **(a)**, alongside the individual average ranks across the 120 datasets for F1-score **(b)**, AUROC **(c)**, and AUPRC **(d)**. Lower numerical rank values indicate superior robustness.

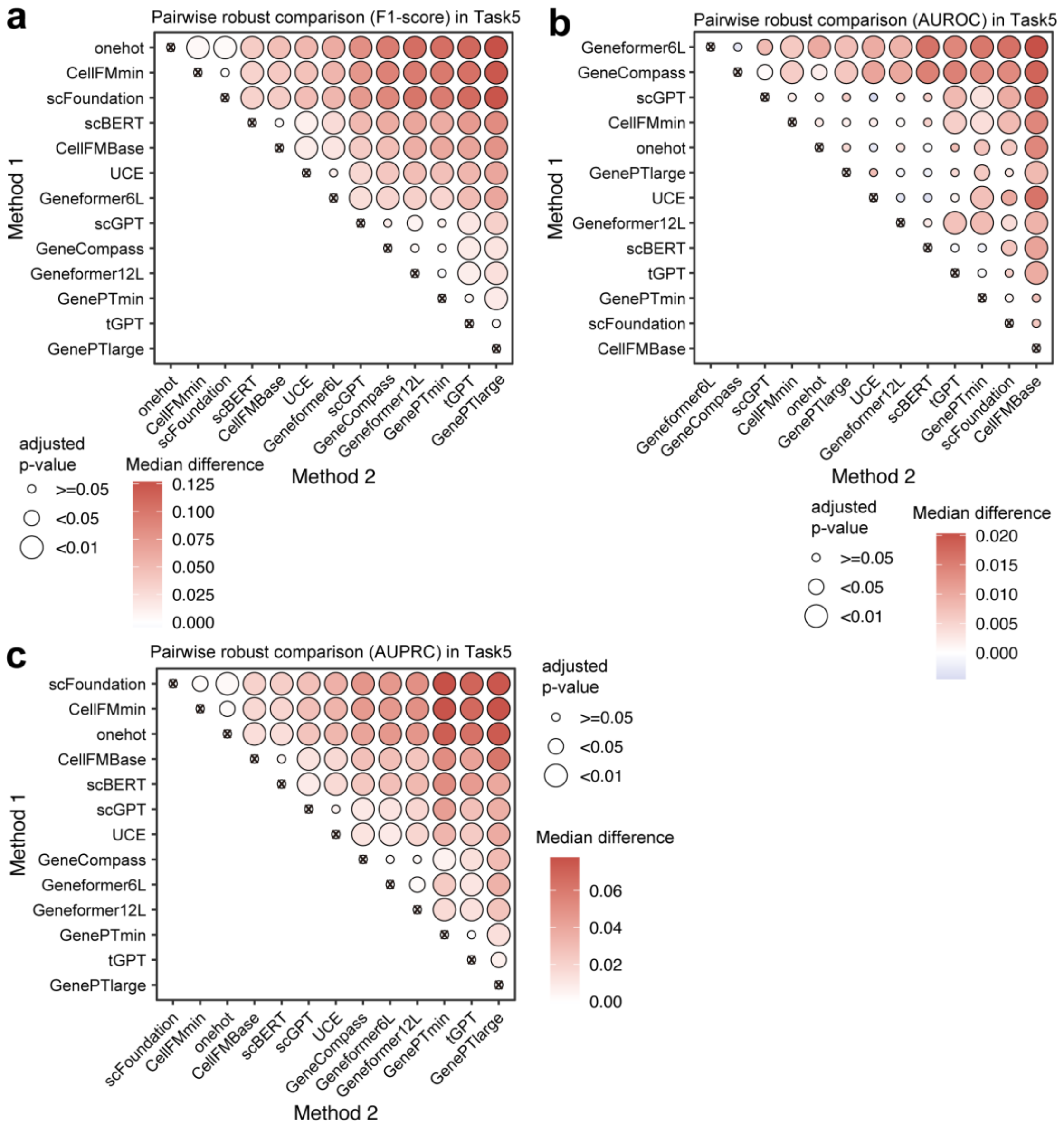


**Supplementary Fig. 39 | Pairwise statistical robustness comparisons of zero-shot gene function prediction. a-c,** Bubble plots visualize the results of one-sided Wilcoxon tests comparing method robustness across F1-score **(a)**, AUROC **(b)**, and AUPRC **(c)** metrics, with algorithms ordered along the axes by their independent average ranks. Bubble color intensity represents the median robustness difference between the Method 2 and Method 1 algorithms, while bubble size indicates the adjusted p-value.

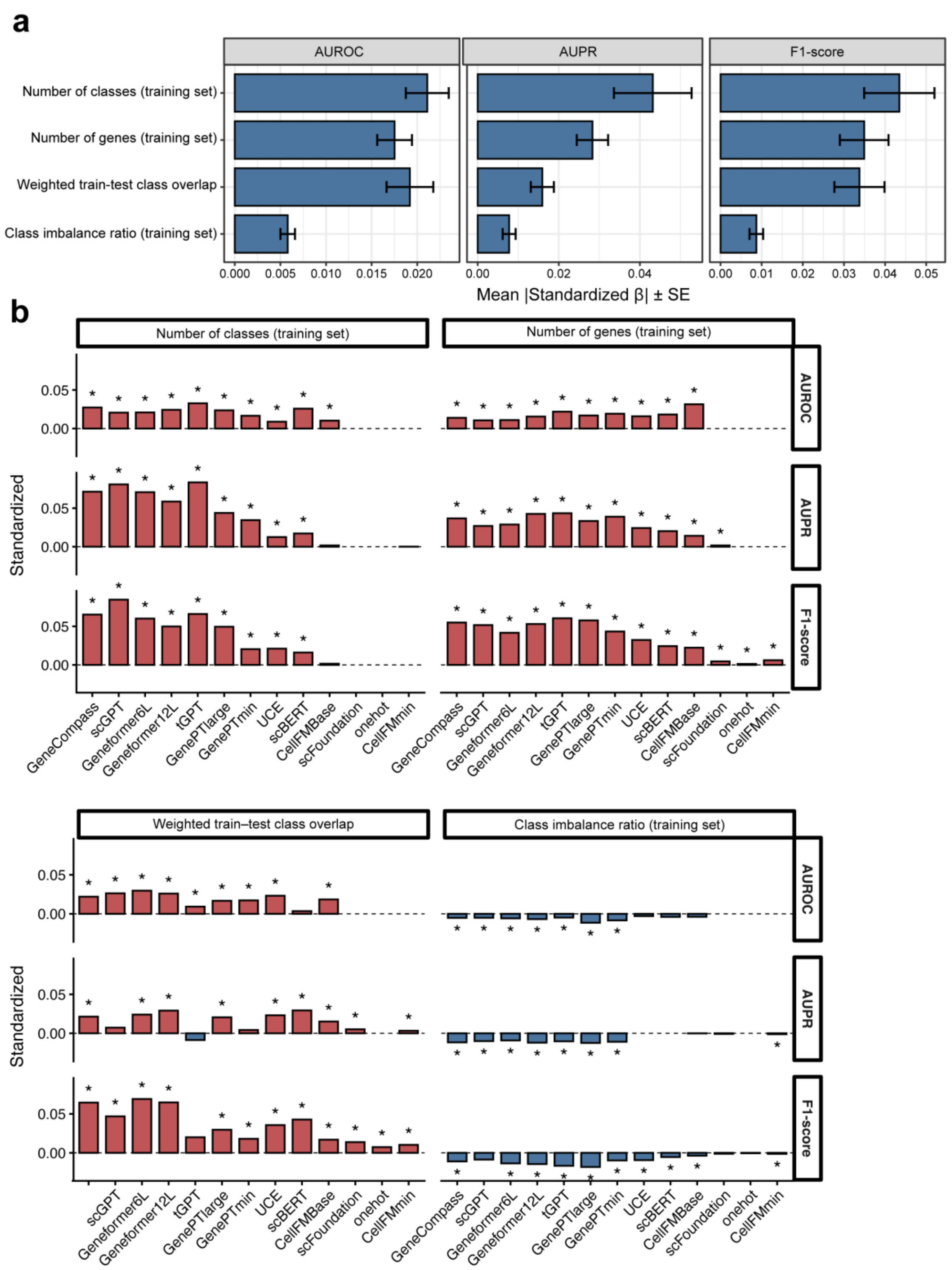


**Supplementary Fig. 40 | Regression-based driver analysis of model performance.** **a,** Bar plots summarize the aggregated regression coefficients for dataset features across all evaluated methods, with error bars indicating 95% confidence intervals. **b,** Regularized regression coefficients quantify the associations between dataset features and model metrics. The magnitude and direction of these coefficients reveal how specific structural data features systematically drive algorithmic performance across the evaluated methods.

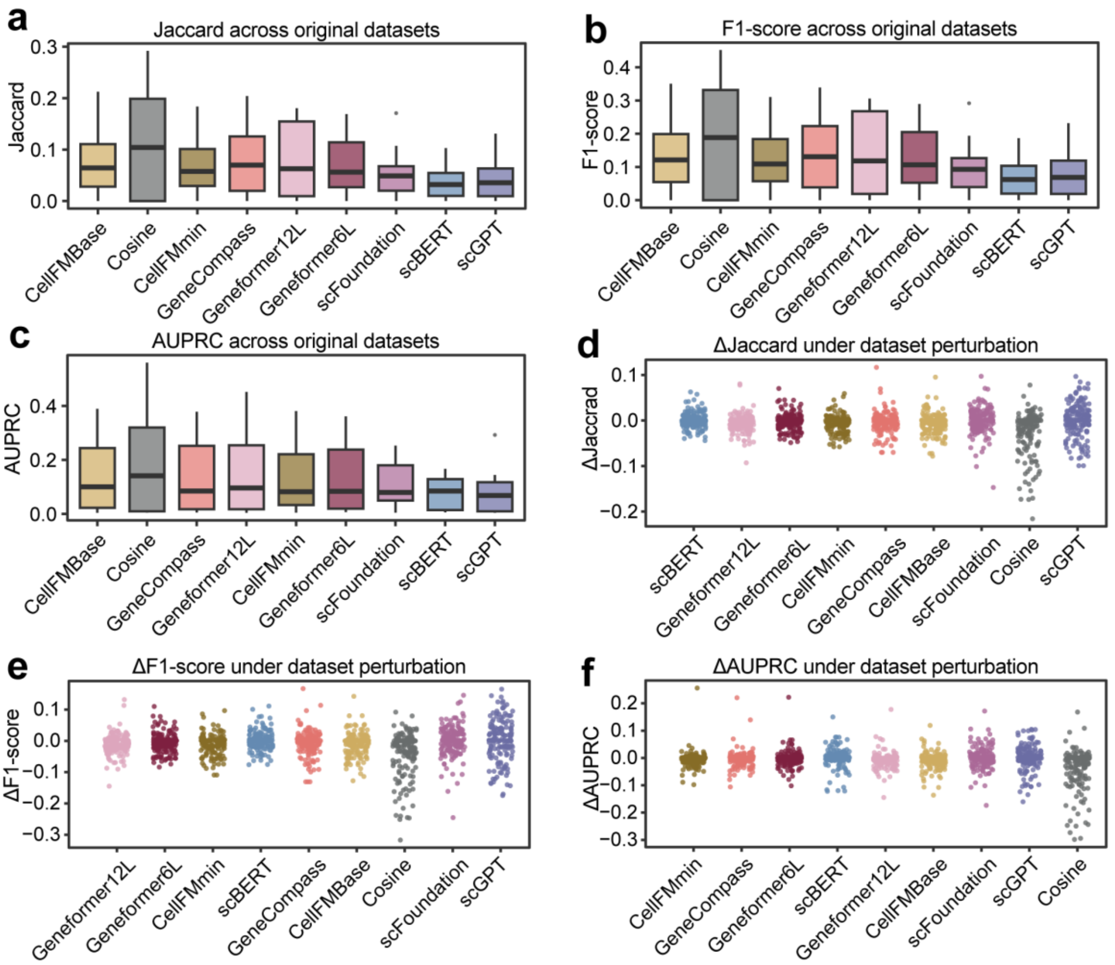


**Supplementary Fig. 41 | Utility and robustness evaluation on the gene regulatory network inference. a-c,** Boxplots illustrating the predictive performance of evaluated methods across original datasets, measured by (a) Jaccard, (b) F1-score, and (c) AUPRC. **d-f,** Scatter plots displaying the robustness of the methods under dataset perturbation. The y-axis represents the change in performance for (d) Jaccard index, (e) F1-score, and (f) AUPRC after introducing perturbations. Methods with performance changes distributed closer to zero exhibit higher robustness against dataset variations.

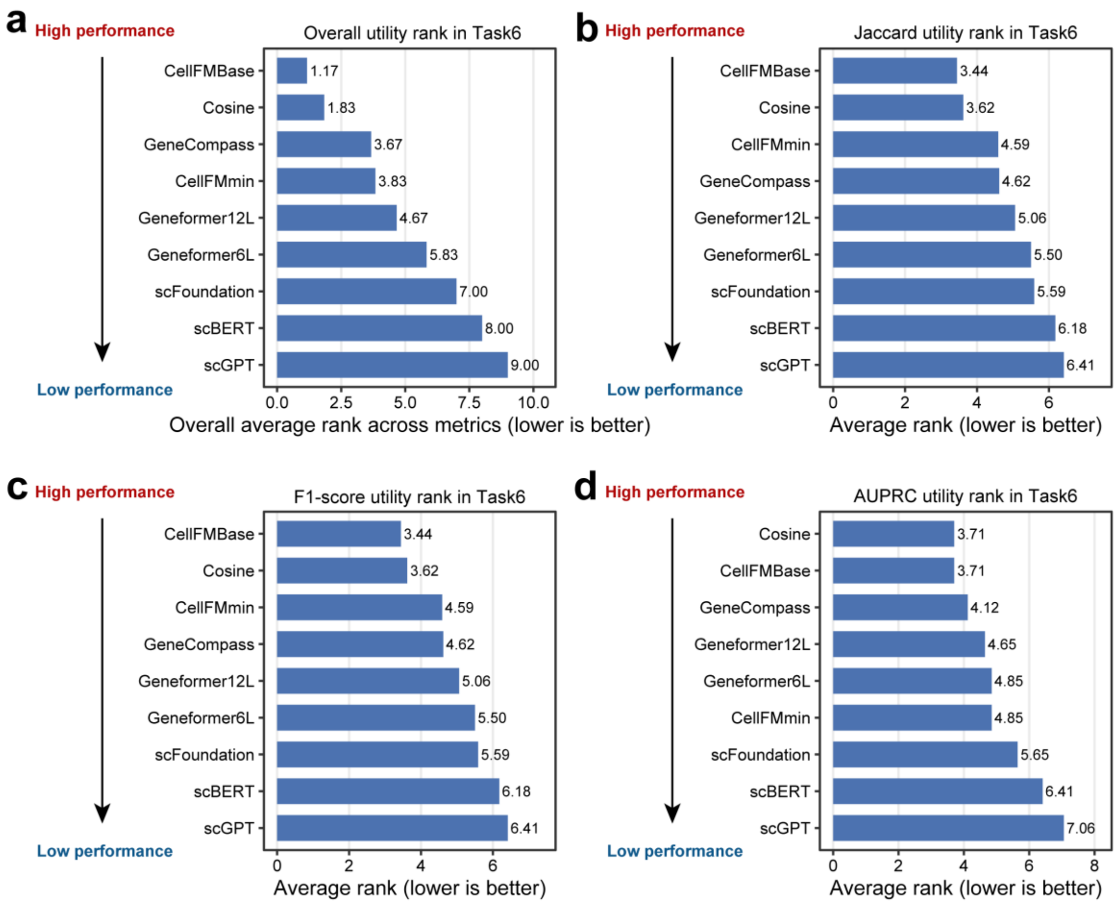


**Supplementary Fig. 42 | Average utility rankings for zero-shot gene regulatory network inference.** Bar plots display the overall aggregated rank across all three metrics **(a)**, alongside the individual average ranks across the 17 datasets for Jaccard **(b)**, F1-score **(c)**, and AUPRC **(d)**. Lower numerical rank values indicate superior method performance.

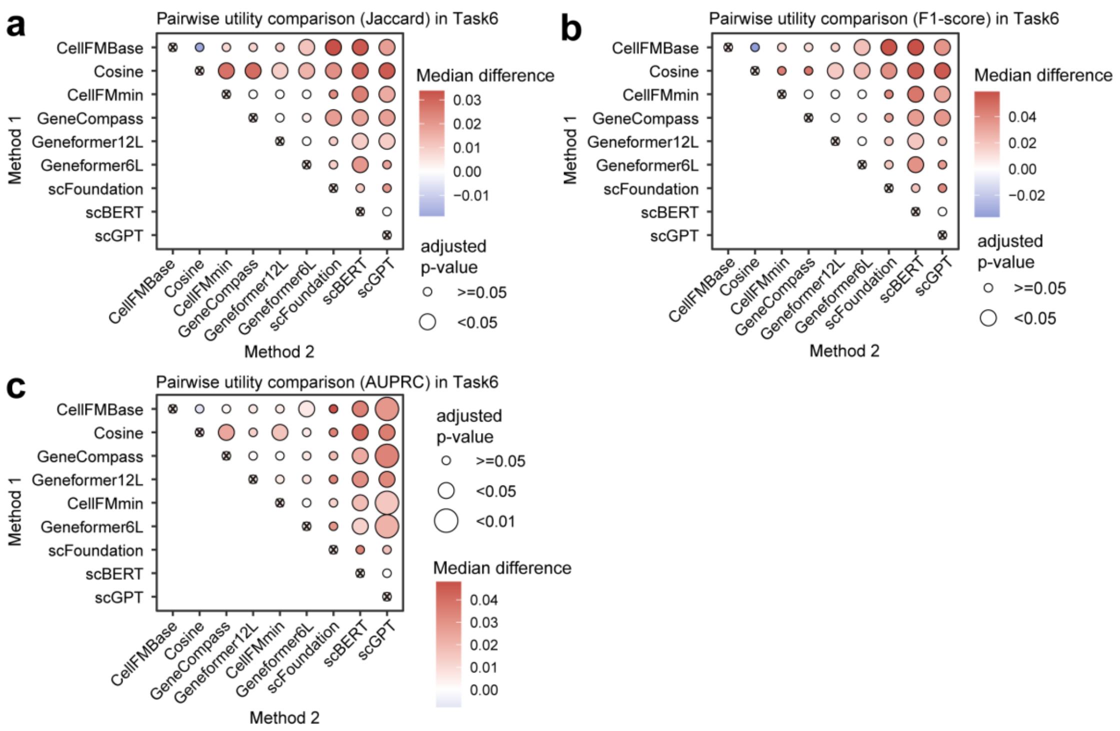


**Supplementary Fig. 43 | Pairwise statistical comparisons of zero-shot utility in gene regulatory network inference. a-c,** Bubble plots visualize the results of one-sided Wilcoxon tests comparing method performance across Jaccard **(a)**, F1-score **(b)**, and AUPRC **(c)** metrics, with algorithms ordered along the axes by their independent average ranks. Bubble color intensity represents the median performance difference between the Method 1 and Method 2 algorithms, while bubble size indicates the adjusted p-value.

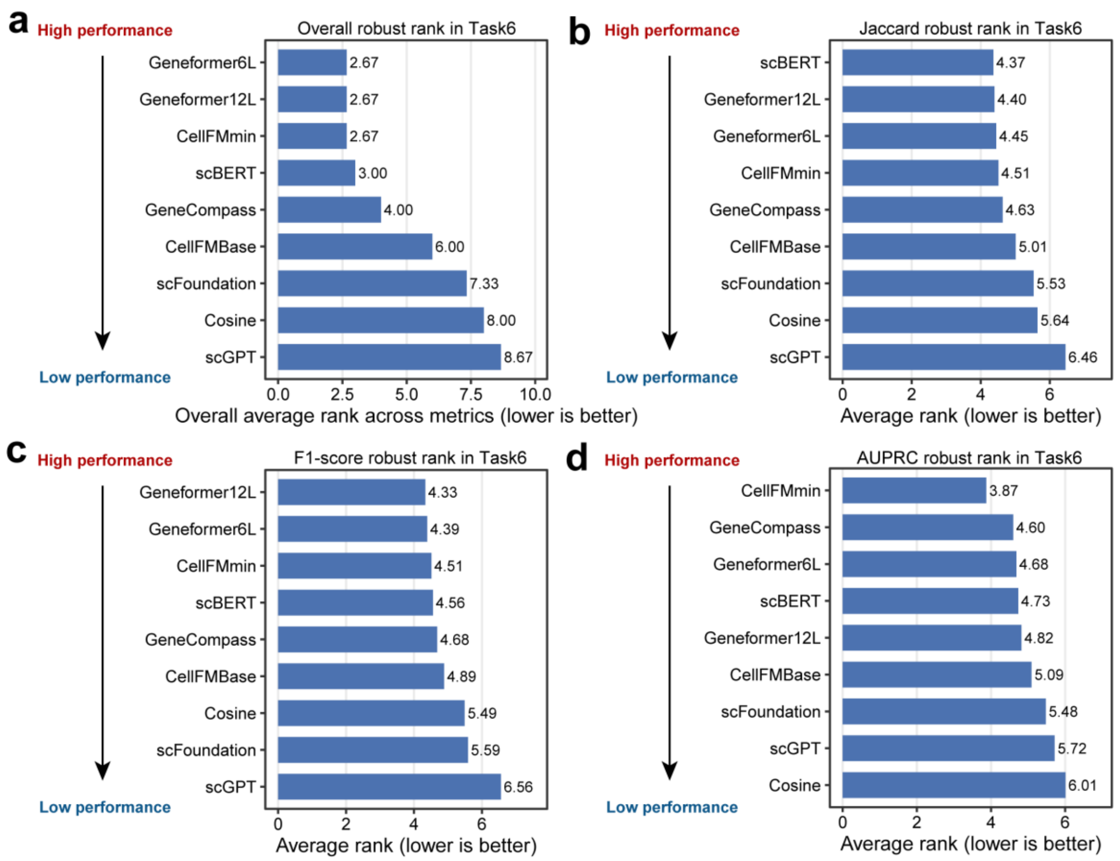


**Supplementary Fig. 44 | Average robustness rankings for zero-shot gene regulatory network inference.** Bar plots display the overall aggregated rank across all three metrics **(a)**, alongside the individual average ranks across the 167 datasets for Jaccard **(b)**, F1-score **(c)**, and AUPRC **(d)**. Lower numerical rank values indicate superior robustness.

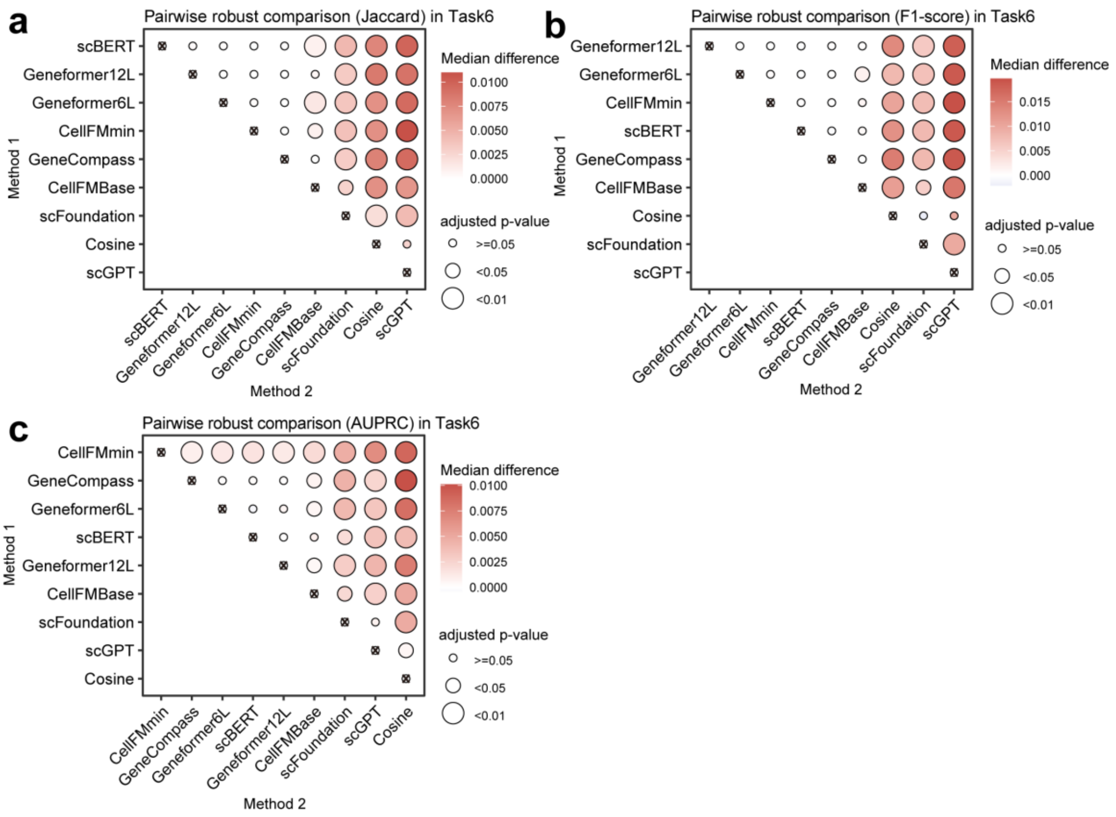


**Supplementary Fig. 45 | Pairwise statistical robustness comparisons of zero-shot gene regulatory network inference. a-c,** Bubble plots visualize the results of one-sided Wilcoxon tests comparing method robustness across Jaccard **(a)**, F1-score **(b)**, and AUPRC **(c)** metrics, with algorithms ordered along the axes by their independent average ranks. Bubble color intensity represents the median robustness difference between the Method 2 and Method 1 algorithms, while bubble size indicates the adjusted p-value.

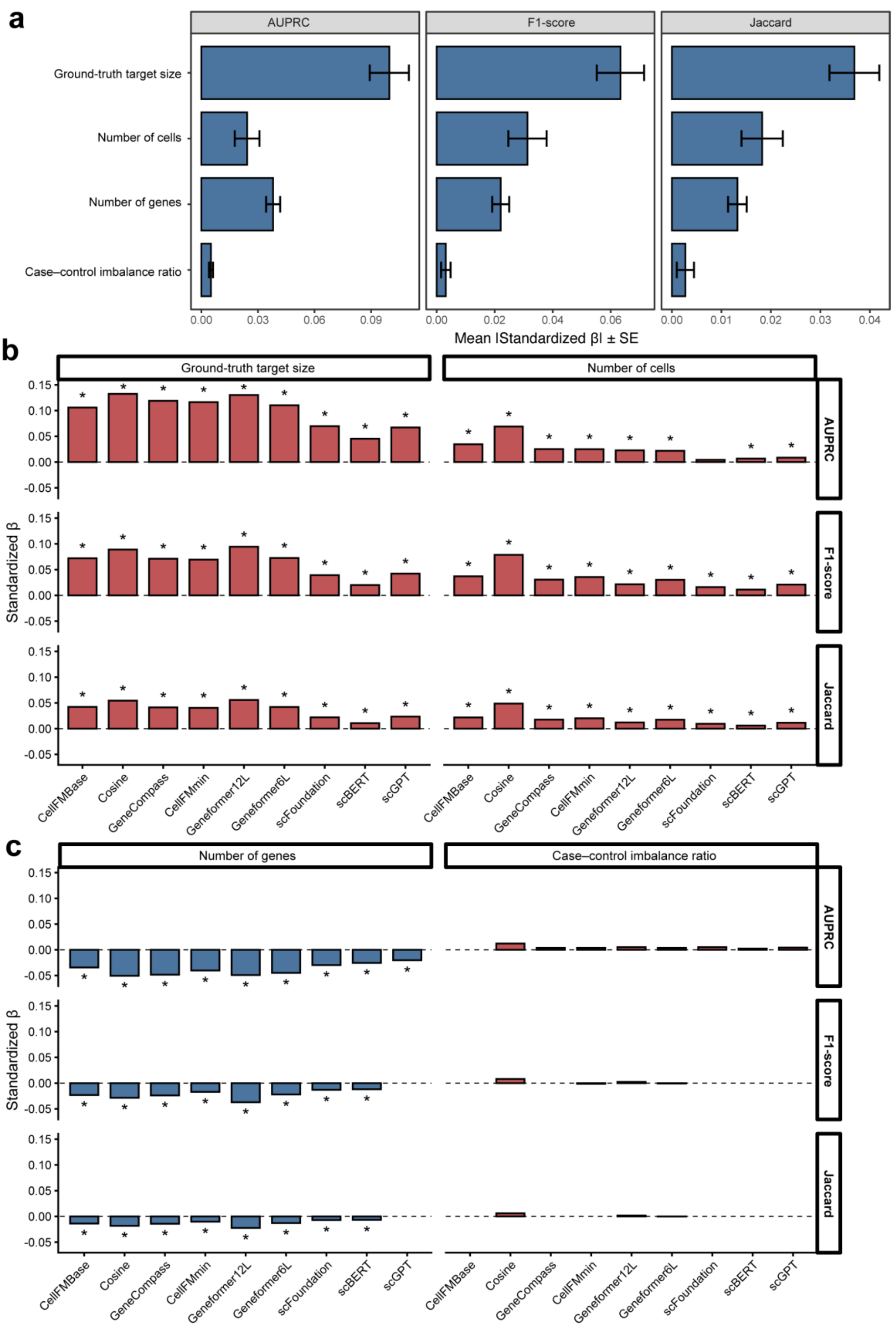


**Supplementary Fig. 46 | Regression-based driver analysis of model performance.** **a,** Bar plots summarize the aggregated regression coefficients for dataset features across all evaluated methods, with error bars indicating 95% confidence intervals. **b,**

Regularized regression coefficients quantify the associations between dataset features and model metrics. The magnitude and direction of these coefficients reveal how specific structural data features systematically drive algorithmic performance across the evaluated methods.
